# Global evolutionary and spatiotemporal transmission dynamics of human respiroviruses and orthorubulaviruses

**DOI:** 10.64898/2026.11.26.769081

**Authors:** Gavin J.D. Smith, Peter Cronin, Rong Zhang, Bo Wang, Dongqiang Cheng, Yan Zhuang, Foong Ying Wong, Jordan Ma, Stephanie S.M. Goay, Chiang Yong Shi, Giselle G.K. Ng, Yaw Shin Ooi, Ashley St. John, De-Yun Wang, Tamara Garcia-Salum, Jean-Michel Héraud, Martin Schutten, Theo P. Sloots, Florette K. Treurnicht, Ana Maria Contreras, Marcela Ferrés, Julian D. Druce, Norosoa Harline Razanajatovo, Benjamin J. Cowling, Dominic E. Dwyer, Cui Lin, Cheryl Cohen, David W. Smith, Rafael A. Medina, Yvonne C.F. Su

## Abstract

Human parainfluenza viruses (HPIVs) cause recurrent epidemics and are a major cause of hospitalization worldwide. However, the evolutionary and transmission dynamics driving HPIV persistence remain unclear. We analysed 1,161 archived samples representing four HPIV types from seven countries using genomic sequencing and Bayesian phylodynamic inference. We show that *Respirovirus* (HRV-1 and HRV-3) displayed greater fluctuations in genetic diversity whereas *Orthorubulavirus* (HOV-2 and HOV-4) showed more gradual changes. Despite comparable evolutionary rates, the four HPIV types exhibited contrasting lineage dynamics. HRV-1 showed prolonged lineage persistence, HRV-3 underwent greater lineage turnover, and HOV-2 and HOV-4 maintained fewer co-circulating lineages. Positive selection was more pronounced in respiroviruses, particularly in the phosphoprotein gene, while positively selected sites overlapped predicted hemagglutinin-neuraminidase B-cell epitopes. Phylogeographic analyses revealed repeated interregional introductions, bidirectional transmission, and regional persistence exceeding a decade for some lineages. These findings identify distinct evolutionary and transmission dynamics underlying HPIV persistence and inform the development of broadly protective vaccines and therapeutics.

## Introduction

Human parainfluenza viruses (HPIVs) are leading causes of acute lower respiratory infections (ALRIs) worldwide and impose a substantial public health burden. They are associated with an estimated 13% of ALRI cases, 4–14% of ALRI-related hospitalizations, and 4% of ALRI-related deaths globally in children younger than five years (1, 2). Infection affects all age groups, particularly infants, young children, adults over 65 years, and immunocompromised individuals (2). Waning HPIV-specific immunity permits lifelong reinfection, and common coinfections with rhinoviruses/enteroviruses, seasonal coronaviruses, and other respiratory viruses further complicate disease outcomes (3, 4).

First recognized in the late 1950s, HPIVs were isolated as several ‘influenza-like’ viruses from children with croup and lower respiratory tract infections. Despite causing influenza-like symptoms, they are biologically and taxonomically distinct from influenza viruses (5). By 1960, four antigenically distinct types (HPIV-1 to HPIV-4) had been defined. Infections are generally mild and resemble other respiratory viral illnesses, but disease associations vary by type (2, 5). HPIV-1 and HPIV-2 are more often associated with upper respiratory tract disease, including croup, whereas HPIV-3 is a major cause of bronchiolitis and pneumonia (2). HPIV-4 comprises lineages 4A and 4B and can be associated with upper and lower respiratory tract infections (6). Across age groups, HPIVs can exacerbate asthma and chronic obstructive pulmonary disease. In immunocompromised patients, particularly hematopoietic stem cell transplant recipients, infection may progress to severe or prolonged lower respiratory tract disease and respiratory failure (2, 7).

HPIVs show type-specific seasonality in temperate regions such as the United States. HPIV-1 peaks in autumn with biennial surges; HPIV-2 also circulates in autumn and may alternate with HPIV-1; and HPIV-3 peaks from spring to early summer (2, 8). Although HPIV-4 seasonality is less well defined because surveillance data are limited. PCR-based surveillance suggests recurrent activity from late summer through winter and intermittent year-round detection (8, 9). Climatic factors may also influence HPIV seasonality. Higher temperature has been associated with increased HPIV-3 incidence, whereas HPIV-4 incidence has been negatively associated with temperature and positively associated with rainfall (9). HPIVs also cause outbreaks in community and healthcare settings. HPIV-1 has been associated with childcare outbreaks, including a kindergarten outbreak in Beijing in 2018 (10). HPIV-3 is frequently linked to healthcare-associated transmission, including neonatal intensive care unit (NICU) outbreaks in the United States in 2019 and Saudi Arabia in 2022–2023 (11, 12). HPIV-4 has been associated with institutional and nosocomial outbreaks (13), whereas HPIV-2 has been reported less frequently in such outbreaks.

HPIVs are members of the family *Paramyxoviridae*, which comprises four subfamilies and 17 genera, including the phylogenetically distinct *Respirovirus* and *Orthorubulavirus*. The genus *Respirovirus* contains viruses of diverse host origins, including rodents, swine, goats, and cattle and is closely related to *Aquaparamyxovirus* and *Ferlavirus*, which infect fish and reptiles, respectively (Fig. 1A). The *Orthorubulavirus* genus mainly contains viruses of bat and non-human primate origin and is sister to the predominantly bat origin *Pararubulavirus*. HPIV-1 and HPIV-3 were historically classified in *Respirovirus*, whereas HPIV-2 and HPIV-4 were assigned to *Rubulavirus* (5). These enveloped, nonsegmented, negative-sense RNA viruses have ∼16 kb genomes encoding six major proteins involved in entry, replication, and assembly (Fig. 1B). HPIV-1 and HPIV-3 have similar genome lengths (∼15.6 kb), whereas HPIV-2 and HPIV-4 have longer intergenic regions. The hemagglutinin-neuraminidase glycoprotein (HN) mediates entry by binding sialic acid receptors and facilitates virion release through neuraminidase activity. The fusion protein (F) drives membrane fusion and cell-to-cell spread, often producing syncytia (5). The nucleocapsid protein (N) encapsidates viral RNA and provides the template for transcription and replication; the phosphoprotein (P) and large polymerase (L) form the core replication machinery; and the matrix protein (M) coordinates virion assembly at the plasma membrane (5). In HPIV-2 and HPIV-4, an additional non-structural accessory V protein modulates the innate immune response (5).

**Figure 1.**
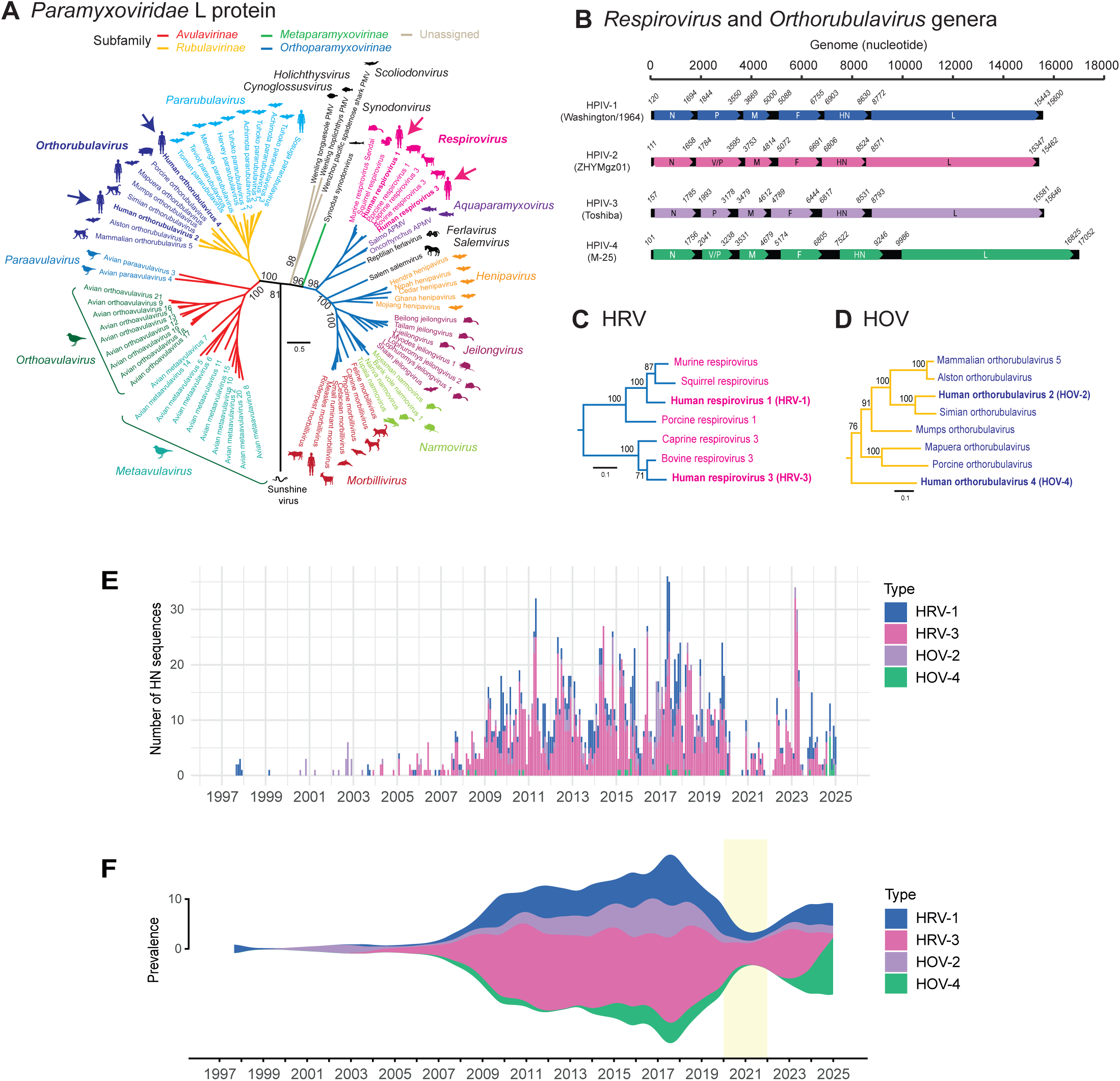
Classification and global epidemiology of human parainfluenza viruses. (*A*) Taxonomic relationships of the genera *Respirovirus* and *Orthorubulavirus* within the family *Paramyxoviridae*, inferred from L protein amino acid sequences. Coloured arrows indicate human respirovirus 1 (HRV-1), human respirovirus 3 (HRV-3), human orthorubulavirus 2 (HOV-2) and human orthorubulavirus 4 (HOV-4) types. Coloured branches denote subfamilies, with bootstrap support values indicated on the branches. Host species corresponding to each virus species are indicated. (*B*) Whole-genome organization of *Respirovirus* and *Orthorubulavirus*, showing the gene regions; numbers above the genome schematics indicate nucleotide positions. GenBank accession numbers for the reference strains are as follows: AF457102 for HRV-1, EU326526 for HRV-3, X57559 for HOV-2, and AB543336 for HOV-4. (*C*) Evolutionary relationships of HRV-1, HRV-3 and related viruses. (*D*) Evolutionary relationships of HOV-2, HOV-4 and related viruses. (*E*) Number of available hemagglutinin-neuraminidase (HN) sequences by month for each HPIV type, including sequences generated in this study and retrieved from NCBI GenBank. Thin lines show monthly counts and thick lines show LOESS fits. (*F*) Streamline plot showing the number of available HN sequences by month for each HPIV type. Yellow vertical bar indicates the COVID-19 period during 2020 to 2022.

Evolutionary studies of HPIVs have focused largely on HN, particularly in HPIV-3, and have generally been regional, with studies in Argentina, China, Japan, Russia, the United States, and Viet Nam (3, 14–18). Under current International Committee on Taxonomy of Viruses nomenclature, the viruses are human respirovirus 1 (*Respirovirus laryngotracheitidis*), human orthorubulavirus 2 (*Orthorubulavirus laryngotracheitidis*), human respirovirus 3 (*Respirovirus pneumoniae*), and human orthorubulavirus 4 (*Orthorubulavirus hominis*) (19). We use HPIV to refer to all four viruses collectively. HRV-1 and HRV-3 denote HPIV-1 and HPIV-3, and HOV-2 and HOV-4 denote HPIV-2 and HPIV-4, respectively (Fig. 1C and D). HOV-4 comprises two genetically distinct lineages, HOV-4A and HOV-4B, which diverged after HOV-4 emerged and subsequently co-circulated (6, 20).

Despite more than 70 years of co-circulation, the evolutionary dynamics of HPIV types remain poorly understood, especially for infrequently detected types with limited genomic data. The COVID-19 pandemic caused an unprecedented disruption of respiratory virus transmission, causing widespread reductions in virus circulation followed by the resurgence of multiple respiratory viruses after the relaxation of public health control measures (21–23). Whether this disruption altered HPIV population dynamics, lineage composition, or genetic diversity is unclear. Here we analyzed 1,161 HPIV-positive clinical samples collected from seven locations on multiple continents during 1988–2018, together with all available global genomic sequences spanning 1957–2025. Using next-generation sequencing and phylodynamic and phylogeographic analyses, we examined the evolutionary trajectories, drivers of genetic diversification, and global transmission of all four types. We identify marked differences between *Respirovirus* and *Orthorubulavirus*; HRV-1 and HRV-3 showed greater lineage diversification, temporal fluctuations in genetic diversity, and lineage expansion than HOV-2 and HOV-4 despite comparable evolutionary rates. HRV-1 and HRV-3 also showed stronger signatures of positive selection in HN and P, including sites overlapping predicted B-cell epitopes. We also identified recurrent interregional introductions, prolonged regional persistence, and bidirectional intercontinental transmission and examined changes in HPIV circulation and lineage composition following the COVID-19 pandemic. Together, these analyses characterize the evolutionary and epidemiological processes shaping HPIV persistence and global dissemination.

## Results

### Global patterns of recurrent HPIV epidemics and post-pandemic resurgence

We obtained 1,161 archived HPIV-positive samples collected over a 30-year period (1988–2018) from seven collaborating laboratories in Australia, Chile, Hong Kong, Madagascar, the Netherlands, Singapore, and South Africa (*SI Appendix*, Fig. S1 and Table S1). This geographically and temporally diverse collection enabled comprehensive analyses of HPIV evolution. Using next-generation sequencing, we generated 279 complete genomes across all four HPIV types (HRV-1, HRV-3, HOV-2, and HOV-4), increasing the number of publicly available genomes by nearly 30% and substantially expanding global sequence representation (*SI Appendix*, Fig. S1).

Because early studies focused primarily on the HN gene, the number of available HN sequences increased from 2009 onwards, with only intermittent detection in 2020 (likely reflecting COVID-19-related disruptions) (Fig. 1E). HRV-3 was the most frequently detected type, accounting for 67% of available HN sequences, whereas HRV-1, HOV-2, and HOV-4 each accounted for 2.5%–19.8% despite co-circulating in most years (Fig. 1E; *SI Appendix*, Table S1). HRV-3 showed pronounced spring seasonality, while HRV-1 peaks tended to alternate with those of HRV-3 (Fig. 1E). HOV-2 showed a seasonal pattern similar to HRV-3, whereas HOV-4 seasonality was weaker. Viral activity occurred primarily in winter in Europe but year-round in South and Southeast Asia (*SI Appendix*, Fig. S2). Sequence numbers before 2007 were low, and changes in these counts over time reflect testing and sequencing effort as well as virus circulation, so they do not indicate incidence or prevalence.

The number of available HPIV sequences increased from mid-2017 and declined sharply during the COVID-19 pandemic (Fig. 1E), mirroring the decline reported for many other respiratory pathogens, including influenza viruses, respiratory syncytial virus, pneumococcal disease and pertussis, which was likely due to non-pharmaceutical interventions that reduced respiratory pathogen transmission and possibly due to reduced testing for viruses other than SARS-CoV-2. All four HPIV types subsequently re-emerged and re-established co-circulation. HRV-3 sequence numbers increased markedly in early 2023, whereas HRV-1 and HOV-4 detections increased substantially from early to mid-2024, coinciding with comparatively fewer HRV-3 sequences (Fig. 1F).

### HRV-1 and HRV-3 exhibit distinct evolutionary histories associated with lineage emergence and persistence

HRV-1 and HRV-3 evolved at comparable genome-wide substitution rates (7.97 × 10^−4^ and 6.73 × 10^−4^ substitutions/site/year, respectively (*SI Appendix*, Fig. S3 and Table S2), but their evolutionary histories and clade dynamics differed markedly. HRV-1 comprised two major clades that diverged in the early 1990s (mean TMRCA, June 1994; Fig. 2A; *SI Appendix*, Table S2). Seven nonsynonymous amino acid substitutions in the P, F, HN, and L proteins were associated with this divergence, and both clades have circulated globally for more than three decades (Fig. 2A; *SI Appendix*, Table S3). Clade 1 subsequently split into clades 1.1 and 1.2. Subsequent diversification was largely confined to clade 1.2 and accompanied by sequential amino acid substitutions, particularly in P (e.g., S274L, P441S, R92G, S442N, V271I, D11N, and D256E) and HN (e.g., V573I, I60M, Q132R, and A40T; Fig. 2A). Clade 1.2 underwent further lineage expansion and was associated with regional outbreaks in Europe (2022–2023) and North America (2024–2025) (shaded blue box in Fig. 2A; *SI Appendix,* Fig. S4). In contrast, clade 1.1 accumulated fewer substitutions and was rarely detected after 2014. Clade 2 split into subclades 2.1 and 2.2, with substantial expansion mainly in clade 2.2 associated with earlier outbreaks in Chile (2017; red triangle in Fig. 2A) and Africa (2013–2014).

**Figure 2.**
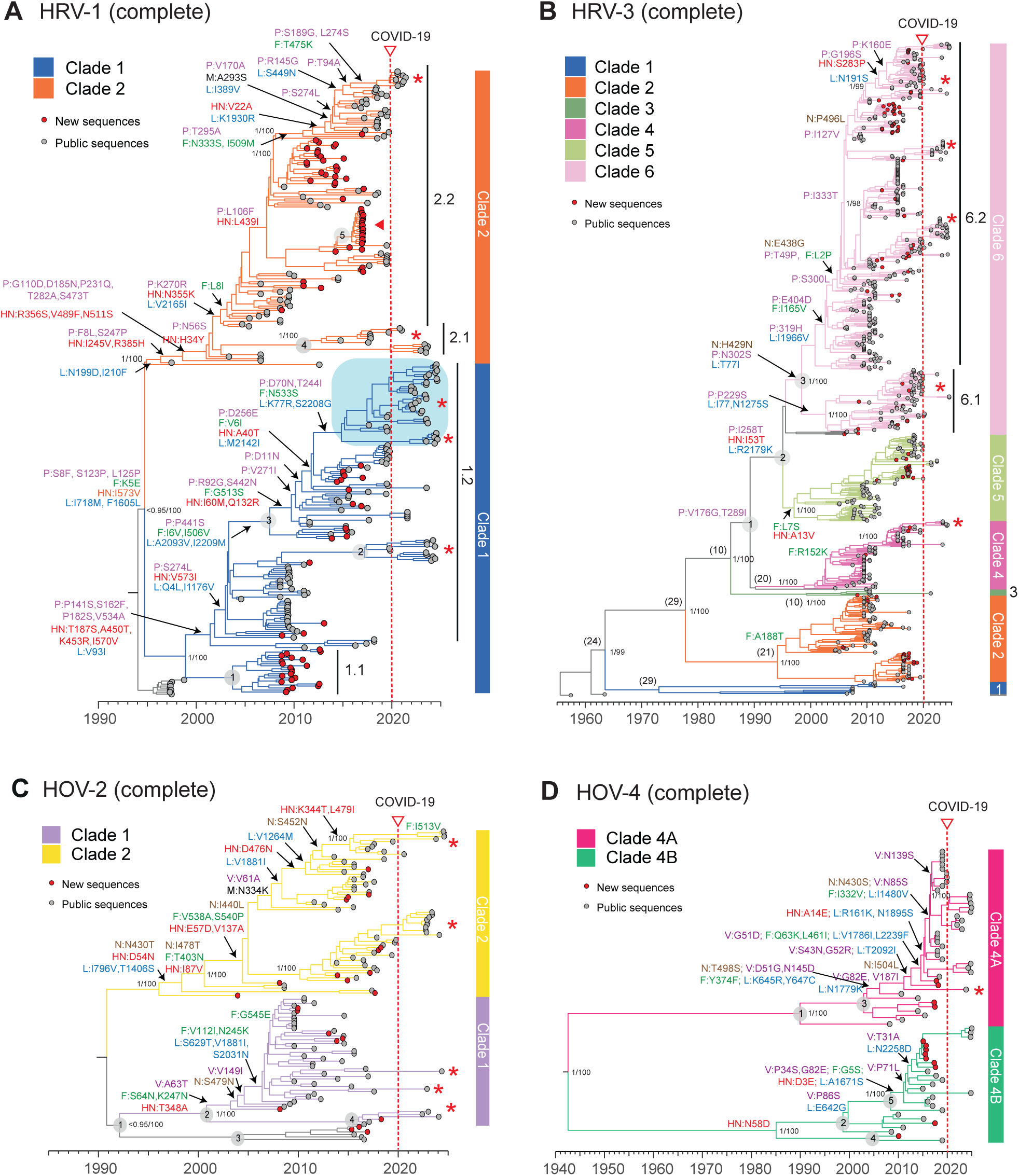
Evolutionary dynamics of human parainfluenza viruses before and after COVID-19 pandemic. Evolutionary relationships and divergence times of (*A*) HRV-1, (*B*) HRV-3, (*C*) HOV-2, and (*D*) HOV-4. For each type, maximum clade credibility (MCC) tree was reconstructed based on full-genome sequences. Coloured branches denote clades, and sub-lineages are indicated. Red tip circles denote sequences generated from this study, whereas grey tip circles represent available sequences from NCBI GenBank. Vertical red dotted line marks the onset of 2020 COVID-19 pandemic. Red asterisks indicate viral lineages that re-emerged in the post-pandemic era. Amino acid substitutions were mapped at major nodes. Substitutions for a sub-lineage within HRV-1 clade 1.2 cluster, highlighted by a blue shaded box, are displayed in Appendix Figure S4. Bayesian posterior probability and bootstrap support values are indicated at key nodes. Abbreviations for genes: N, nucleocapsid protein; P, phosphoprotein; M, matrix protein; F, fusion protein; HN, hemagglutinin-neuraminidase protein and L, large polymerase protein.

HRV-3 displayed a more ladder-like phylogeny characterized by successive lineage replacement and the early accumulation of numerous amino acid substitutions. This pattern was consistent with adaptation to humans after zoonotic transmission from caprine or bovine ancestors (Fig. 1A and C). Whole-genome evolutionary analysis identified six major clades (clades 1–6; Fig. 2B) based on predefined criteria of >0.7% mean pairwise genetic distance, >1 amino acid substitution across the genome, and >90% bootstrap support, providing a more robust classification than the conventional HN-based system. The mean whole-genome TMRCA of HRV-3 was July 1955 (SI Appendix, Table S2).

HRV-3 clade 1 emerged by 1973 and comprised viruses from the United States and Australia, with 29 amino acid substitutions and relatively long branch lengths that likely reflect limited early sampling (Table 1). Since the 1990s, five additional clades (clades 2–6) have emerged and co-circulated globally (Fig. 2B). Substitutions along the HRV-3 phylogenetic trunk occurred mainly in P and L, with fewer changes in HN (e.g., I53T). A pronounced diversification period occurred from the late 2000s to the early 2010s, coinciding with the peak of the global influenza A(H1N1)pdm09 pandemic (24) suggesting that changes in population immunity or contact patterns may have influenced HRV-3 diversification, although this overlap may be coincidental.

**Table 1.** Evolutionary rates (nucleotide substitutions/site/year) and estimated time to most recent common ancestor (TMRCA) of human respiroviruses and human orthorubulaviruses based on the full genome and clades.

| Type | Clade | TMRCA dates | 95% HPD intervals |
|---|---|---|---|
| HRV-1 | Clade 1 | 1998.46 (1998-06-17) | 1995.68 – 2001.05 (1995-09-06 – 2001-01-17) |
|  | Clade 2 | 1995.16 (1995-02-28) | 1993.30 – 1996.99 (1993-04-20 – 1996-12-26) |
| HRV-3 | Clade 1 | 1973.08 (1973-02-01) | 1967.27 – 1979.96 (1967-04-08 – 1979-12-16) |
|  | Clade 2 | 1994.08 (1994-02-01) | 1990.11 – 1997.45 (1990-02-09 – 1997-06-13) |
|  | Clade 3 | 1999.31 (1999-04-23) | 1994.02 – 2003.64 (1994-01-10 – 2003-08-21) |
|  | Clade 4 | 1990.22 (1990-03-21) | 1985.70 – 1994.11 (1985-09-12 – 1994-02-09) |
|  | Clade 5 | 1997.05 (1997-01-21) | 1994.91 – 1999.08 (1994-11-28 – 1999-02-01) |
|  | Clade 6 | 1995.54 (1995-07-16) | 1993.21 – 1997.71 (1993-03-17 – 1997-09-16) |
| HOV-2 | Clade 1 | 1987.38 (1987-05-17) | 1984.31 – 1992.26 (1984-04-23 – 1992-04-03) |
|  | Clade 2 | 1988.76 (1988-10-03) | 1986.48 – 1992.34 (1986-06-23 – 1992-05-05) |
| HOV-4 | Clade 4A | 2001.54 (2001-07-15) | 1995.12 – 2006.46 (1995-02-13 – 2006-06-17) |
|  | Clade 4B | 1993.57 (1993-07-27) | 1985.32 – 2000.56 (1985-04-27 – 2000-07-24) |

During the late 2010s, clades 5 and 6 underwent substantial lineage expansion, whereas most lineages in clades 1–4 did not persist. After the COVID-19 pandemic, multiple lineages in clades 4–6 persisted, in contrast to HRV-1 lineages, which showed longer-term persistence across successive decades, particularly within clade 1(Fig. 2A).

### Contrasting evolutionary trajectories and lineage dynamics of HOV-2 and −4

HOV-2 and HOV-4 were detected less frequently and evolved at similar rates (6.49 × 10^−4^ and 7.00 × 10^−4^ substitutions/site/year, respectively; *SI Appendix*, Table S2). Whole-genome phylogenies resolved each type into two major clades, with clade-specific substitutions accumulating along the trunk (Fig. 2C and D). HOV-2 clade 2 acquired more sequential substitutions than clade 1, particularly in HN (e.g. D54N, I87V, E57D, and V137A), resulting in a more ladder-like phylogeny (Fig. 2C). Similarly, HOV-4 clade 4A accumulated sequential amino acid substitutions, particularly in the V and L proteins (Fig. 2D).

Nevertheless, both clades of each virus continued to co-circulate after 2020. TMRCA estimates based on whole genomes and HN genes differed for HRV-1 (June 1994 and October 1969, respectively) and HOV-4 (July 1942 and March 1989) but were similar for HRV-3 (July 1955 and March 1957) and HOV-2 (March 1991 and September 1982) (SI Appendix, Table S2). The estimated genome-wide evolutionary rates of the four HPIV types were broadly consistent with previous estimates (3.59–8.09 × 10^−4^ substitutions/site/year) (3, 15, 20). Overall, our TMRCA estimates were more recent than previous estimates (3, 15, 20). For example, our genome-based TMRCA estimate for HRV-3 of July 1955 (95% HPD 1954– 1957) was more recent than previous estimates of approximately 1905 and 1938 (3, 25), possibly because our datasets included more genomes sampled over a longer period.

The HOV-2 HN phylogeny broadly mirrored the whole-genome tree. Clades 1 and 2 were estimated to have emerged at similar times – May 1987 and October 1988, respectively (*SI Appendix*, Fig. S5, Table S1). Sequences generated in this study revealed greater geographic mixing and repeated transmissions. By contrast, HOV-4 HN showed greater geographic structuring, although this may reflect sparse sampling and low detection frequency rather than restricted viral dispersal (*SI Appendix*, Fig. S5).

### Differential population and transmission dynamics shape HPIV epidemics

To characterize HPIV population dynamics, we used Bayesian skyride analyses to estimate temporal variation in relative genetic diversity. HRV-1 and HRV-3 showed broadly similar fluctuations, with clearer seasonal oscillations after 2005 as sequence sampling increased (Fig. 3A). The occurrence of peak diversity differed between types: HRV-1 peaked in July 2009 and May 2017, whereas HRV-3 peaked in December 2008 and March 2017 (Fig. 3A). HOV-2 showed weaker seasonal variation but maintained relatively high genetic diversity from 2003 to 2015 before declining sharply, whereas HOV-4 remained stable at consistently lower levels (Fig. 3B). Thus, HRV-1 and HRV-3 showed greater temporal variation in relative genetic diversity than HOV-2 and HOV-4.

**Figure 3.**
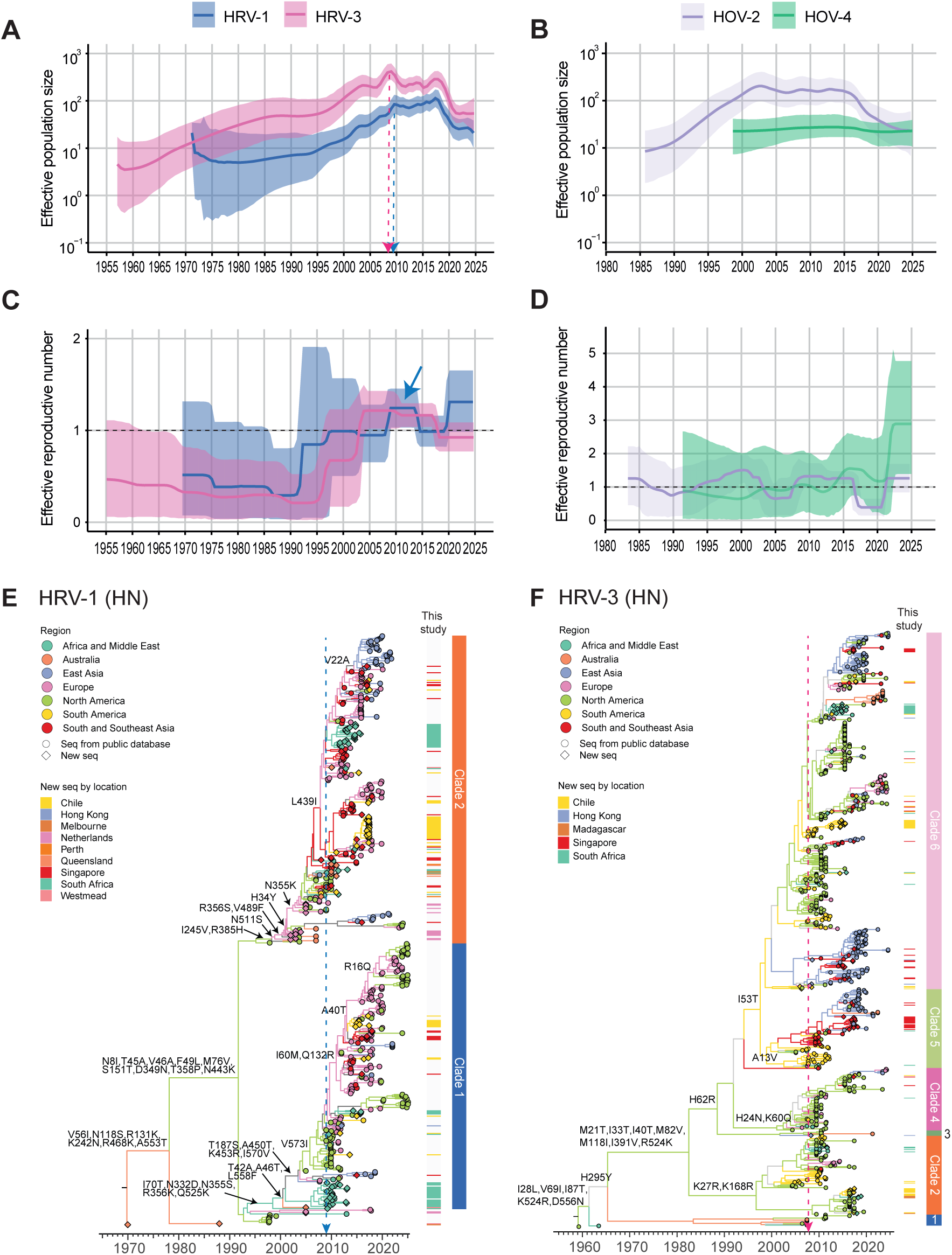
Changes in effective population size and transmission dynamics of the four HPIV types. Estimates of effective population size (N_e_) for (*A*) HRV-1 and HRV-3 and (*B*) HOV-2 and HOV-4 were inferred using the Bayesian skyride analysis with a Gaussian Markov random field (GMRF) coalescent model implemented in BEAST. The blue and pink vertical dashed lines indicate the peak relative genetic diversity for HRV-1 in 2009 and HRV-3 in 2008, respectively. Temporal changes in the effective reproductive number (R_e_) for (*C*) HRV-1 and HRV-3 and (*D*) HOV-2 and HOV-4, estimated using the birth-death skyline (BDSKY) model. The blue arrow denotes the R_e_ above 1 for HRV-1. The grey horizontal dashed line indicates R_e_ = 1. Solid lines represent posterior mean estimates, and shaded areas indicate the 95% highest posterior density (HPD) intervals. Time-scaled phylogenies of the HN genes of (*E*) HRV-1 and (*F*) HRV-3. Maximum clade credibility (MCC) trees were inferred using Bayesian skyride analyses in BEAST, illustrating lineage diversification before the COVID-19 pandemic followed by continued circulation in the post-pandemic era. Coloured branches denote the geographic locations. Diamonds at the tips denote sequences generated in this study, while open circles represent sequences obtained from NCBI GenBank. Amino acid substitutions were mapped at major nodes. The geographic locations of newly generated sequences and clade numbering are indicated by bars.

We estimated the effective reproductive number (R_e_) for each virus type using independent Bayesian analyses. Before 2004, HRV-1 and HRV-3 had R_e_<1, consistent with limited transmission (Fig. 3C). HRV-3 R_e_ subsequently exceeded 1 around 2004, followed by HRV-1 in 2008. HRV-1 remained above 1 for approximately six years (2008–2014; mean R_e_ ∼1.24), whereas HRV-3 remained above 1 for approximately 13 years (2004–2017) and peaked at 1.22 (blue arrow in Fig. 3C). These periods coincided with peaks in relative genetic diversity in 2008 for HRV-3 and 2009 for HRV-1 (dotted vertical lines in Fig. 3A), suggesting that increased transmission contributed to lineage expansion and genetic diversification.

During the COVID-19 pandemic, HRV-3 transmissibility declined to near the epidemic threshold (R_e_ ∼1), consistent with disruption of respiratory virus transmission by non-pharmaceutical interventions such as lockdowns and reduced international travel. HRV-3 R_e_ remained <1 after the pandemic. In contrast, HRV-1 exhibited increased transmissibility, R_e_ increased to ∼1.65 during 2020–2024, exceeding its 2008–2014 level (Fig. 3C). HOV-2 showed intermittent periods of R_e_>1 during 1993–2002, 2008–2016, and 2021–2024 (range: 1.26–1.51), whereas HOV-4 showed more sustained R_e_>1 from 2007 to 2019 and increased after the pandemic to 2.89 (Fig. 3D). The R_e_>1 for HRV-1, HOV-2 and HOV-4 after 2020 indicates ongoing transmission of these types despite non-pharmaceutical interventions, although estimates for this period are based on few sequences and have wide HPD intervals (Fig. 3C and D). Because these skyline estimates depend on the number and temporal distribution of sequences, they can also be affected by changes in testing and sampling over time.

Greater relative genetic diversity and transmissibility in HRV-1 and HRV-3 were associated with lineage expansion (vertical dotted lines in Fig. 3E and F indicate peak relative genetic diversity in 2009 for HRV-1 and 2008 for HRV-3). For HRV-3, peak diversity followed the increase in R_e_ above 1 by approximately four years, whereas for HRV-1 the two coincided, consistent with the time required for lineages to accumulate and diversify under sustained transmission. Multiple smaller clades emerged after 2008 and contributed to regional outbreaks. In North America, HRV-1 formed distinct clusters within clade 1 (Fig. 3E), consistent with sustained local community transmission. HRV-3 showed a similar pattern: North American sequences clustered within clades 4 and 6, and larger East Asian clusters occurred in clades 5 and 6 (Fig. 3F). Singaporean sequences were interspersed throughout the phylogeny, consistent with multiple independent introductions. HOV-2 showed greater geographic mixing whereas HOV-4 sequences were predominantly from North America, with introductions into Singapore forming a small, distinct cluster (*SI Appendix,* Fig. S5). Individual gene phylogenies were broadly concordant with the corresponding HN phylogenies across all four HPIV types (*SI Appendix*, Figs. S6–S25).

Following the COVID-19 pandemic, most previously circulating lineages of all four HPIV types were no longer detected in the sampled population (Fig. 3E and F). However, all four types subsequently resumed circulation and continued to diversify in human populations.

### Differential selection pressures among virus types

We then evaluated selection pressures across the four virus types using site-level Single-Likelihood Ancestor Counting (SLAC) and branch-level Mixed Effects Model of Evolution (MEME) analyses. Across the examined periods, HRV-1, HRV-3, and HOV-4 had higher global dN/dS ratios in the phosphoprotein (P) gene (0.3–0.6) than in other genes (Fig. 4A; *SI Appendix*, Table S4); the pattern was similar when all years were combined (*SI Appendix*, Fig. S26 and Table S5). HRV-1, HOV-2, and HOV-4 also had higher mean substitution rates in P (1.04 × 10^−3^–2.47 × 10^−3^ substitutions/site/year) than in other genes (SI Appendix, Fig. S3 and Table S6). The P gene also contained more positively selected sites than other genes, with 53 in HRV-3 and 30 in HRV-1, of which 18 and 10, respectively, were identified by both SLAC and MEME. In contrast, no positively selected sites were detected in HOV-2 and only two in HOV-4 (Table 2). Most positively selected P sites in HRV-1 and HRV-3 occurred in the N-terminal domain (NTD; residues 1–350).

**Figure 4.**
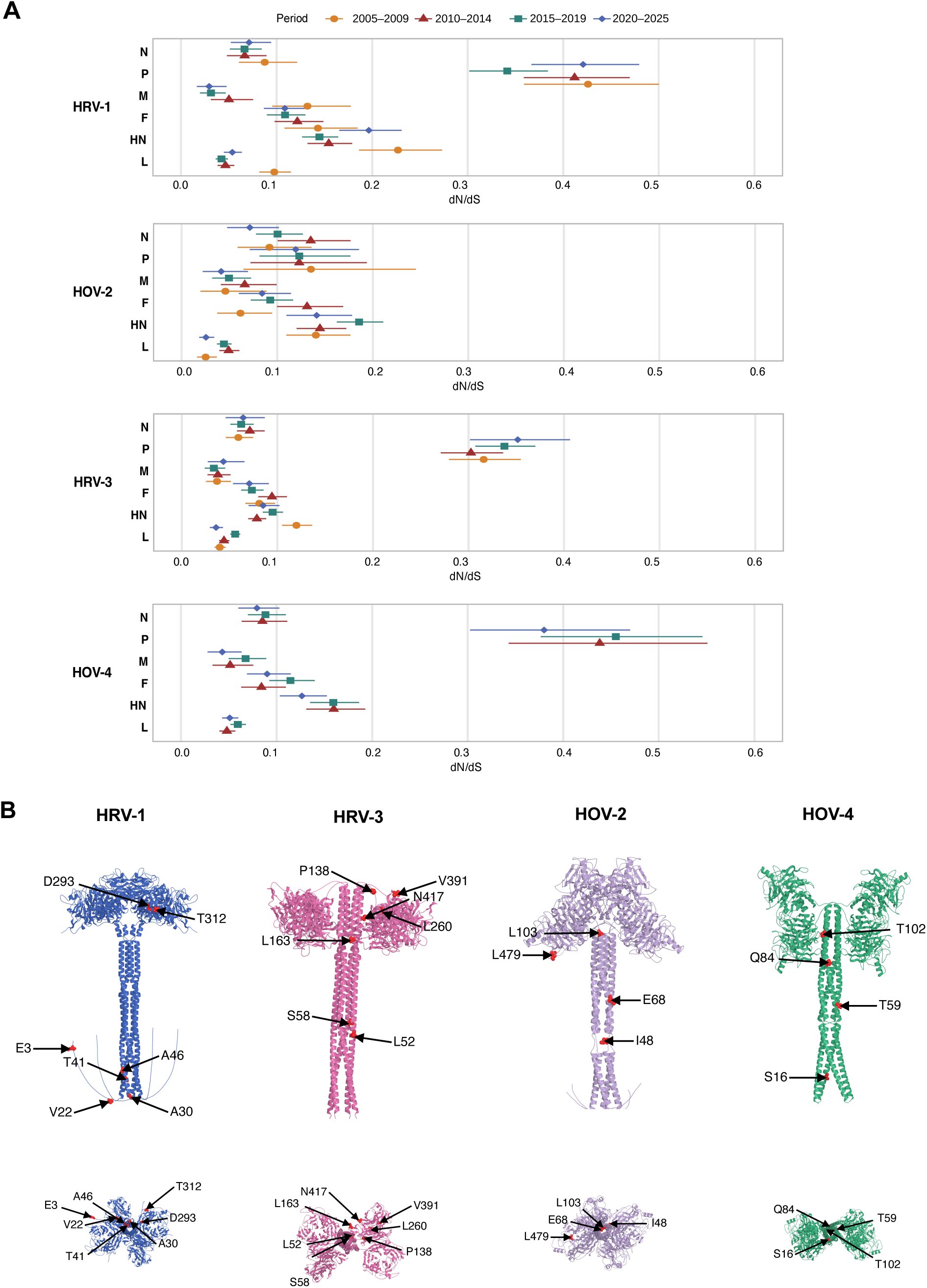
Comparative selection pressures across HPIV genes and positively selected sites in the HN proteins. (*A*) Global d_N_/d_S_ estimates for six viral proteins, including nucleocapsid protein (N), phosphoprotein (P), matrix protein (M), fusion protein (F), hemagglutinin-neuraminidase glycoprotein (HN), and large polymerase protein (L), of the four HPIV types across four time periods: 2005–2009, 2010–2014, 2015–2019, and 2020–2025. Global d_N_/d_S_ ratios were estimated using the single-likelihood ancestor counting (SLAC) method. Means of d_N_/d_S_ ratios are indicated by distinct symbols, with error bars representing 95% confidence intervals. (*B*) Positively selected amino acid sites in the HN protein, identified using SLAC and the mixed effects model of evolution (MEME) methods, mapped onto predicted three-dimensional (3D) HN protein structures for the four HPIV types. HN tetramer structures were predicted using AlphaFold3, with the globular head and stalk domains shown. Positively selected residues are highlighted in red. Structural representations are shown as front (top) and top-down (bottom) views.

**Table 2.**
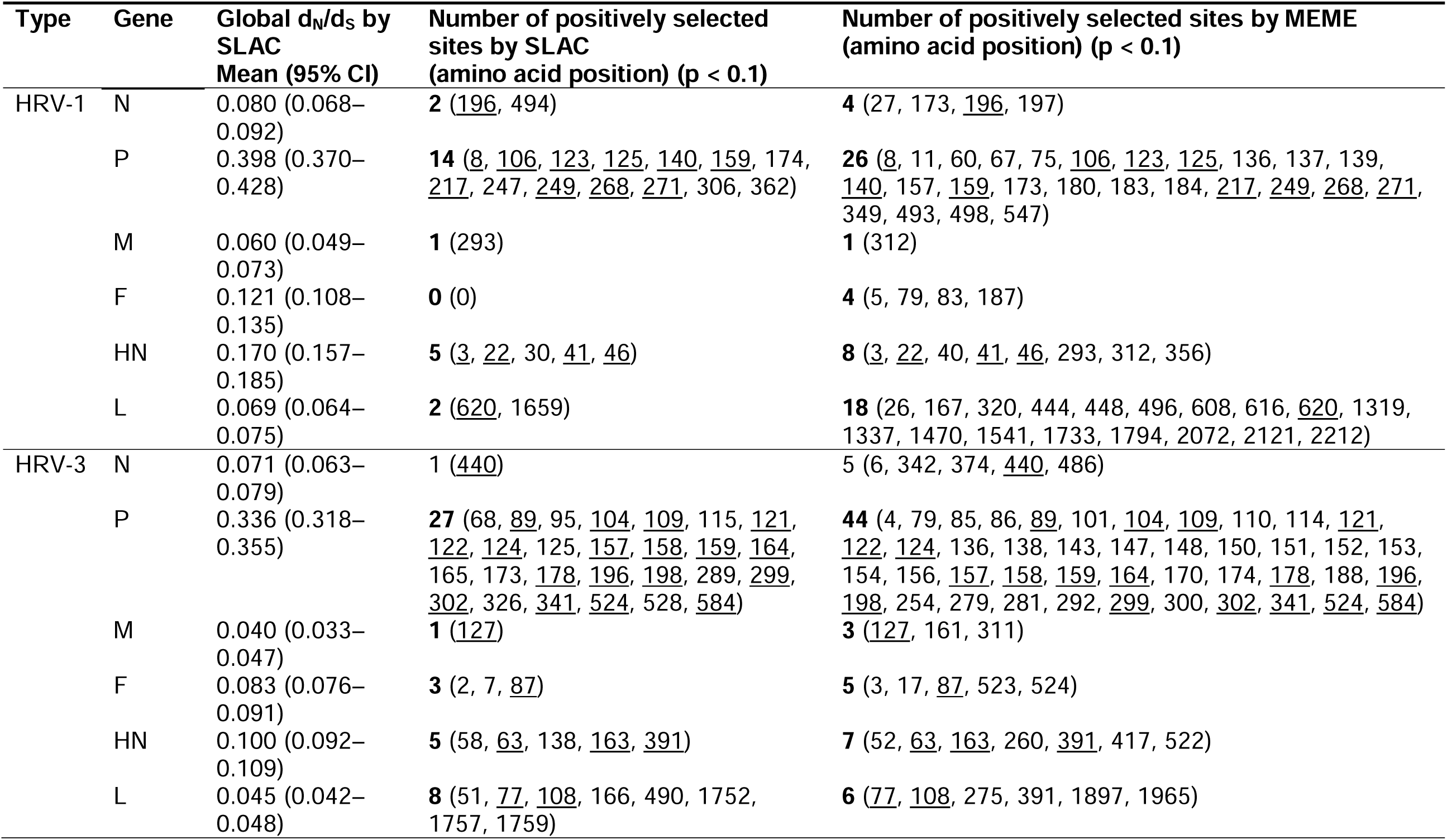
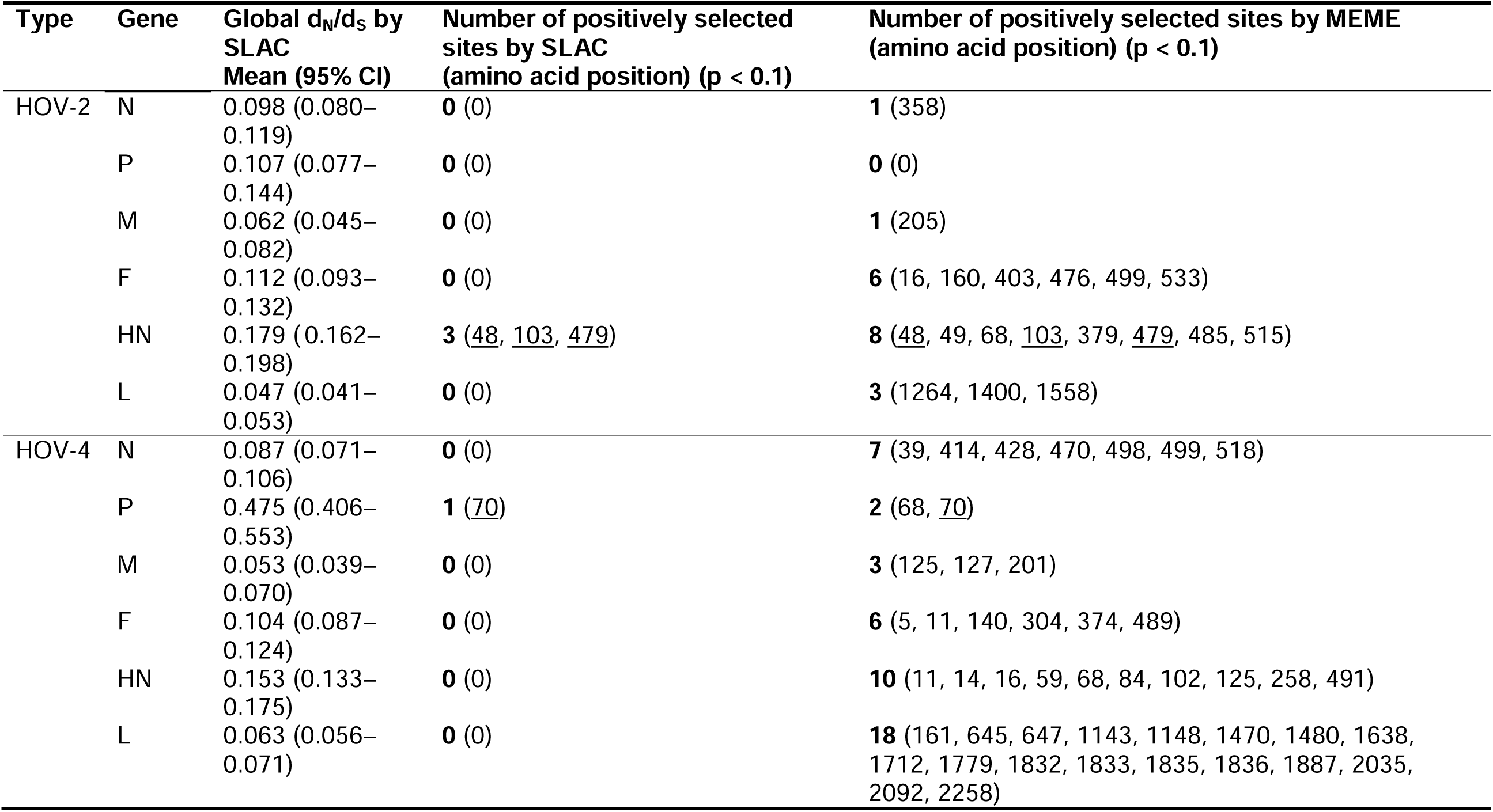
Gene-specific selection pressure estimates for human respiroviruses and human orthorubulaviruses. Sites identified by both SLAC and MEME are underlined.

| **Type** | **Gene** | **Global d<sub>N</sub>/d<sub>S</sub> by SLAC<br>Mean (95% CI)** | **Number of positively selected sites by SLAC<br>(amino acid position) (p < 0.1)** | **Number of positively selected sites by MEME<br>(amino acid position) (p < 0.1)** |
|---|---|---|---|---|
| HRV-1 | N | 0.080 (0.068–0.092) | **2** (196, 494) | **4** (27, 173, 196, 197) |
|  | P | 0.398 (0.370–0.428) | **14** (8, 106, 123, 125, 140, 159, 174, 217, 247, 249, 268, 271, 306, 362) | **26** (8, 11, 60, 67, 75, 106, 123, 125, 136, 137, 139, 140, 157, 159, 173, 180, 183, 184, 217, 249, 268, 271, 349, 493, 498, 547) |
|  | M | 0.060 (0.049–0.073) | **1** (293) | **1** (312) |
|  | F | 0.121 (0.108–0.135) | **0** (0) | **4** (5, 79, 83, 187) |
|  | HN | 0.170 (0.157–0.185) | **5** (3, 22, 30, 41, 46) | **8** (3, 22, 40, 41, 46, 293, 312, 356) |
|  | L | 0.069 (0.064–0.075) | **2** (620, 1659) | **18** (26, 167, 320, 444, 448, 496, 608, 616, 620, 1319, 1337, 1470, 1541, 1733, 1794, 2072, 2121, 2212) |
| HRV-3 | N | 0.071 (0.063–0.079) | 1 (440) | 5 (6, 342, 374, 440, 486) |
|  | P | 0.336 (0.318–0.355) | **27** (68, 89, 95, 104, 109, 115, 121, 122, 124, 125, 157, 158, 159, 164, 165, 173, 178, 196, 198, 289, 299, 302, 326, 341, 524, 528, 584) | **44** (4, 79, 85, 86, 89, 101, 104, 109, 110, 114, 121, 122, 124, 136, 138, 143, 147, 148, 150, 151, 152, 153, 154, 156, 157, 158, 159, 164, 170, 174, 178, 188, 196, 198, 254, 279, 281, 292, 299, 300, 302, 341, 524, 584) |
|  | M | 0.040 (0.033–0.047) | **1** (127) | **3** (127, 161, 311) |
|  | F | 0.083 (0.076–0.091) | **3** (2, 7, 87) | **5** (3, 17, 87, 523, 524) |
|  | HN | 0.100 (0.092–0.109) | **5** (58, 63, 138, 163, 391) | **7** (52, 63, 163, 260, 391, 417, 522) |
|  | L | 0.045 (0.042–0.048) | **8** (51, 77, 108, 166, 490, 1752, 1757, 1759) | **6** (77, 108, 275, 391, 1897, 1965) |
| **Type** | **Gene** | **Global d<sub>N</sub>/d<sub>S</sub> by SLAC<br>Mean (95% CI)** | **Number of positively selected sites by SLAC<br>(amino acid position) (p < 0.1)** | **Number of positively selected sites by MEME<br>(amino acid position) (p < 0.1)** |
| HOV-2 | N | 0.098 (0.080–0.119) | **0** (0) | **1** (358) |
|  | P | 0.107 (0.077–0.144) | **0** (0) | **0** (0) |
|  | M | 0.062 (0.045–0.082) | **0** (0) | **1** (205) |
|  | F | 0.112 (0.093–0.132) | **0** (0) | **6** (16, 160, 403, 476, 499, 533) |
|  | HN | 0.179 ( 0.162–0.198) | **3** (48, 103, 479) | **8** (48, 49, 68, 103, 379, 479, 485, 515) |
|  | L | 0.047 (0.041–0.053) | **0** (0) | **3** (1264, 1400, 1558) |
| HOV-4 | N | 0.087 (0.071–0.106) | **0** (0) | **7** (39, 414, 428, 470, 498, 499, 518) |
|  | P | 0.475 (0.406–0.553) | **1** (70) | **2** (68, 70) |
|  | M | 0.053 (0.039–0.070) | **0** (0) | **3** (125, 127, 201) |
|  | F | 0.104 (0.087–0.124) | **0** (0) | **6** (5, 11, 140, 304, 374, 489) |
|  | HN | 0.153 (0.133–0.175) | **0** (0) | **10** (11, 14, 16, 59, 68, 84, 102, 125, 258, 491) |
|  | L | 0.063 (0.056–0.071) | **0** (0) | **18** (161, 645, 647, 1143, 1148, 1470, 1480, 1638, 1712, 1779, 1832, 1833, 1835, 1836, 1887, 2035, 2092, 2258) |

The dN/dS ratios of the HN genes were comparable across the four types (0.1–0.17; *SI Appendix*, Fig. S26 and Table S5). However, the dN/dS ratio of HRV-1 HN was higher during 2020–2025 than during 2015–2019, whereas that of HRV-3 declined moderately after the COVID-19 pandemic (Fig. 4A). This pattern was consistent with greater accumulation of trunk amino acid substitutions in HRV-1 HN during the 2020s (Fig. 4A). HRV-1 and HRV-3 had similar numbers of positively selected HN sites, but their distributions differed (Table 2). In HRV-1, two sites (D293L and K355S) were in the globular head and five were in the stalk. HRV-3 HN had four sites in the head (residues 138, 260, 391, and 417) and three in the stalk (residues 52, 58, and 163) (Fig. 4B; Table 2). HOV-2 and HOV-4 had fewer positively selected HN sites, predominantly in the stalk (Fig. 4B; Table 2). Positively selected sites also occurred in the F proteins of all four types, with variable distributions (Table 2). Co-occurring F and HN substitutions within the same lineages suggested potential epistatic interactions. For example, HRV-3 F L7S and HN A13V co-occurred with the emergence of clade 5 (Fig. 2B), suggesting that coordinated changes in these proteins may have contributed to lineage diversification.

To assess the potential antigenic effects of these evolutionary changes, we performed *in silico* B-cell epitope analyses of F and HN proteins. Several predicted epitope regions were shared across the four virus types, including regions in F (e.g., 95–110, 360–375, 400–410, 435–445, and 540–555) and HN (e.g., 3–30, 130–157, 215–221, 257–264, 275–285, 341–380, 457–471, and 552–556) (Fig. 5A). HRV-3 HN residue 138, a positively selected site, overlapped a predicted epitope. Phylogenetic mapping showed that L138P arose independently in multiple clades, with a minority of viruses carrying L138Q (*SI Appendix*, Fig. S27). Other positively selected sites, including residues 58, 63, 163, and 391, also showed independent substitutions, although most occurred as singletons (*SI Appendix*, Figs. S28–S31). Predicted B-cell epitopes in HRV-3 were concentrated on the lateral region of the globular head, whereas those in HRV-1 were primarily localized to the globular head. HOV-2 showed a more diffuse distribution across the head and stalk, whereas HOV-4 epitopes were concentrated toward the lower head region (Fig. 5B).

**Figure 5.**
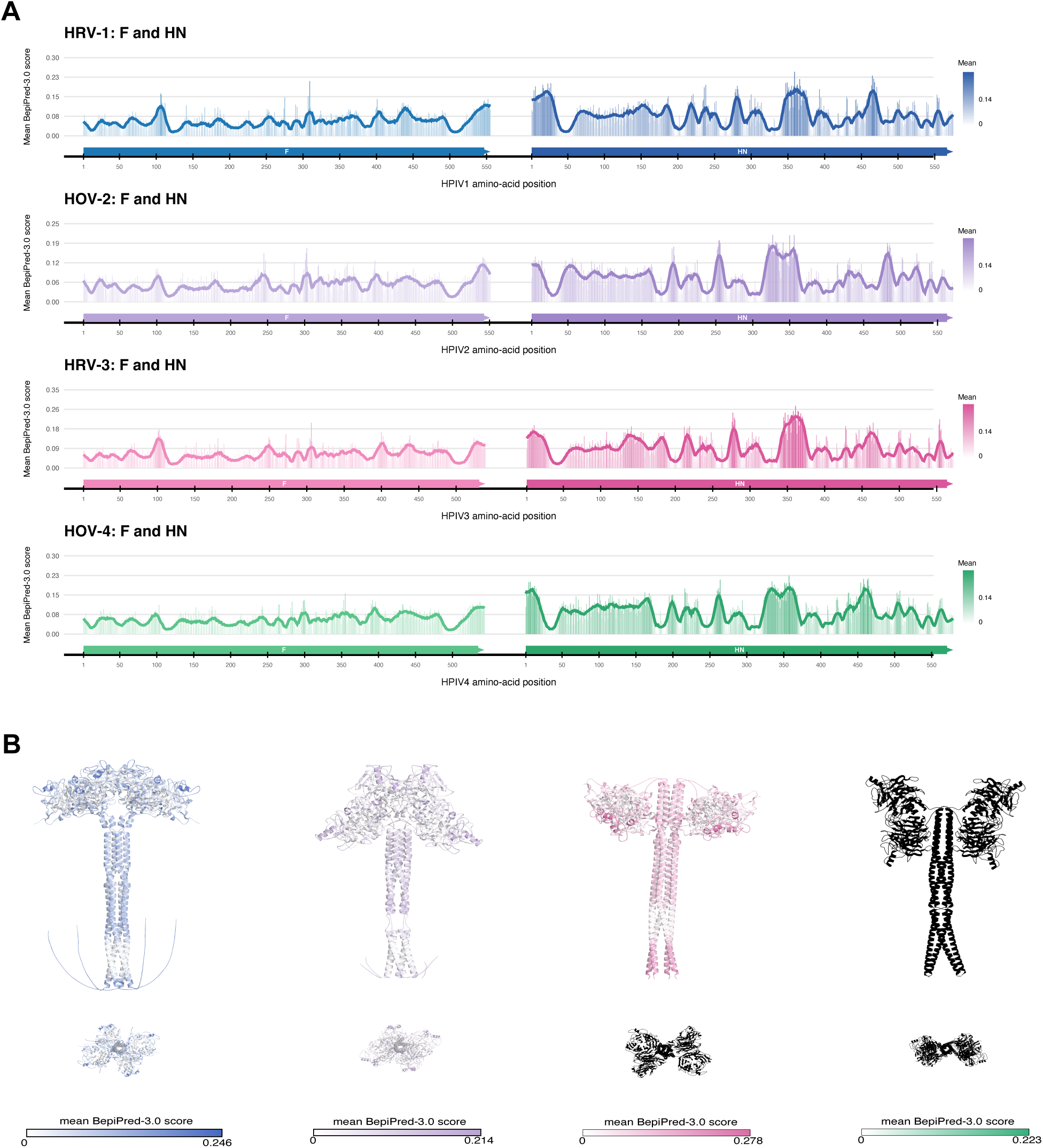
Epitope prediction in the F and HN proteins of the four HPIV types. (*A*) Mean B-cell epitope prediction scores for each amino-acid position across the F (left) and HN (right) genes of HRV-1, HOV-2, HRV-3, and HOV-4 (top to bottom). BepiPred-3.0 epitope prediction scores were calculated for each sequence at every amino acid position and averaged across all analysed sequences. *(B)* AlphaFold3-predicted HN tetramer structures for HRV-1, HOV-2, HRV-3, and HOV-4 (left to right), shown as full side views (globular head and stalk domains) and top-down views of the head domain. Structures are colour-coded by mean BepiPred-3.0 score, with white indicating low predicted epitope probability and darker colour indicating higher predicted epitope probability.

Together, these findings suggest stronger host-mediated selection on the HN and P proteins of HRV-1 and HRV-3. In HRV-3, selection was concentrated in the globular HN head, where predicted B-cell epitopes were also located, whereas positively selected sites were largely absent from the globular heads of HOV-2 and HOV-4. These selection pressures may have contributed to greater lineage expansion, onward transmission, and broader global circulation of HRV-1 and HRV-3 than of the other two types.

### Spatiotemporal diffusion patterns of HRV and HOV

We used Bayesian phylogeographic reconstruction to infer the spatiotemporal diffusion of HRV and HOV lineages. Discrete phylogeographic analyses used Bayesian stochastic search variable selection (BSSVS), and continuous analyses used a relaxed random walk (RRW) model. HRV-1 was inferred to have originated in Australia before dispersing to North America in the early 1980s (Fig. 6A). During the 2000s, North America became a major source of global dissemination, with inferred transmission to Europe, Africa, and Australia, followed by expansion into South America and Southeast Asia during the 2010s (Fig. 6B).

**Figure 6.**
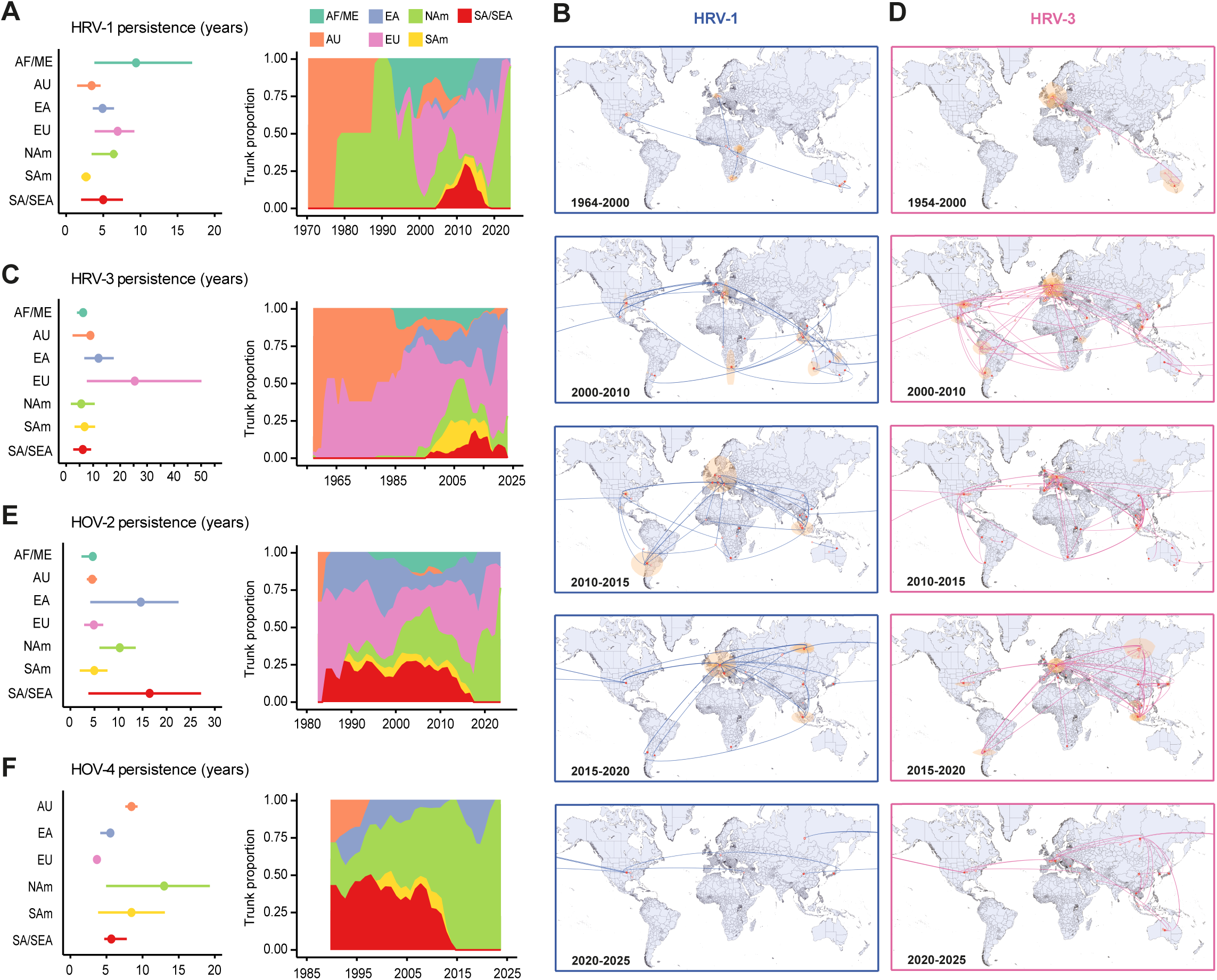
Spatiotemporal dynamics of the four HPIV types inferred from discrete and continuous phylogeographic analyses. (*A*) Discrete phylogeography of HRV-1, showing estimated viral persistence across geographic regions. Coloured bars denote different locations, with dots indicating the mean persistence duration (years) and horizontal bars representing 95% HPD intervals. Trunk proportions over time are shown for HRV-1. (*B*) Continuous phylogeographic diffusion of HRV-1 across five time periods. Spatial diffusion was modelled in continuous geographic space using latitude and longitude as bivariate traits under a Cauchy relaxed random walk (RRW) model. (*C*) Discrete phylogeography of HRV-3. Geographic regions are abbreviated as follows: AF/ME, Africa and Middle East; EA, East Asia; NAm, North America; SA/SEA, South and Southeast Asia; AU, Australia; EU, Europe; and SAm, South America. (*D*) Continuous phylogeographic diffusion of HRV-3 across five time periods. Spatial diffusion was modelled in continuous geographic space using latitude and longitude as bivariate traits under a Cauchy relaxed random walk (RRW) model. Shaded areas denote 80% highest posterior density (HPD). (*E*) Discrete phylogeography of HOV-2. (*F*) Discrete phylogeography of HOV-4, showing estimated viral persistence and trunk proportions over time.

During 2015–2020, China and Europe emerged as major transmission hubs, with Europe showing substantial diffusion into North America, South Asia, and Southeast Asia (Fig. 6A). Diffusion rates from Europe to these regions were statistically significant (*SI Appendix*, Fig. S32 and Table S12). Viral persistence was estimated at approximately five years in most regions but up to 13 years in Africa and the Middle East (Fig. 6A; SI Appendix, Table S7).

HRV-3 was also inferred to have originated in Australia before spreading to Europe in the 1960s; both regions subsequently became major sources of global transmission (Fig. 6C). During the 2000s, HRV-3 spread widely to North America, South America, and Africa, followed by further spread to South and Southeast Asia during 2010–2015. Global transmission intensified after 2015 (Fig. 6D), with statistically significant diffusion from Europe to North America (*SI Appendix*, Fig. S32 and Table S13). Europe showed the longest viral persistence (∼40 years) and was a major source of onward dissemination, with high diffusion rates to North America, South America, Africa, and South and Southeast Asia (Fig. 6D; SI Appendix, Table S7).

Unlike HRV-1 and HRV-3, which diversified into multiple clades, HOV-2 and HOV-4 each comprised two co-circulating clades. Despite its lower detection frequency, HOV-2 likely originated in Europe. Multiple transmission sources, particularly Europe and South and Southeast Asia, contributed to local epidemics by the early 2000s. After 2005, North America emerged as a dominant source, with strong epidemiological links to Africa, the Middle East, and Europe; Europe was also linked to East Asia (*SI Appendix*, Fig. S33 and Table S14).

South and Southeast Asia showed the longest estimated viral persistence (approximately 24 years), followed by East Asia (18 years; Fig. 6E; *SI Appendix*, Fig. S32 and Table S7). HOV-4 also showed extensive interregional circulation, with sustained circulation among North America, Australia, East Asia, and South and Southeast Asia during the mid-2000s (*SI Appendix*, Fig. S33 and Table S15). South and Southeast Asia appeared to be the major source of HOV-4 dissemination to North America and East Asia. North America showed the longest local persistence (approximately 15 years), followed by South America (10 years; Fig. 6F). Together, repeated seeding, prolonged regional persistence, and bidirectional transmission appear to underpin the global spread of HRV and HOV.

## Discussion

Here, we substantially expand the genomic landscape of human parainfluenza viruses by generating 279 whole-genome sequences from five continents across 30 years, increasing the number of genomes publicly available by up to 30%. By reconstructing their evolutionary history, we show that selection pressure and transmission dynamics differed markedly between HPIV types that share a respiratory niche, with HRV-1 and HRV-3 showing more positively selected sites, greater lineage diversification, and higher effective reproductive numbers than HOV-2 and HOV-4. This analysis also revealed persistent gaps in global genomic surveillance, particularly after the COVID-19 pandemic. Addressing these surveillance gaps will be essential for tracking the lineages that sustain HPIV circulation and for guiding the design of broadly protective vaccines and therapeutics.

Phylodynamic analyses revealed distinct evolutionary and epidemiological trajectories among the four HPIV types despite their shared respiratory niche. After the COVID-19 pandemic disrupted viral circulation, all four types re-emerged, with recurrent outbreaks most evident for HRV-1 and HRV-3. We observe that stronger seasonal fluctuations and ongoing genetic drift characterize HRV-1 and HRV-3, whereas HOV-2 and HOV-4 show weaker seasonal dynamics. The diversification of HRV-3 from the late 2000s to the early 2010s coincided with the influenza A(H1N1)pdm09 pandemic, which could have influenced transmission through changes in population immunity, such as viral interference between respiratory viruses, in contact patterns, such as school closures and behavioural change, or in testing. However, this overlap may be coincidental, because HRV-3 relative genetic diversity peaked in December 2008, before the pandemic began, and its effective reproductive number was already above 1 from around 2004.

Before the COVID-19 pandemic, HRV-1 and HRV-3 diversified into multiple genetic lineages, with HRV-3 exhibiting a more pronounced ladder-like phylogeny, resembling influenza A and the B/Victoria lineage viruses, in which successive lineages replace one another, complicating influenza vaccine strain selection by necessitating a change in strain every 1-2 years (26, 27). In contrast, HRV-1 comprised two major clades that diverged in the early 1990s and co-circulated for 30 years, resembling influenza B/Yamagata with lineages that persisted for up to 6 years before the virus was last detected in March 2020. Given their roles in receptor binding, membrane fusion, and immune recognition, HN and F might be expected to experience stronger selection, analogous to hemagglutinin (HA) in seasonal influenza A and B viruses (26, 28). However, we identified more positively selected sites and higher *dN/dS* values in the P gene than in HN or F. These findings suggest that diversification of HRV-1 and HRV-3 is not confined to surface glycoproteins and may also involve selection on proteins involved in viral replication and host–virus interactions, consistent with patterns observed in influenza A and B viruses (26). In contrast, positive selection in RSV is largely restricted to the G attachment glycoprotein (29), and in HCoV-229E and HCoV-OC43 it primarily targets spike domains A and B (30), indicating that selection is more concentrated on surface proteins than in HRV-1 and HRV-3.

The potential functional relevance of specific amino acid substitutions is supported by several studies on HRV-3 HN. During prolonged infection in immunocompromised patients, naturally arising HN substitutions, including H552Q, were associated with increased sialic acid receptor avidity and enhanced fusion-promoting activity (31). Moreover, removal of the N-linked glycan at HN residue 523 altered receptor specificity, delayed erythrocyte elution, and reduced viral growth in LLC-MK2 cells, demonstrating that changes near a putative secondary receptor-binding region can affect viral phenotype (32). In this context, the positively selected HN residue 522 identified in our analysis is adjacent to residue 523 and lies within an antibody-recognized region encompassing residues 513–534, while the predicted B-cell epitope at residues 215–221 overlaps a flexible loop implicated in receptor recognition (33). Likewise, the positively selected HN residue 138 lies near a reported antibody-contact region around residue 139 (34). These spatial relationships suggest plausible antigenic relevance of the substitutions detected here, including altered receptor binding, fusion, or antibody sensitivity of HRV-3. The discovery of these mutations also has potential clinical implications. Alterations in HN receptor engagement and HN-F coordination could influence viral fitness and persistence in particular host environments, including during prolonged infection in immunocompromised patients (31). In addition, structural and functional studies have shown that human monoclonal antibodies (mAbs) targeting prefusion F or HN can neutralize HRV-3 *in vitro* and protect against infection or severe disease *in vivo* (33, 34).

Our analysis indicated that HRV-1 emerged in Australia before dispersing to North America, which became an important source of global dissemination. Together with extensive interregional circulation across all four types (SI Appendix, Tables S8–S11), these findings support roles for regional persistence and repeated geographic dissemination in sustaining global HPIV circulation. Overall, HPIV lineages showed more local transmission and persistence and less global transmission compared to seasonal influenza viruses (24, 26), a pattern also reported for other respiratory viruses such as seasonal human coronaviruses and SARS-CoV-2 (30, 35).

This study has several limitations. First, despite multi-national collection, samples were unevenly distributed across locations and years, and sequence availability was limited during and after the COVID-19 pandemic, possibly reflecting disrupted circulation, reduced testing for viruses other than SARS-CoV-2, or both. Although all four types resumed circulation after the pandemic, the apparent decline of some clades and estimates of effective population size and reproductive number may be biased by reduced sampling and testing.

Sustained, geographically balanced surveillance is needed to characterize long-term evolutionary and transmission dynamics. Second, small and uneven datasets for HOV-2 and HOV-4, partly reflecting their relative rarity, limited the precision of phylogeographic inference, which is sensitive to the presence or absence of sequence data. The earliest sequences are exclusively from Australia (1970–1988), which may bias inferred origins such as that of HRV-1, and fewer than 10 HOV-4 sequences were available in most years, with none for 2012, 2013, or 2021. Third, the antigenic significance of positively selected residues was inferred from predicted structures and proximity to putative B-cell epitopes, which do not establish antibody binding or functional effects and require experimental validation. Furthermore, antigenic characterization was not possible because available HPIV strains are predominantly older and strain-matched antisera and reagents are not readily available.

The WHO Global Influenza Surveillance and Response System (GISRS), which was previously limited to influenza, now also conducts RSV surveillance as RSV vaccines are introduced, with its current phase from 2025 to 2027 estimating RSV-associated burden in young children in 24 low- and middle-income countries to inform national vaccine policy (36). However, HPIV accounts for increasing numbers of hospitalized cases that are comparable to RSV in both children and older adults (37), and HPIV is the second most common cause of childhood hospitalization for acute lower respiratory infection after RSV (38). Increased clinical surveillance and genomic characterization of HPIV, including antigenic characterization and the development of reference reagents and strains, is needed to monitor emerging mutations and guide vaccine and therapeutic development.

Together, these findings demonstrate that HPIV evolution is shaped by the persistence and co-circulation of multiple lineages, largely within restricted geographic regions, rather than the global expansion of a single dominant lineage that characterises influenza evolution, highlighting the need for regional genomic surveillance of locally persistent lineages. The lower dN/dS ratios of HN and F than of P indicate stronger purifying selection, supporting the surface glycoproteins HN and F as targets of antibody-mediated protection. However, the higher evolutionary rates and positively selected sites in the P gene of HRV-1 and HRV-3 suggest that non-surface proteins may also contribute to viral adaptation and warrant further evaluation. Reverse-genetic approaches and appropriate animal models will be valuable for determining the functional effects of nonsynonymous substitutions on viral fitness and pathogenesis.

## Materials and Methods

### Ethics statement

This study was approved by the ethics committees of the National University of Singapore Institutional Review Board (NUS-IRB; reference codes: 13-256E and 2022-320). All clinical samples were de-identified before transfer to the research laboratory at Duke-NUS Medical School.

### Sample collection

A total of 1,161 HPIV-positive nasal swab samples were collected between 1988 and 2018 through seven collaborating laboratories in Australia (*n*=315), Chile (*n*=226), Hong Kong (*n*=38), Madagascar (*n*=75), the Netherlands (*n* = 71), Singapore (*n*=228) and South Africa (*n*=208) (*SI Appendix*, Fig. S1 and Table S1). Viral RNA was extracted from samples using the Direct-zol RNA Miniprep Kit (Zymo Research, USA) according to the manufacturer’s instructions (28). HPIV subtyping was performed by real-time quantitative reverse transcription PCR (RT-qPCR) using previously described primers and probes (39) and AgPath-ID One-Step RT-PCR reagents (Thermo Fisher Scientific, USA).

### Next-generation sequencing and genome assembly

Full genome sequencing was performed on HPIV RNA-positive samples using next-generation sequencing (NGS). Briefly, extracted RNA (1 µg total RNA) was used for direct RNA library preparation with the TruSeq RNA Library Preparation Kit (Illumina International, USA) according to the manufacturer’s instructions. Library quality was assessed using an Agilent Bioanalyzer (Agilent Technologies, USA). Libraries were pooled and sequenced on the Illumina MiSeq platform with 2 x 250-bp paired-end reads. The raw NGS reads were quality-checked and trimmed using Trimmomatic v0.39 (40) to remove adaptor sequences and low-quality reads (30). The trimmed reads were mapped to reference genomes for HRV-1 (AF457102), HRV-3 (AB012132). HOV-2 (X57559), and HOV-4 (AB543336) using the Burrow-Wheeler Aligner (BWA)-maximum extract matches (MEM) algorithm implemented in UGENE v.40 (41, 42). In total, we obtained 279 complete HPIV genomes, comprising 94 HRV-1, 150 HRV-3, 22 HOV-2, and 13 HOV-4 genomes (*SI Appendix*, Table S16). These newly generated whole genomes increased the publicly available genomic datasets by up to 30% (*SI Appendix*, Fig. S1). Additionally, 178 hemagglutinin-neuraminidase (HN) gene sequences were obtained from samples that did not yield complete genomes, including 54 HRV-1, 66 HRV-3, 55 HOV-2, and 3 HOV-4 samples (*SI Appendix*, Table S17).

### Virus classification and sequence distribution

The family *Paramyxoviridae* phylogenetic tree was obtained and modfied from the International Committee on Taxonomy of Viruses Report, based on L protein amino acid sequences and the maximum-likelihood method. Subtrees representing the genera *Respirovirus* and *Orthorubulavirus* were extracted from the family-level phylogeny. The major host species of each virus species were annotated using animal icons. Reference sequences used to depict the genome organization for the four HPIV types were HRV-1 (AF457102), HRV-3 (EU326526), HOV-2 (X57559), and HOV-4 (AB543336). Weekly distributions of HPIV by type and geographic region, as well as monthly numbers of HN sequences by type were plotted based on the number of available HN sequences, including sequences generated in this study (HRV-1, *n*=148; HRV-3, *n*=216; HOV-2, *n*=77; HOV-4, *n*=16) and retrieved from National Center for Biotechnology Information (NCBI) GenBank database (HRV-1, *n*=368; HRV-3, *n*=1,309; HOV-2, *n*=169; HOV-4, *n*=25) (*SI Appendix*, Table S16–18). The distribution and streamline plots were generated using the ggplot2 (43) packages in R.

### Evolutionary analysis

All available whole genomes of the four HPIV types were downloaded from the NCBI GenBank database between 1957 and 2025 (as of 25 March 2025). The final datasets comprised 317 HRV-1, 762 HRV-3, 123 HOV-2 and 68 HOV-4 genomes (*SI Appendix*, Table S17). For each type, multiple sequence alignments were performed independently using MAFFT v7.450 (44). Temporal signal was assessed using Tempest v1.5.3(45), and outlier sequences were excluded prior to phylodynamic analysis. Time-resolved phylogenies were inferred independently for each virus type using BEAST v1.10.5 (46), with an uncorrelated relaxed molecular clock, a Gaussian Markov Random Field (GMRF) Bayesian Skyride coalescent prior and the HKY85 nucleotide substitution model with gamma-distributed rate heterogeneity. The GMRF Bayesian skyline model was used to estimate changes in effective population size over time as a measure of relative genetic diversity (47). For each type, at least four independent Markov chain Monte Carlo (MCMC) runs of 100 million generations were performed, with states sampling every 10,000 generations.

Convergence and adequate mixing were assessed in Tracer v.1.7.2 (48), and the runs were combined after removal of appropriate burn-in, with all parameters achieved sufficient effective sample size (ESS) values. Posterior tree and log files from independent runs were combined usingLogCombiner v1.10.4 (49). For each virus type, nucleotide substitution rates, time to most recent common ancestor (TMRCA), and the past population dynamics were estimated. Maximum clade creditability (MCC) trees were generated usingTreeAnnotator v10.5.0 (46).Amino acid substitutions were inferred and mapped onto the phylogenies using treesub (50). Mean pairwise nucleotide distances within and between HPIV types, as well as inter-clades of each HPIV type were calculated and compared using Molecular Evolutionary Genetics Analysis (MEGA) v12.1.2 (51, 52).

We also downloaded all available HN gene sequences for four HPIV types from the NCBI GenBank. The final HN datasets comprised 548 HRV-1, 287 HOV-2, 762 HRV-3 and 70 HOV-4 sequences. In addition, five other viral genes (N, P, M, F and L) were extracted from full-genome sequences. Time-scaled phylogenies were reconstructed independently for each gene using the same Bayesian phylogenetic framework described above.

### Estimates of effective population size and effective reproductive number

Changes in relative genetic diversity for the four HPIV types were inferred from effective population size (N_e_) estimates generated in Tracer v.1.7.2 (48) using the GMRF smoothing prior. Changes in effective reproductive number (R_e_) over time of the four HPIV types were estimated using BEAST X v10.5.0. R_e_ represent the average number of secondary infections generated by an infected individual in a population and provides an indicator of epidemic growth or decline and the potential impact of interventions (53). Under the birth–death framework, this parameterization is analogous to the rate at which new infectious individuals are generated relative to the rate at which infectiousness terminates. Values of R_e_ greater than 1 indicate epidemic growth, whereas values below 1 indicate epidemic decline (54). To estimate R_e_ for each HPIV type, we used the birth–death skyline serial (BDSKY) model implemented in BEAST X. For each dataset, an optimized relaxed molecular clock model was applied with the HKY85 nucleotide substitution model (55). The clock-rate prior was set to 0.0006 substitutions per site per year, and the kappa prior was set to 5.5. The reproductive number was estimated using 10 dimensions across the study period. Four independent Markov chain Monte Carlo (MCMC) runs of 100 million generations were performed using a Bactrian operator schedule, with sampling every 10,000 generations. Convergence and mixing of the MCMC runs were assessed in Tracer v1.7.2, and appropriate burn-in values were removed before downstream analysis (48). Log files from independent runs were combined using LogCombiner v1.10.4. Posterior mean estimates and 95% highest posterior density (HPD) intervals of R_e_ over time were calculated and visualized using the R package bdskytools (35). Final plots were generated using ggplot2 (43) in R.

### Selection analysis

Selection pressure was compared among the four HPIV types for each individual gene. The ratio of non-synonymous to synonymous substitutions (d_N_/d_S_) was estimated using the FitMG94 model in HyPhy. Codon sites subject to episodic positive selection were identified using the Single-Likelihood Ancestor Counting (SLAC) method in HyPhy and the Mixed Effects Model of Evolution (MEME) implemented in Datamonkey v3, with statistical significance defined as *P* < 0.1. Positively selected amino acid residues were mapped onto the predicted three-dimensional (3D) structures of the HN proteins for each virus type, generated using AlphaFold 3 (56), and visualized using PyMOL v2.3.5 (57). Amino acid substitutions were reconstructed using treesub and mapped onto the corresponding phylogenies and protein structures.

### B-cell linear epitope prediction

Linear B-cell epitopes were predicted for the F and HN glycoproteins of each virus using BepiPred-3.0 (58), which combines sequence-based features with ESM-2 protein language model embeddings to generate a per-residue epitope propensity score. For each virus type, the mean predicted score at each amino-acid position was then calculated across all sequences to generate a consensus epitope profile. Positional scores were plotted against amino-acid position for each protein where a high score indicates a greater likelihood that the residue is surface-exposed and antigenic. HN sequence epitope profiles were mapped onto representative structural models generated using AlphaFold 3 and visualised inPyMOL v2.3.5 using a white-to-colour gradient to visualize the spatial distribution of predicted epitopes on the folded protein.

### Phylogeographic analysis

Discrete phylogeographic inference was conducted independently for each of four virus type in BEAST X v10.5.0 (59) using the previously inferred time-resolved MCC tree as a fixed topology. HN sequences with associated sampling locations were grouped into seven geographic regions: Australia, Africa and the Middle East (AfricaME), East Asia, Europe, North America, South America, and South and Southeast Asia (SSEAsia). An asymmetric substitution matrix was used to estimate non-reversible transition rates among regions.

Bayesian Stochastic Search Variable Selection (BSSVS) was implemented in BEAST, and Bayes factor (BF) support for inferred transition rates was summarized using SpreaD3 (60). The PACT program was used to estimate trunk rewards and regional persistence (30, 35). Significant migration pathways were summarized based on the combination of both BF > 3 and the mean indicator of > 0.5. We defined the degree of rate support as follows: BF≥ 1,000 indicates decisive, 100 ≤ BF < 1,000 indicates very strong support, 10 ≤ BF < 100 indicates strong support and 3≤ BF < 10 indicates supported (24).

In addition, we performed continuous phylogeography independently for each virus type to reconstruct the spatial dispersal patterns through time and space. Sampling coordinates were assigned from latitude and longitude data generated using thetidygeocoder package (61).Markov chain Monte Carlo (MCMC) analyses were run for 100 million states, with parameters and trees sampled every 10,000 steps. Sequence evolution was modelled using an HKY substitution model with gamma-distributed among-site rate heterogeneity with four categories. A Bayesian SkyGrid coalescent prior was used for demographic reconstruction, with 50 log population-size parameters, 49 grid points, and a cutoff of 9 years. Branch rates for sequence evolution were modelled under an uncorrelated lognormal relaxed molecular clock. Spatial diffusion was modelled in continuous geographic space using latitude and longitude as a bivariate trait under a Cauchy relaxed random walk model. Convergence was assessed in Tracer v1.7.2, and all parameters achieved ESS values than 200. Spatial diffusion was summarized using the spread command and visualized using spread.gl (62) to display inferred phylogeographic dispersal patterns in geographic space.

## Acknowledgements

This study was supported and funded by Singapore Academic Research Fund Tier 3 Project ID MOE-000095-01 (MOE-MOET32020-0002) and the FONDECYT grants 1162791 and 1212023 from Agencia Nacional de Investigacion y Desarrollo (ANID) of Chile. GJDS and YCFS are supported by the Duke-NUS Signature Research Programme funded by the Ministry of Health, Singapore and by contract 75N93021C00016 from the National Institute of Allergy and Infectious Diseases, National Institutes of Health, Department of Health and Human Services, USA. PC is supported by the National Research Foundation, Singapore, under its Postdoctoral Award (FY2025-NRF-PA-0002).

## Additional information

Accession codes: The new sequences generated in this study have been deposited in GenBank database under accession numbers XXXXXX to XXXXXX (*SI Appendix,* Table S16–17).

## Author contributions

G.J.D.S. and Y.C.F.S. conceived and designed the research. P.C., R.Z., B.W., D.C., Y.Z., W.F.Y., J.M. and Y.C.F.S. analysed the data. Y.Z., W.F.Y., C.Y.S., G.S.M. and G.G.K.N. performed the experiments. G.J.D.S., Y.S.O., A.S.J., D.-Y.W., T.G.-S., J.M.H., M.S., T.P.S., F.K.T., A.M.C., M.F., J.D.D., N.H.R., B.J.C., D.E.D., C.L., C.C., D.W.S., R.A.M. and Y.C.F.S. contributed materials and/or analysis tools. G.J.D.S., Y.S.O., A.S.J., D.-Y.W. and Y.C.F.S. contributed to securing funding for the broader research programme that supported the continuation of this work. P.C. and Y.C.F.S. prepared the figures. G.J.D.S., P.C. and Y.C.F.S. drafted the manuscript. All authors reviewed and edited the manuscript.

## Data and materials availability

All data are available in the main text or the supplementary materials.

---

**Appendix Figure S1.**
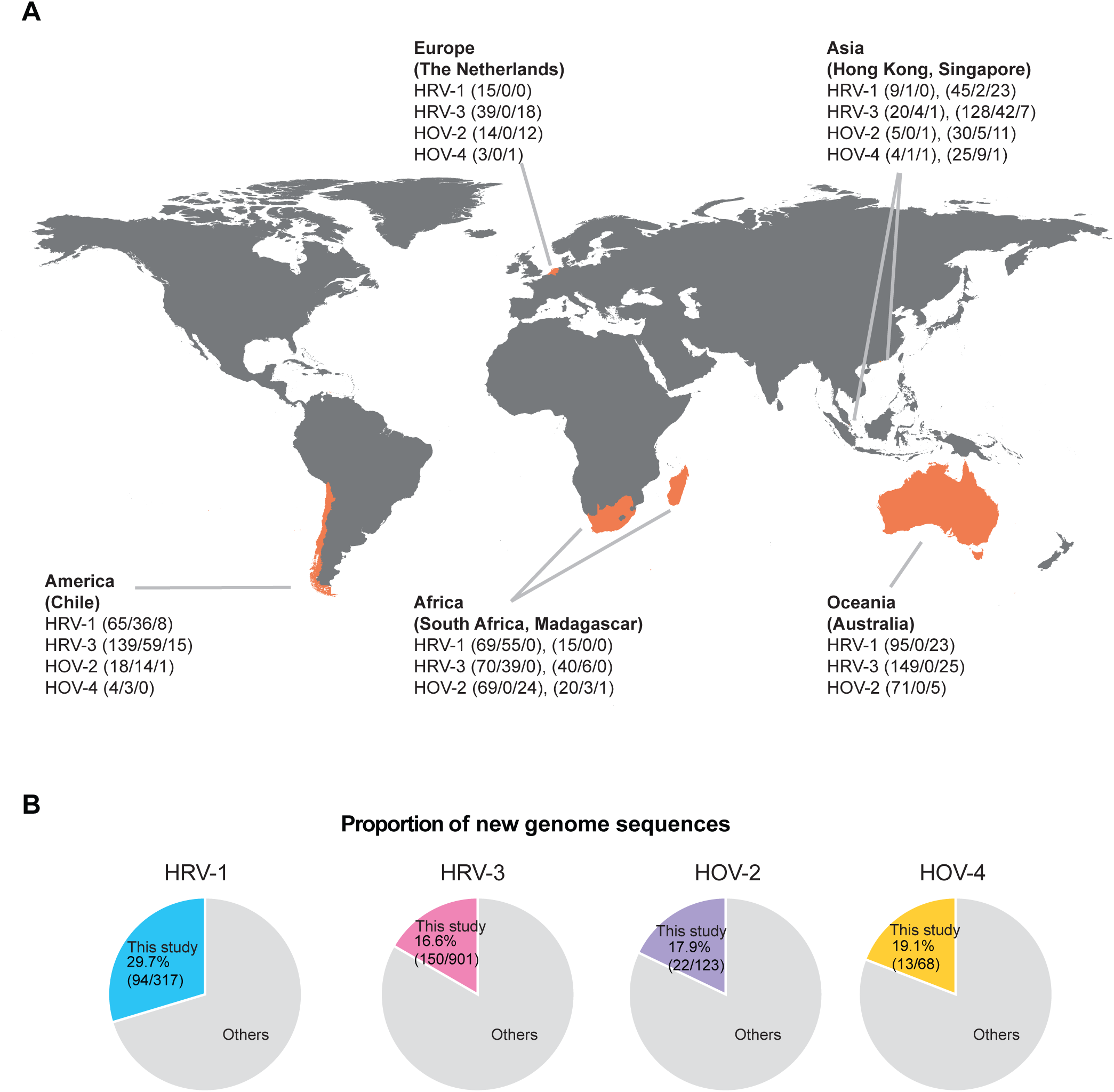
Multi-national collection of clinical samples of human parainfluenza viruses. (*A*) Sample collection sites across seven geographic regions, including Australia, Chile, Hong Kong, Madagascar, the Netherlands, Singapore, and South Africa, are marked in orange. For each virus type, the numbers in brackets represent the total number of samples collected at each location, followed by the numbers of full genomes and HN gene sequences generated in this study, respectively. The world map was generated using the open-source Quantum Geographic Information System (QGIS), version 3.38, with raster map data from Natural Earth. (*B*) Pie charts show the proportion of newly generated full genomes for each of the four HPIV types relative to all corresponding sequences available in GenBank, highlighting the expansion of available genomic datasets through this study. (*C*) Proportion of newly generated HN gene sequences for four HPIV types in this study relative to all corresponding HN sequences available in GenBank, highlighting the expansion of available HN sequences through this study.

**Appendix Figure S2.**
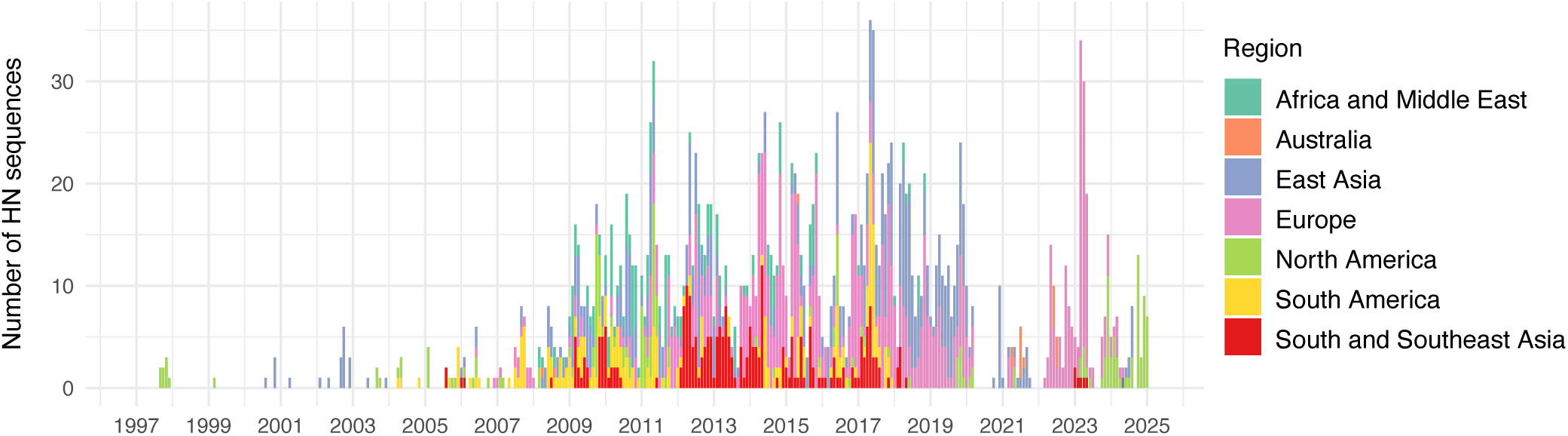
Weekly number of HPIV detections by geographic regions. Stacked coloured bars show the number of available hemagglutinin-neuraminidase (HN) sequences from each geographic region.

**Appendix Figure S3.**
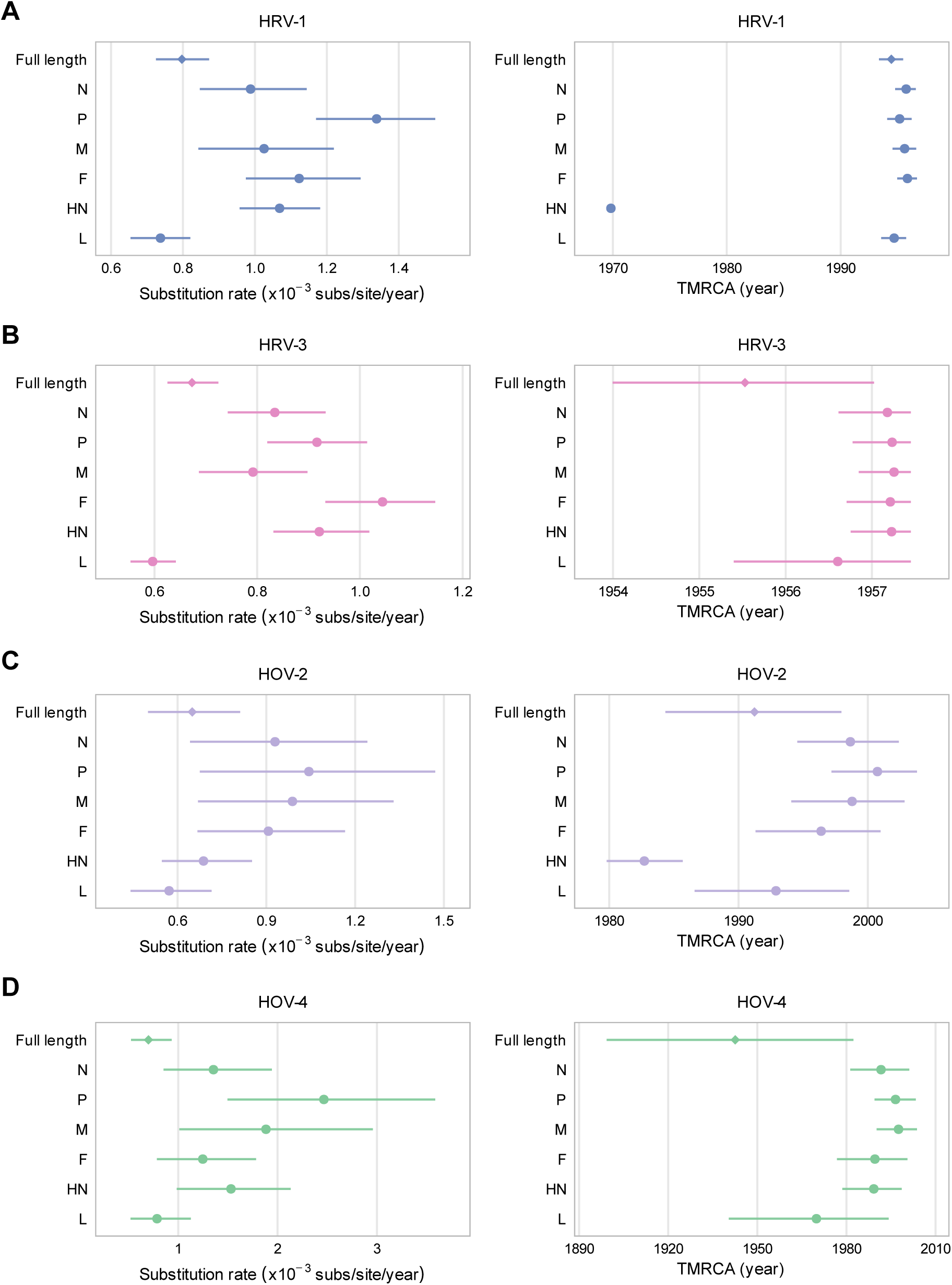
Comparison of evolutionary rates and estimated time to the most recent common ancestor (TMRCA) among HPIV types based on complete genome and individual gene sequences. Substitution rates (nucleotide substitutions per site per year) and TMRCA estimates were simultaneously inferred for each virus type using Bayesian skyride analyses implemented in BEAST. Dots indicate mean estimates and lines indicate 95% highest posterior density (HPD) intervals for (*A*) HRV-1, (*B*) HRV-3, (*C*) HOV-2, and (*D*) HOV-4.

**Appendix Figure S4.**
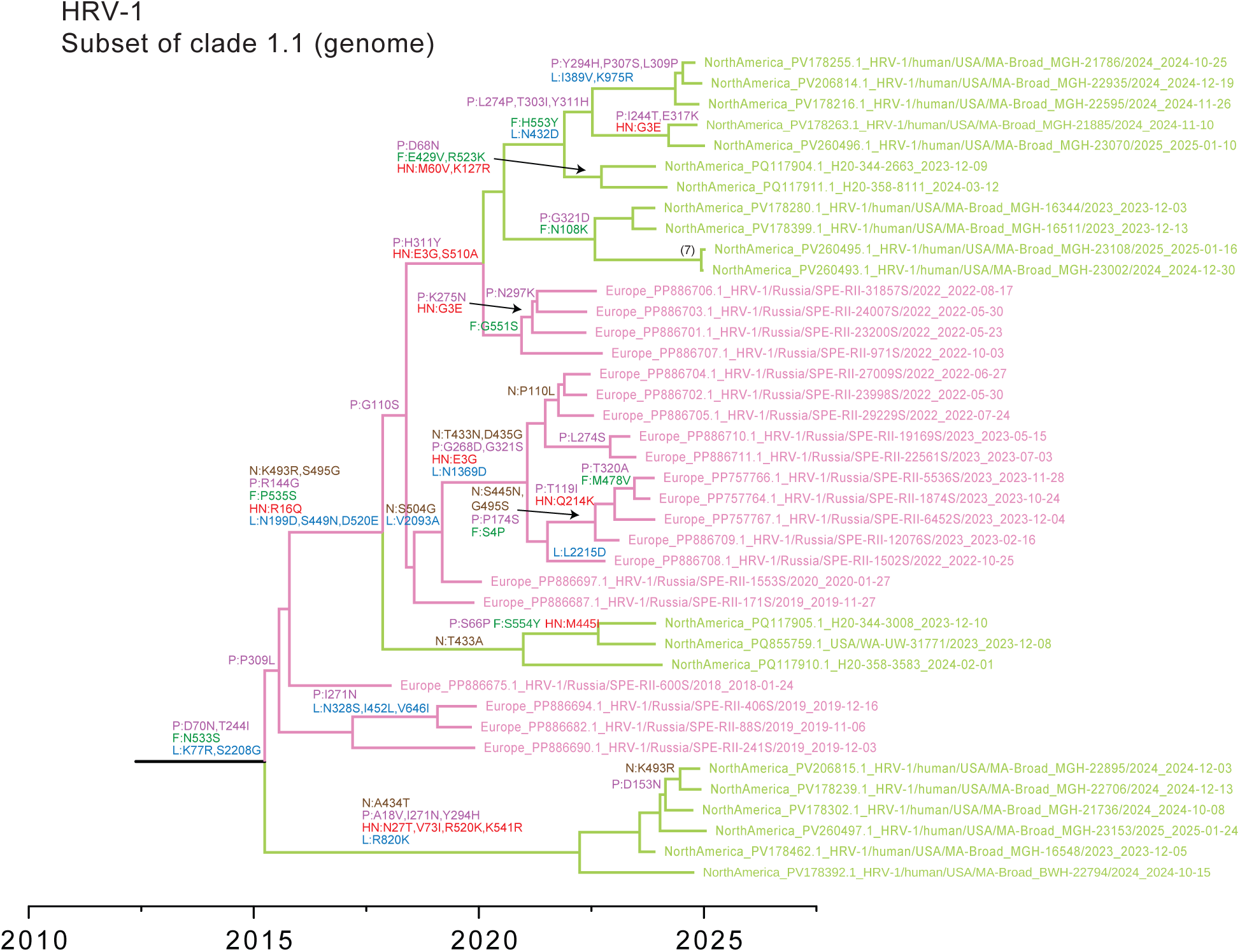
Subset of human respirovirus 1 (HRV-1) evolution showing amino acid substitutions within the cluster indicated by the blue box in Fig 2A. Time-scaled phylogeny of HRV-1 based on complete genomes. Pink and green branches denote sequences from Europe and North America, respectively.

**Appendix Figure S5.**
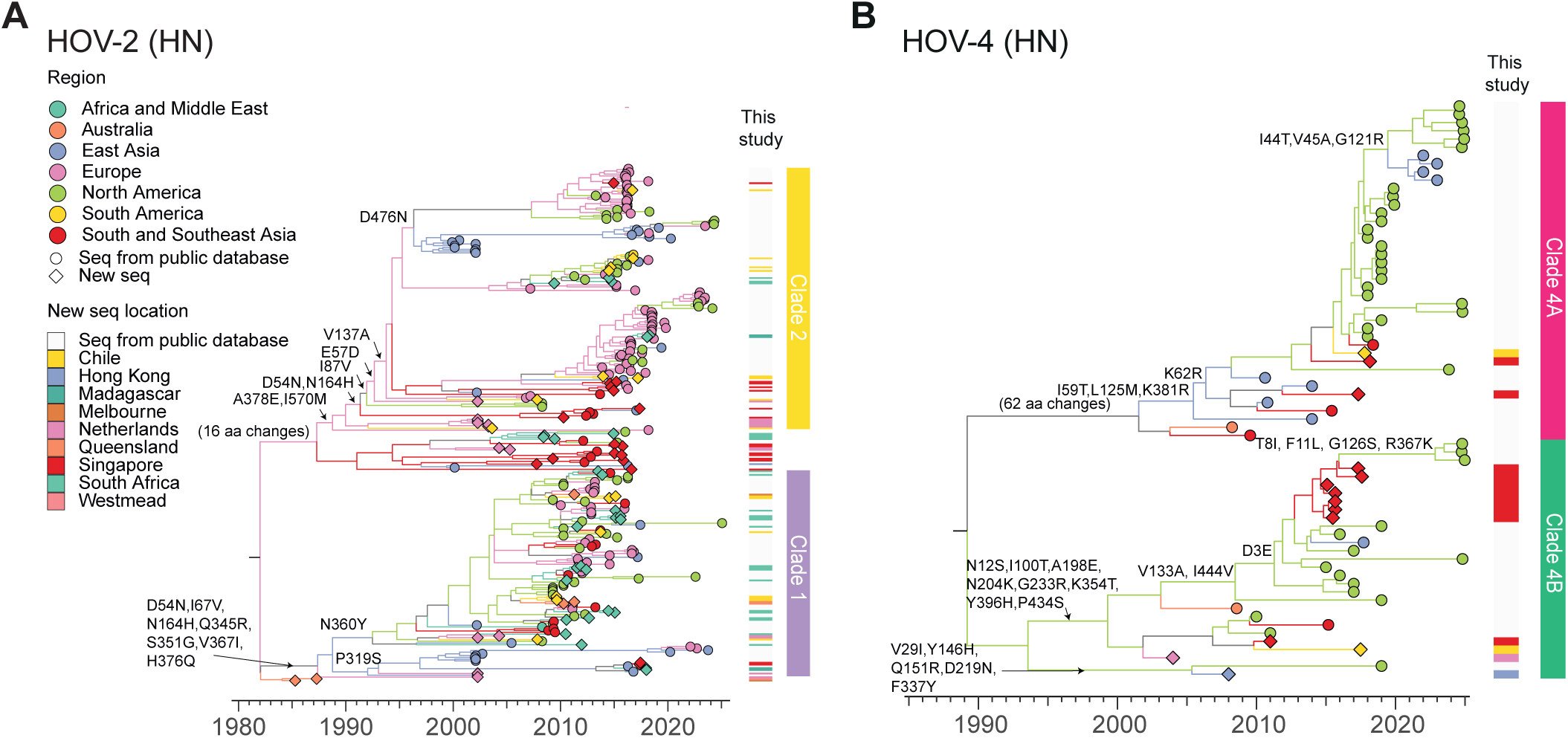
Time-scaled phylogeny of the HN gene of human orthorubulavirus 2 (HOV-2) and 4 (HOV-4). Maximum clade credibility (MCC) trees show the evolution of two co-circulating clades for each virus type. Coloured branches denote the geographic locations. Diamonds at the tips denote sequences generated in this study, while open circles represent sequences obtained from NCBI GenBank. Amino acid substitutions were mapped at major nodes. The geographic locations of newly generated sequences and clade numbering are indicated by bars.

**Appendix Figure S6.**
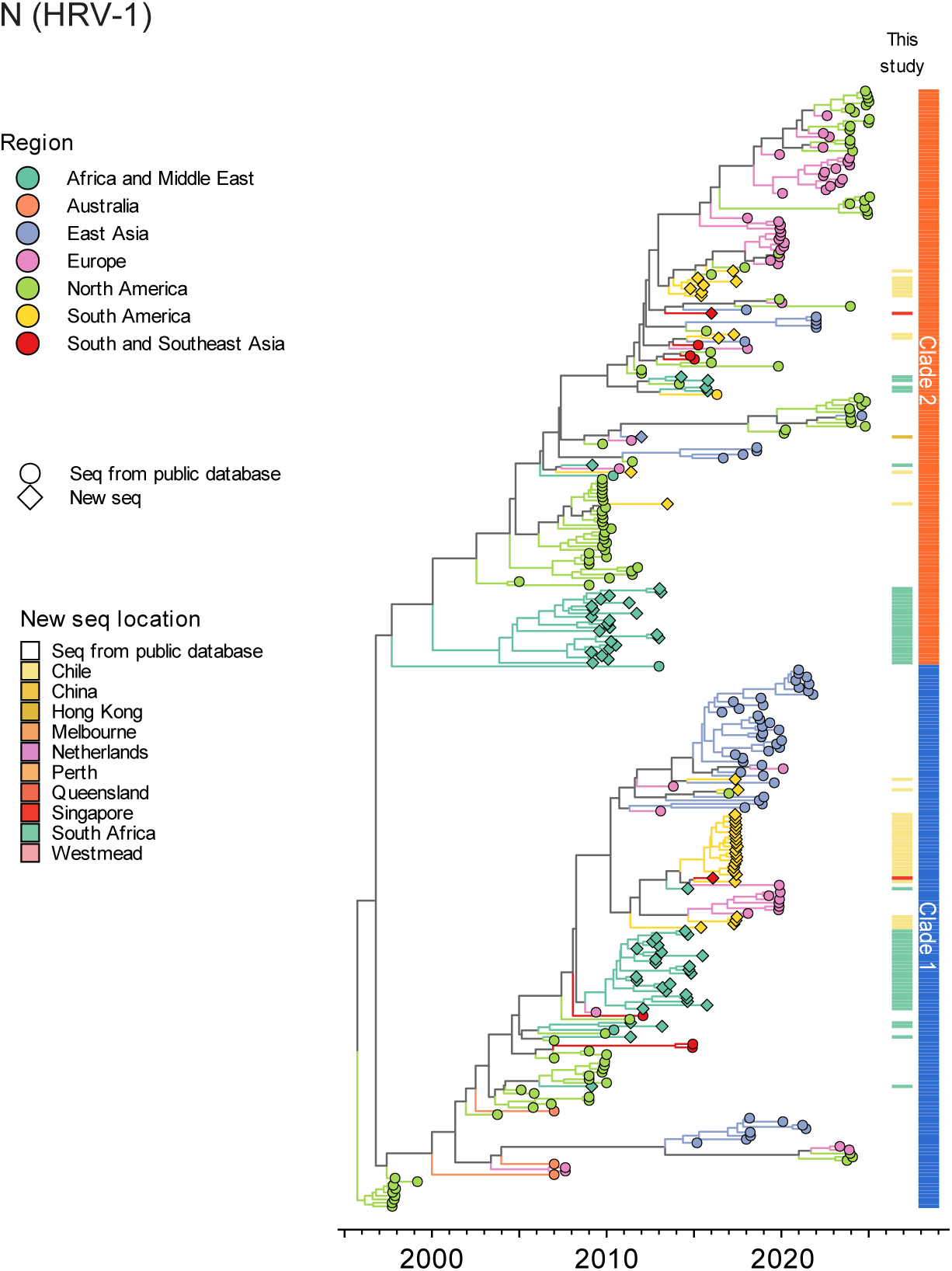
Time-scaled phylogeny of the N gene of HRV-1. Coloured branches denote the geographic locations. Diamonds at the tips denote sequences generated in this study, while open circles represent sequences obtained from NCBI GenBank. Amino acid substitutions were mapped at major nodes. The geographic locations of newly generated sequences and clade numbering are indicated by bars.

**Appendix Figure S7.**
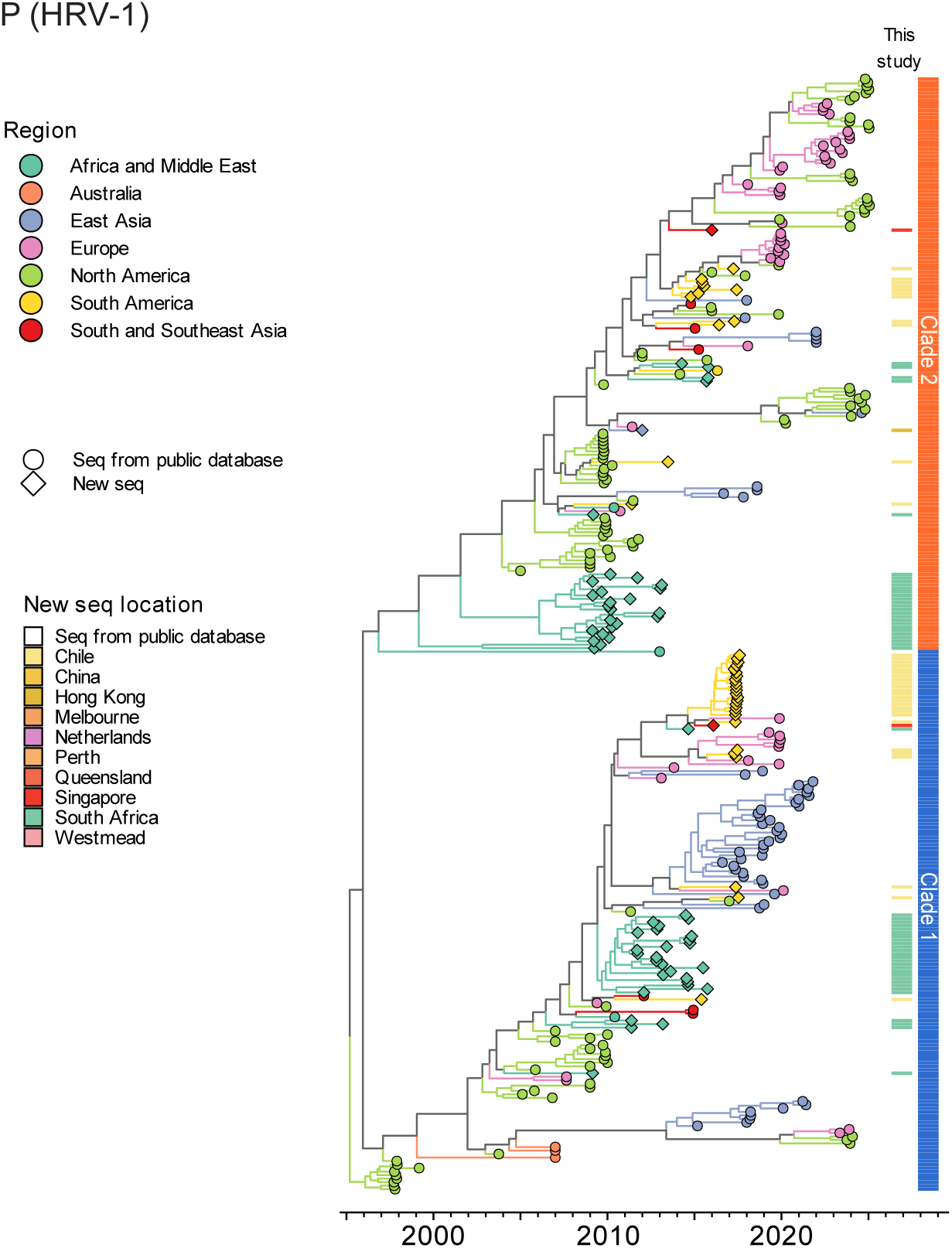
Time-scaled phylogeny of the P gene of HRV-1. Coloured branches denote the geographic locations. Diamonds at the tips denote sequences generated in this study, while open circles represent sequences obtained from NCBI GenBank. Amino acid substitutions were mapped at major nodes. The geographic locations of newly generated sequences and clade numbering are indicated by bars.

**Appendix Figure S8.**
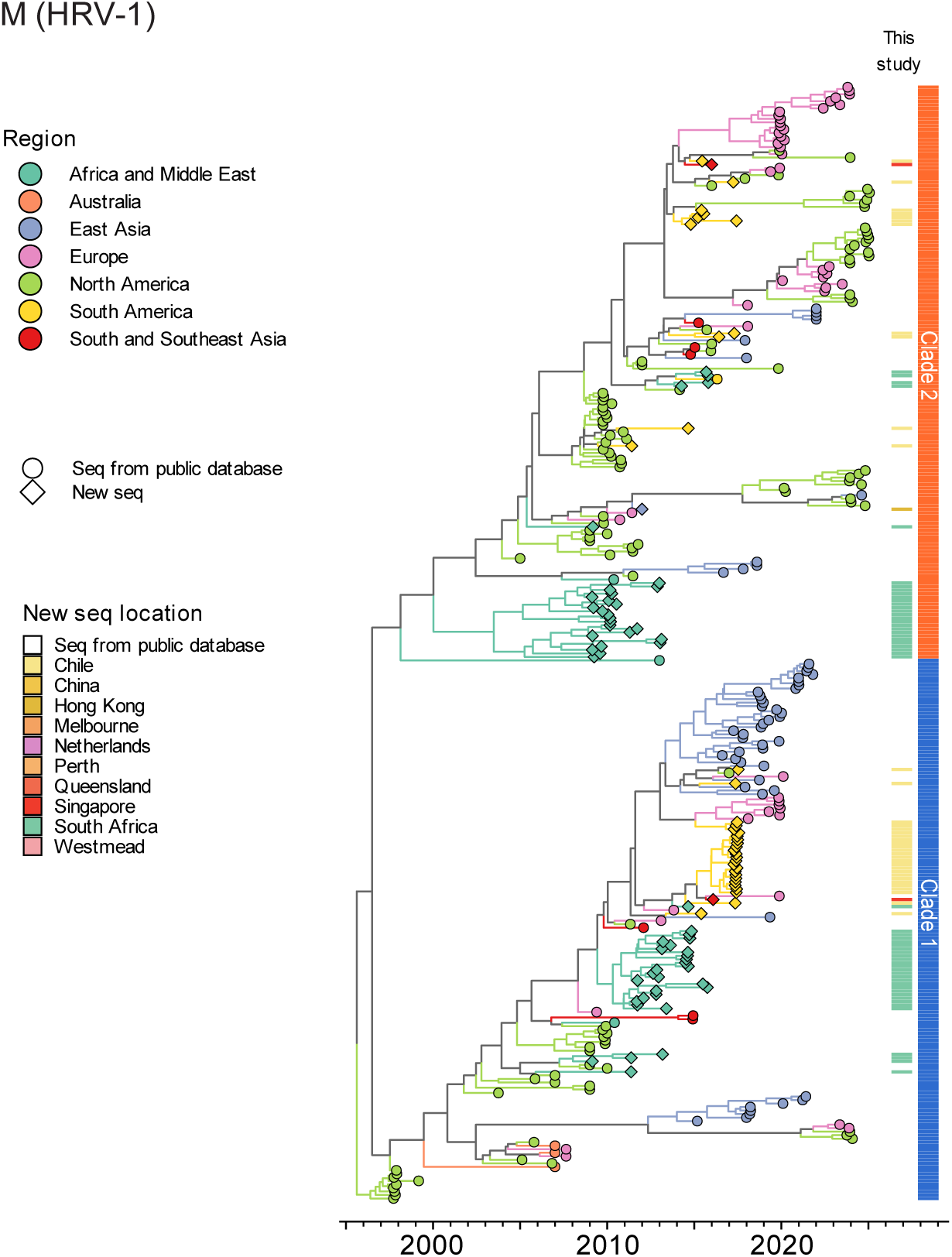
Time-scaled phylogeny of the M gene of HRV-1. Coloured branches denote the geographic locations. Diamonds at the tips denote sequences generated in this study, while open circles represent sequences obtained from NCBI GenBank. Amino acid substitutions were mapped at major nodes. The geographic locations of newly generatedsequences and clade numbering are indicated by bars.

**Appendix Figure S9.**
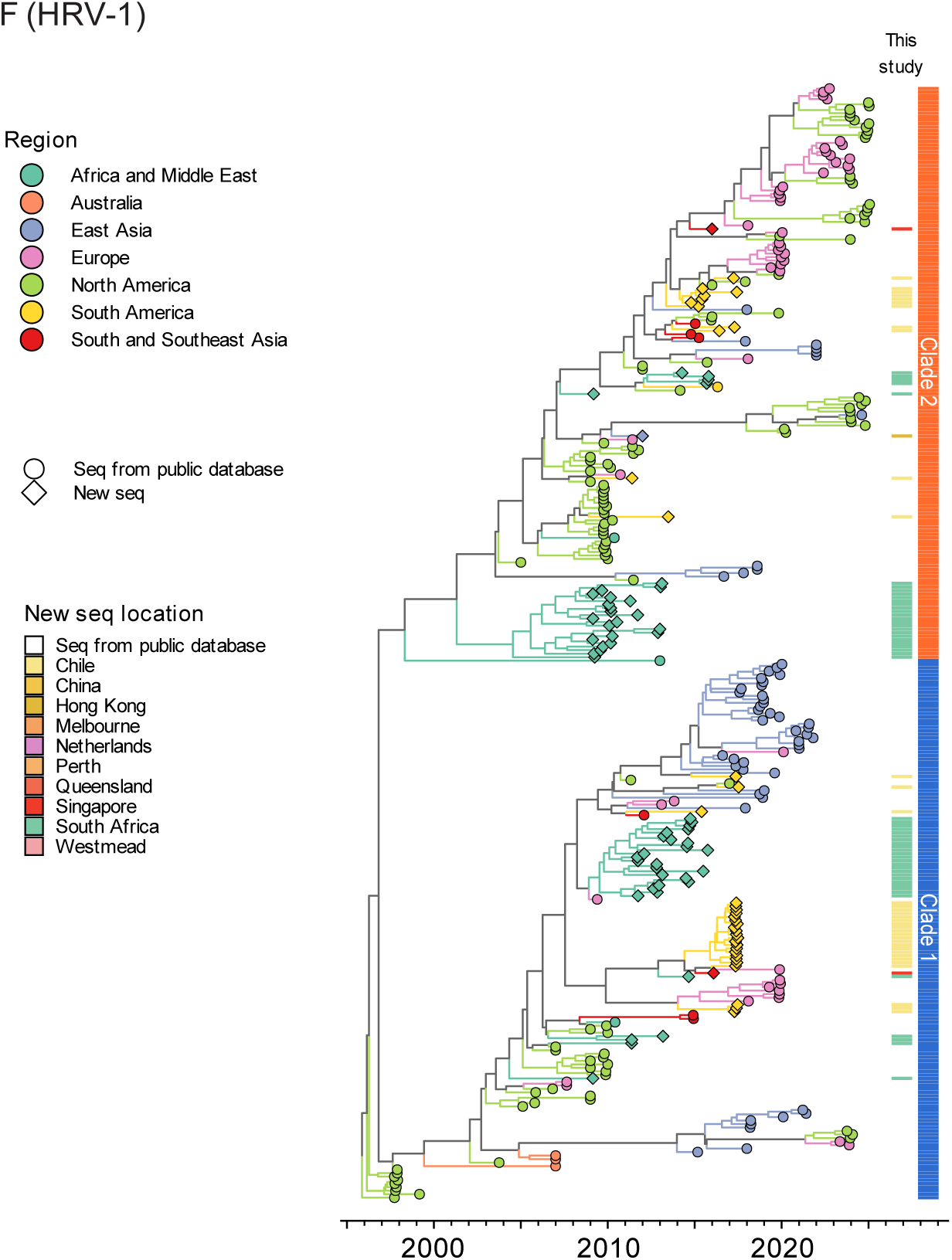
Time-scaled phylogeny of the F gene of HRV-1. Coloured branches denote the geographic locations. Diamonds at the tipsdenote sequences generated in this study, while open circles represent sequences obtained from NCBI GenBank. Amino acid substitutions were mapped at major nodes. The geographic locations of newly generatedsequences and clade numbering are indicated by bars.

**Appendix Figure S10.**
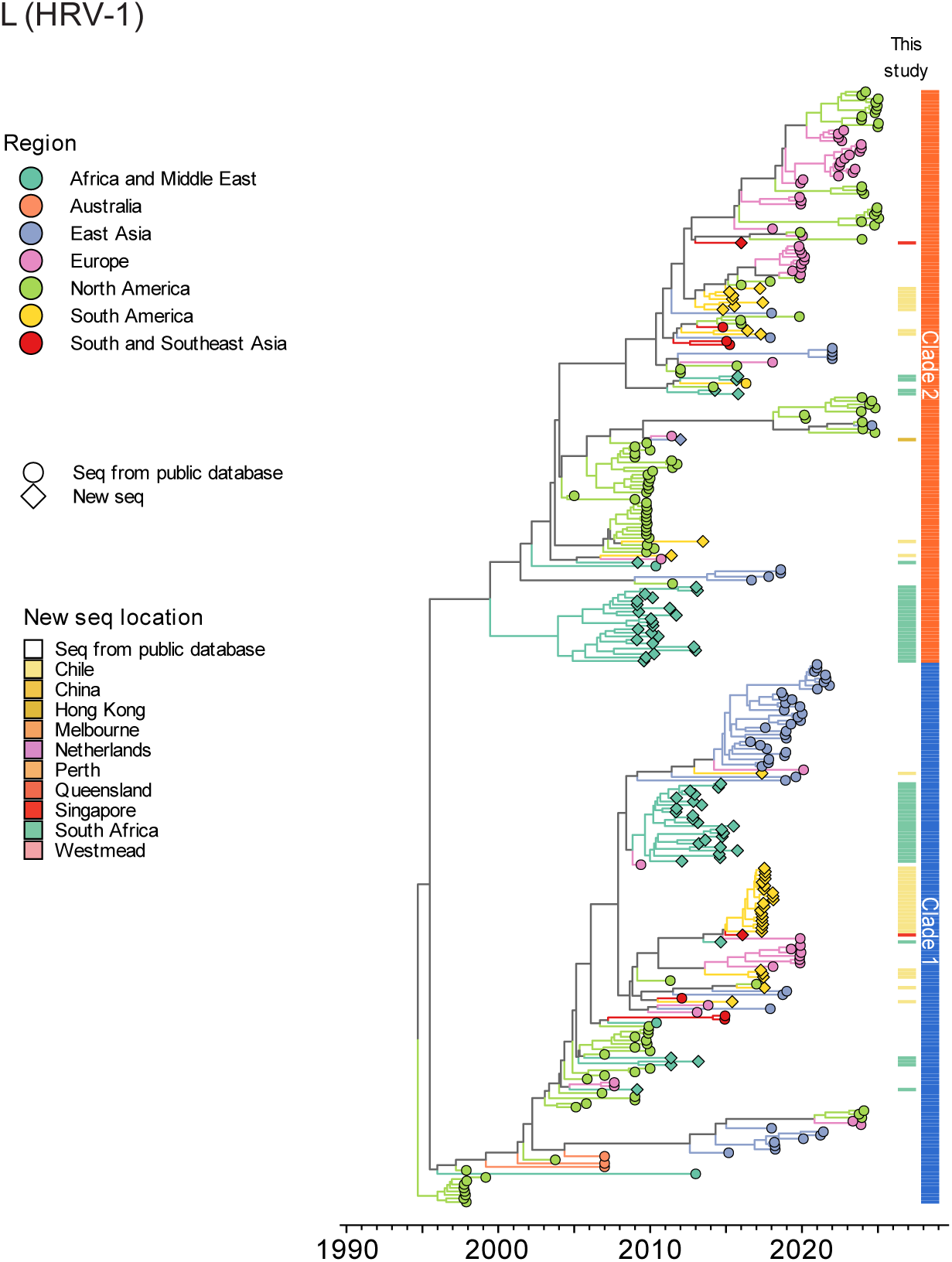
Time-scaled phylogeny of the L gene of HRV-1. Coloured branches denote the geographic locations. Diamonds at the tips denote sequences generated in this study, while open circles represent sequences obtained from NCBI GenBank. Amino acid substitutions were mapped at major nodes. The geographic locations of newly generated sequences and clade numbering are indicated by bars.

**Appendix Figure S11.**
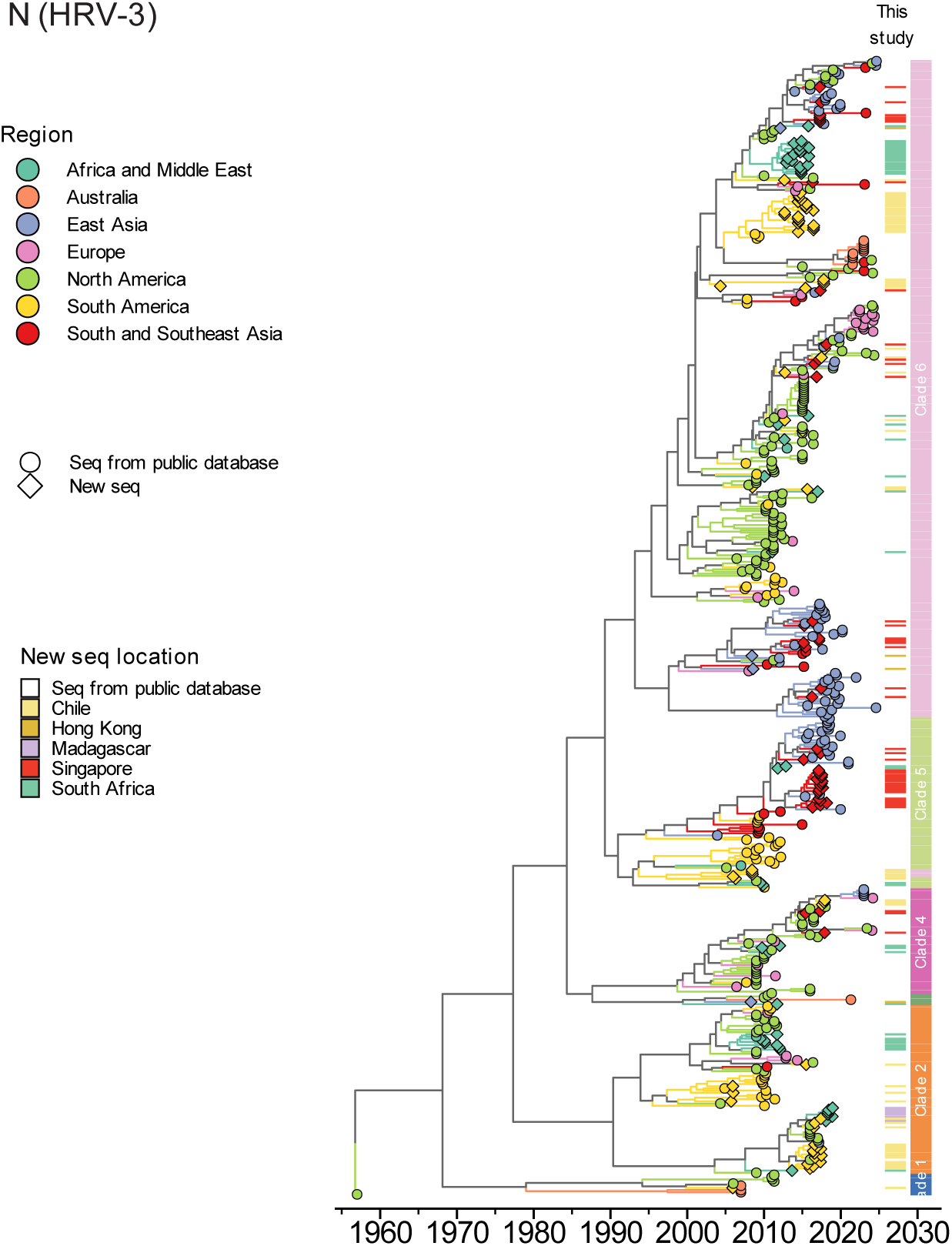
Time-scaled phylogeny of the N gene of HRV-3. Coloured branches denote the geographic locations. Diamonds at the tips denote sequences generated in this study, while open circles represent sequences obtained from NCBI GenBank. Amino acid substitutions were mapped at major nodes. The geographic locations of newly generated sequences and clade numbering are indicated by bars.

**Appendix Figure S12.**
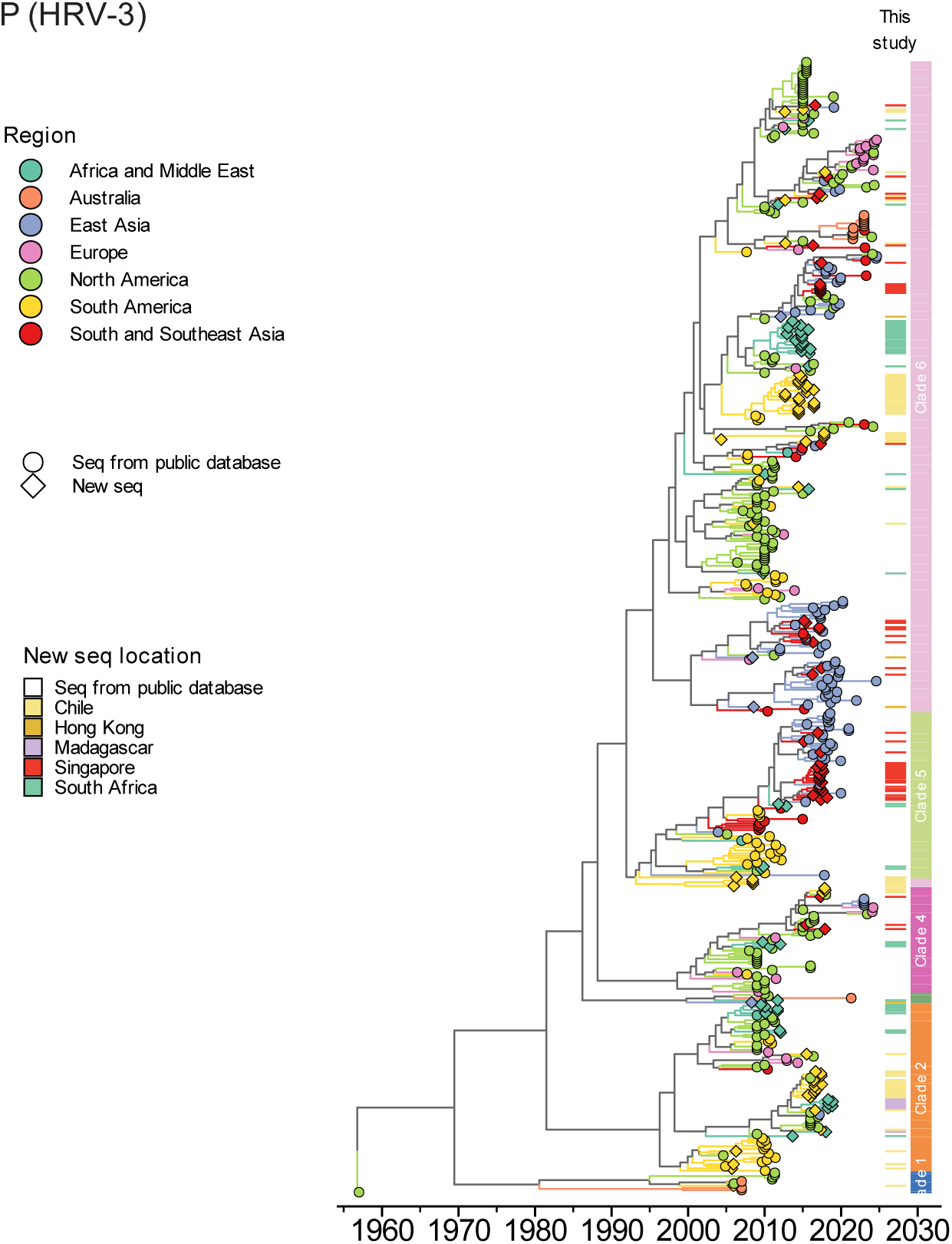
Time-scaled phylogeny of the P gene of HRV-3. Coloured branches denote the geographic locations. Diamonds at the tips denote sequences generated in this study, while open circles represent sequences obtained from NCBI GenBank. Amino acid substitutions were mapped at major nodes. The geographic locations of newly generated sequences and clade numbering are indicated by bars.

**Appendix Figure S13.**
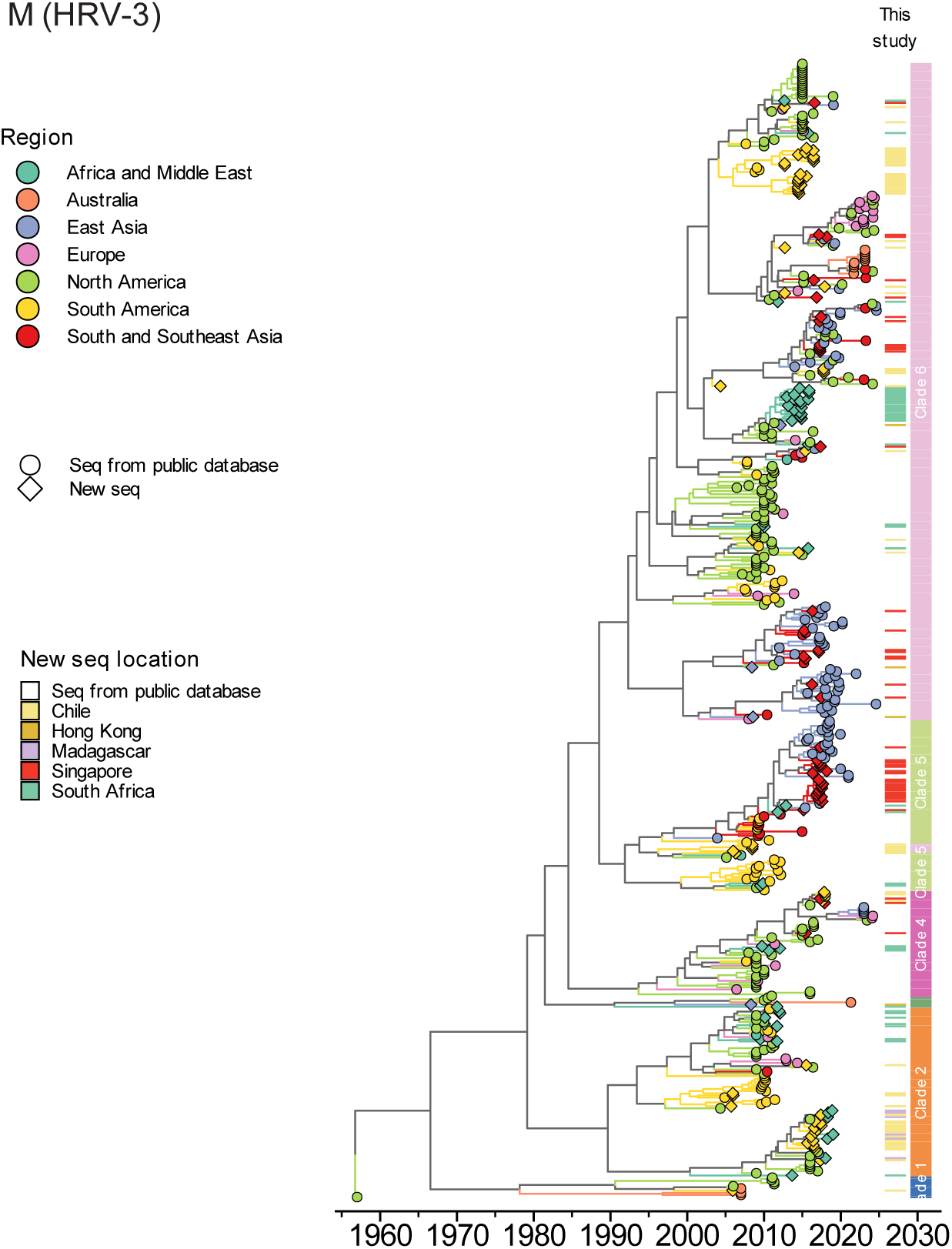
Time-scaled phylogeny of the M gene of HRV-3. Coloured branches denote the geographic locations. Diamonds at the tips denote sequences generated in this study, while open circles represent sequences obtained from NCBI GenBank. Amino acid substitutions were mapped at major nodes. The geographic locations of newly generatedsequences and clade numbering are indicated by bars.

**Appendix Figure S14.**
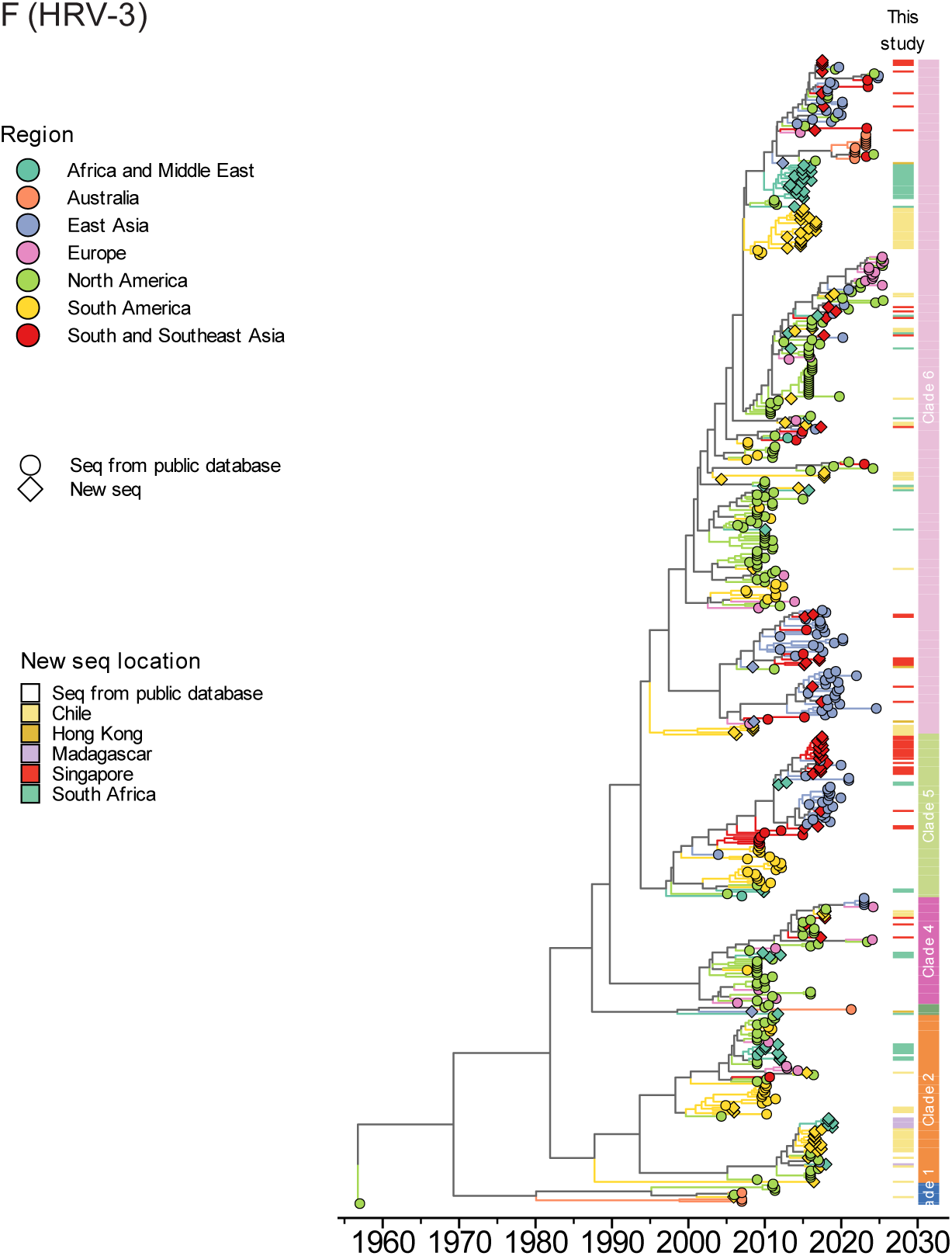
Time-scaled phylogeny of the F gene of HRV-3. Coloured branches denote the geographic locations. Diamonds at the tips denote sequences generated in this study, while open circles represent sequences obtained from NCBI GenBank. Amino acid substitutions were mapped at major nodes. The geographic locations of newly generated sequences and clade numbering are indicated by bars.

**Appendix Figure S15.**
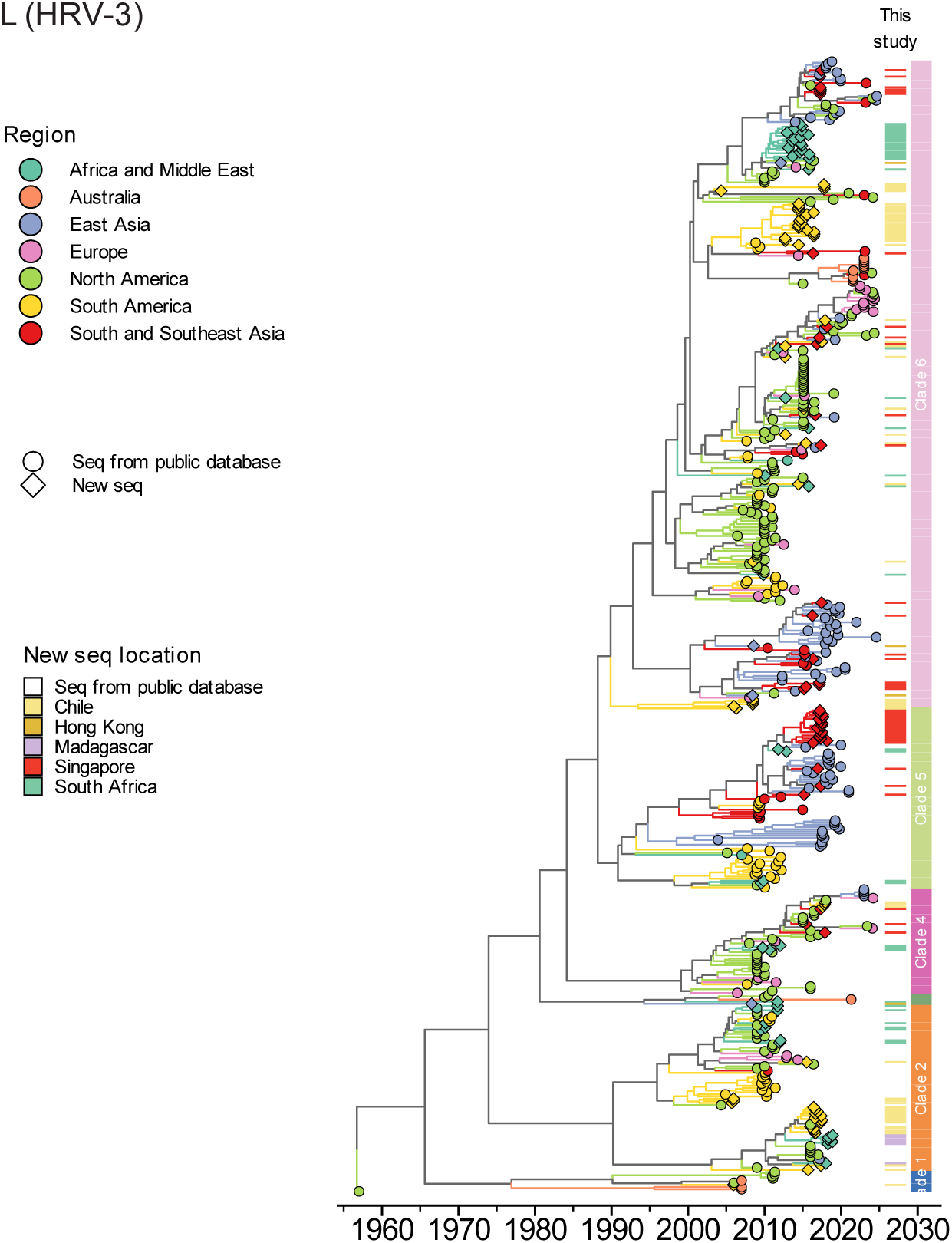
Time-scaled phylogeny of the L gene of HRV-3. Coloured branches denote the geographic locations. Diamonds at the tips denote sequences generated in this study, while open circles represent sequences obtained from NCBI GenBank. Amino acid substitutions were mapped at major nodes. The geographic locations of newly generated sequences and clade numbering are indicated by bars.

**Appendix Figure S16.**
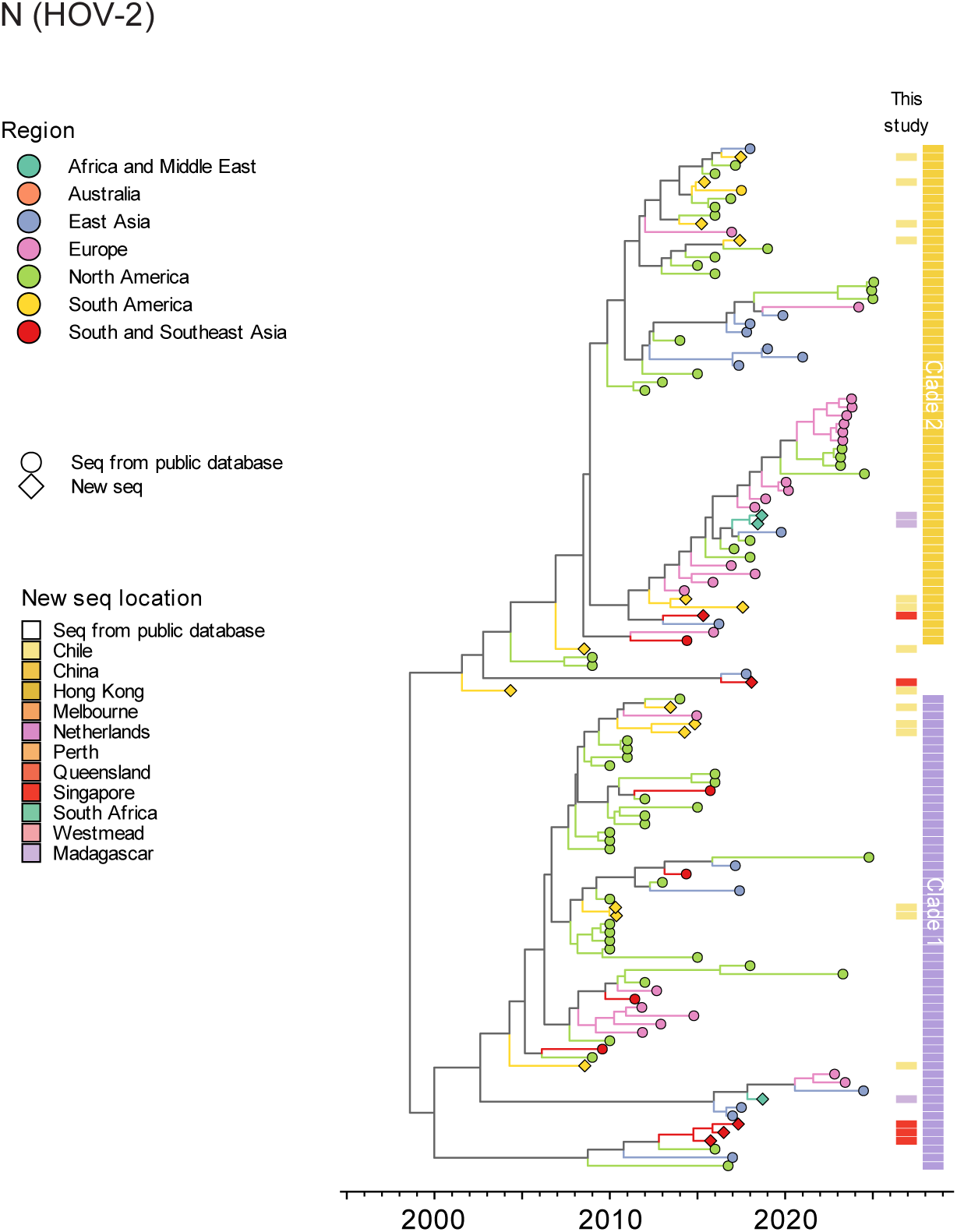
Time-scaled phylogeny of the N gene of HOV-2. Coloured branches denote the geographic locations. Diamonds at the tips denote sequences generated in this study, while open circles represent sequences obtained from NCBI GenBank. Amino acid substitutions were mapped at major nodes. The geographic locations of newly generated sequences and clade numbering are indicated by bars.

**Appendix Figure S17.**
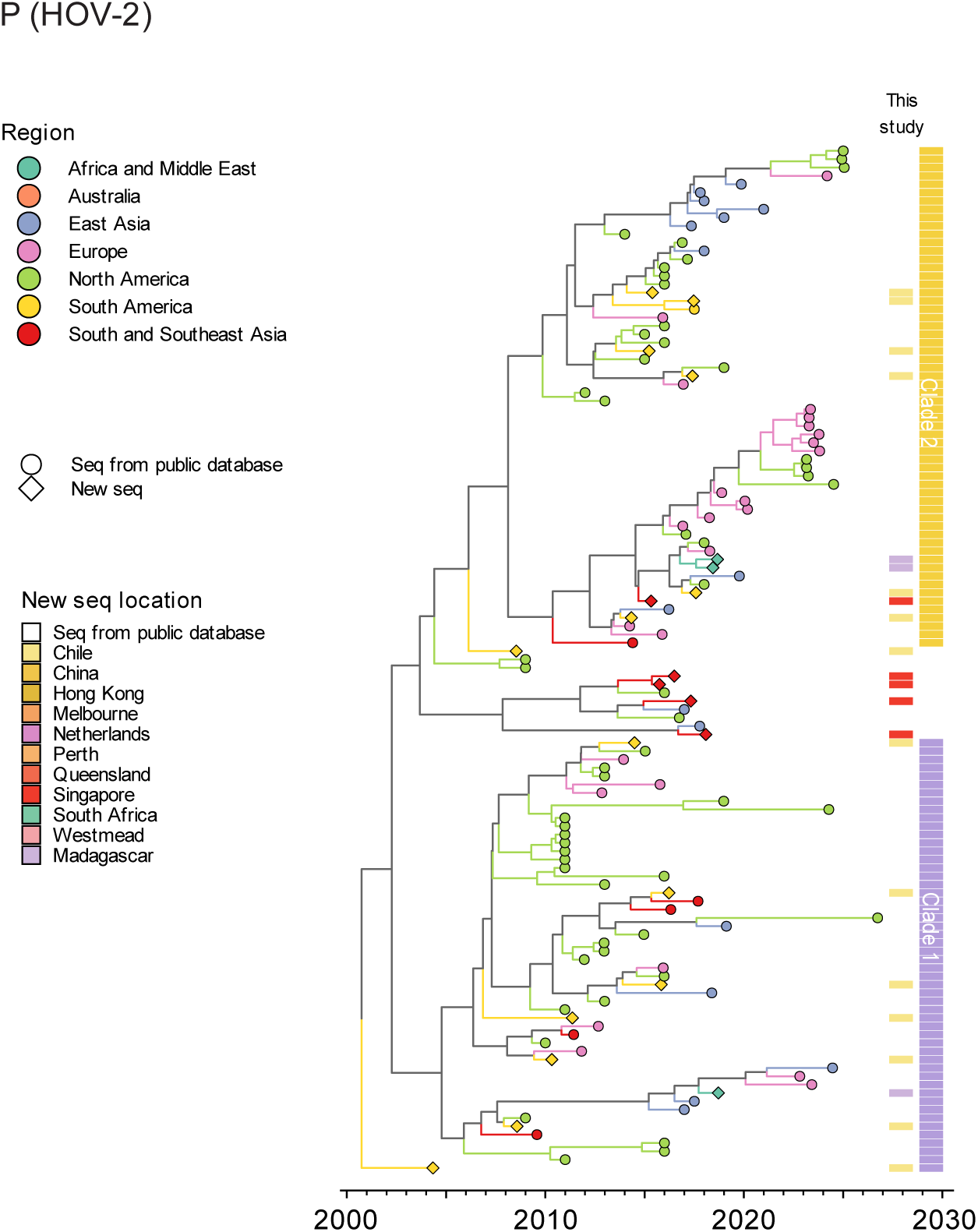
Time-scaled phylogeny of the P gene of HOV-2. Coloured branches denote the geographic locations. Diamonds at the tips denote sequences generated in this study, while open circles represent sequences obtained from NCBI GenBank. Amino acid substitutions were mapped at major nodes. The geographic locations of newly generated sequences and clade numbering are indicated by bars.

**Appendix Figure S18.**
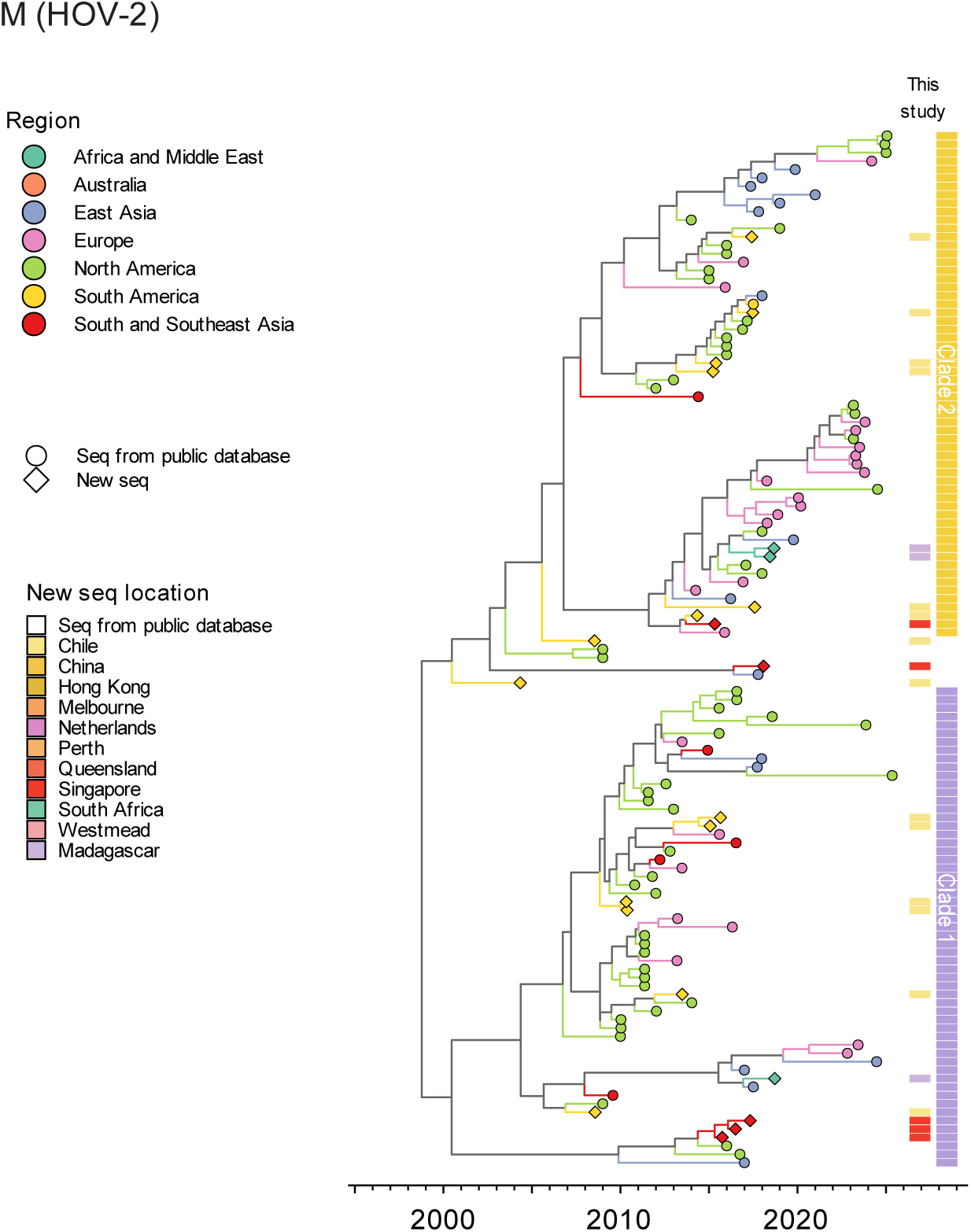
Time-scaled phylogeny of the M gene of HOV-2. Coloured branches denote the geographic locations. Diamonds at the tips denote sequences generated in this study, while open circles represent sequences obtained from NCBI GenBank. Amino acid substitutions were mapped at major nodes. The geographic locations of newly generated sequences and clade numbering are indicated by bars.

**Appendix Figure S19.**
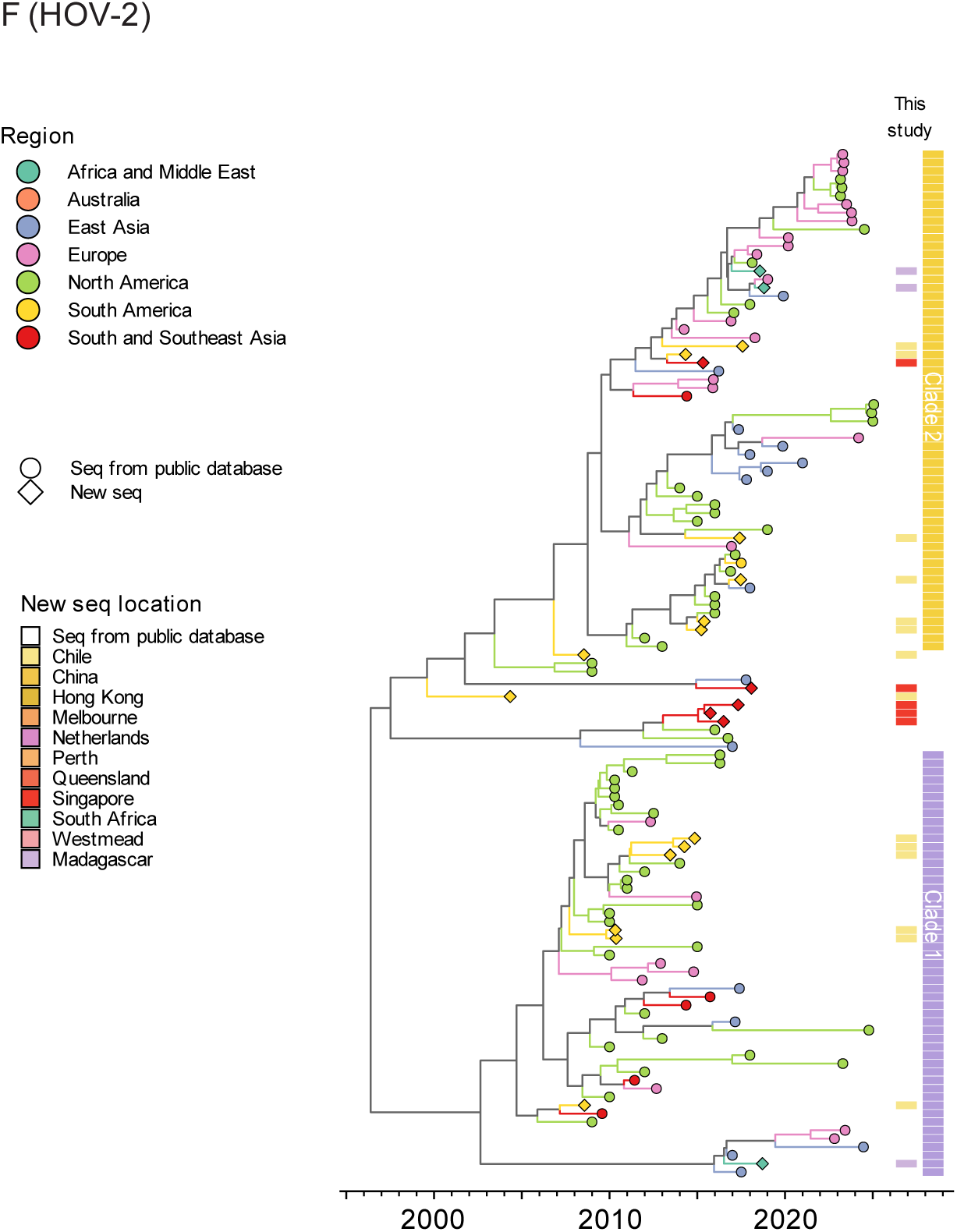
Time-scaled phylogeny of the F gene of HOV-2. Coloured branches denote the geographic locations. Diamonds at the tips denote sequences generated in this study, while open circles represent sequences obtained from NCBI GenBank. Amino acid substitutions were mapped at major nodes. The geographic locations of newly generated sequences and clade numbering are indicated by bars.

**Appendix Figure S20.**
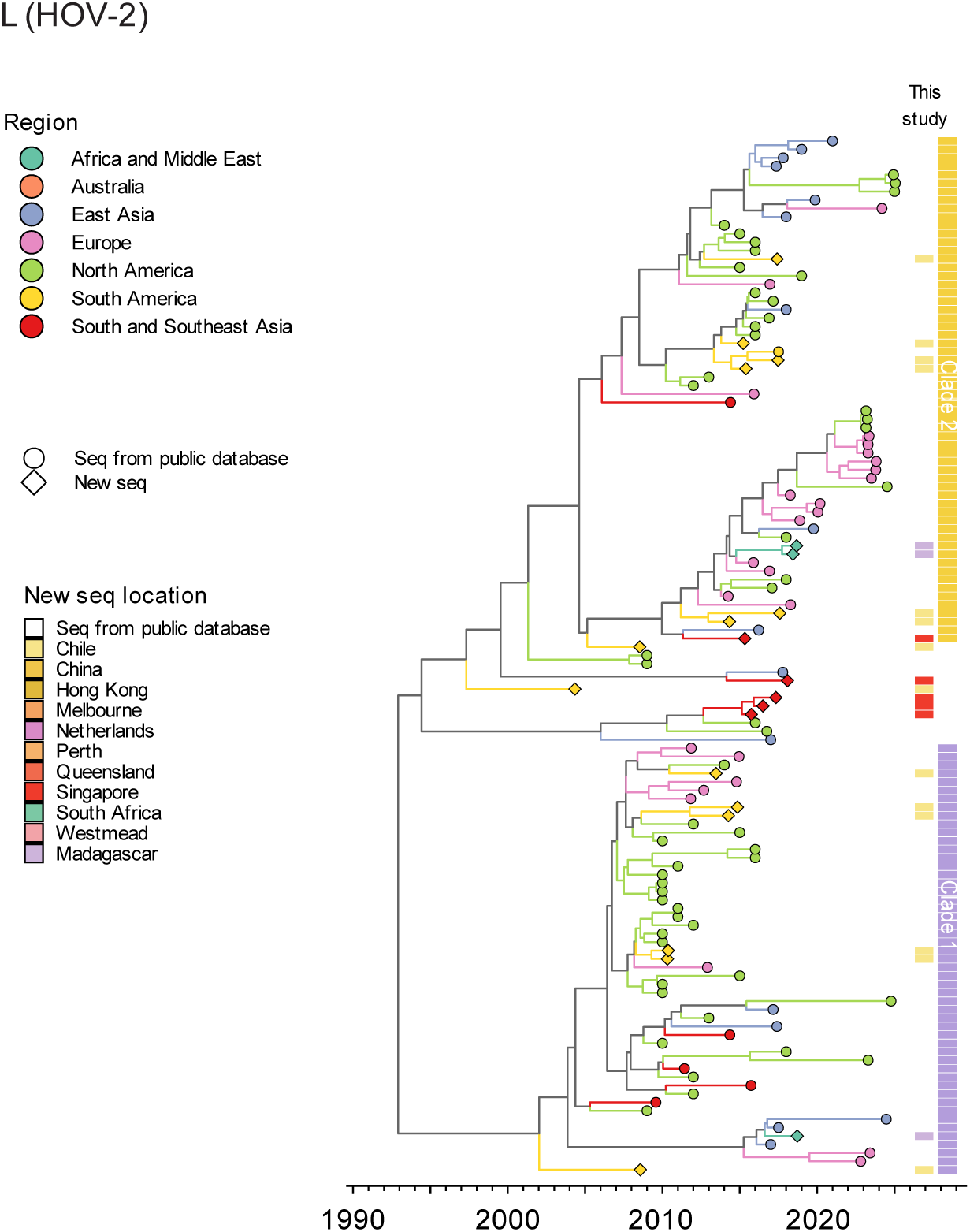
Time-scaled phylogeny of the L gene of HOV-2. Coloured branches denote the geographic locations. Diamonds at the tips denote sequences generated in this study, while open circles represent sequences obtained from NCBI GenBank. Amino acid substitutions were mapped at major nodes. The geographic locations of newly generated sequences and clade numbering are indicated by bars.

**Appendix Figure S21.**
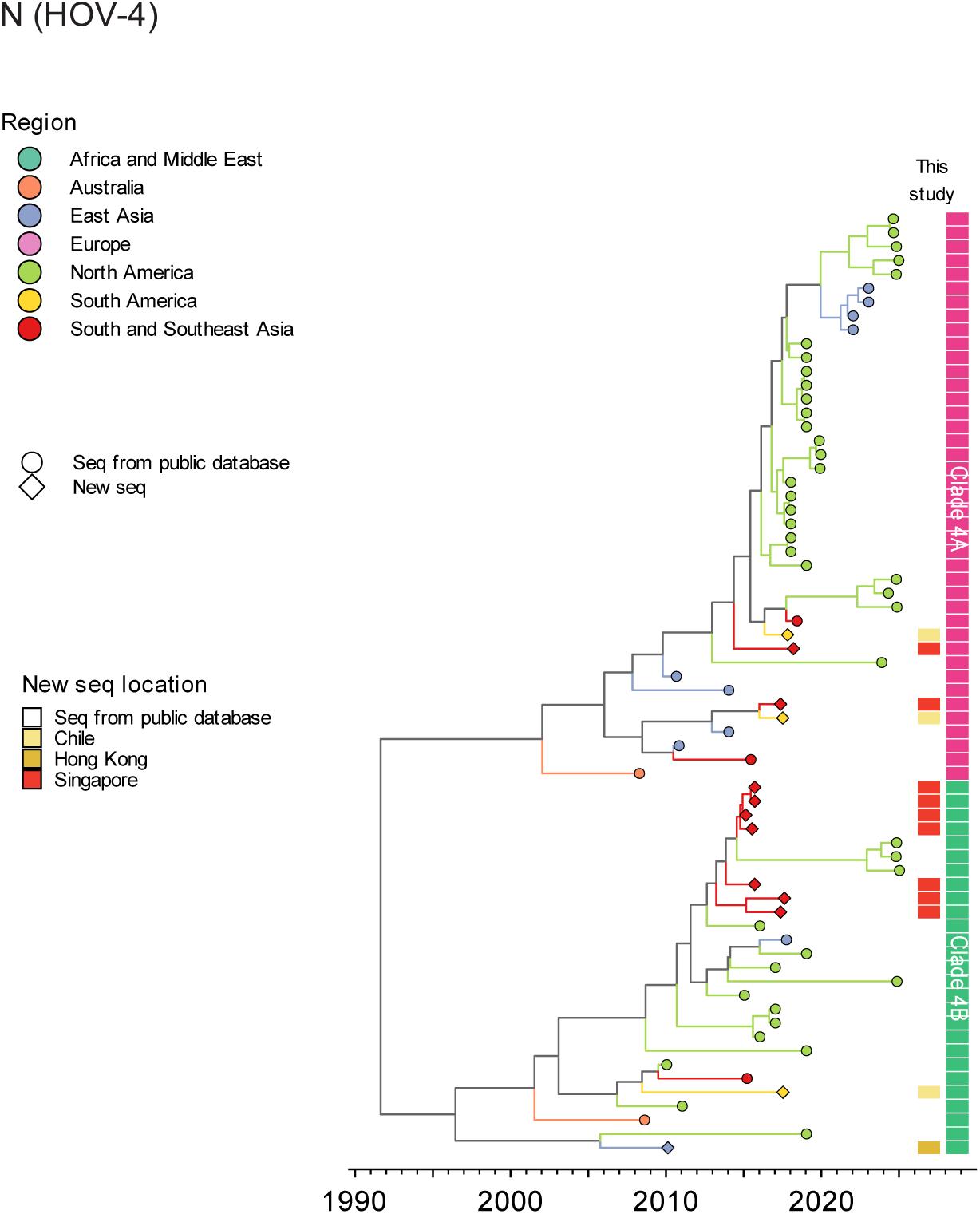
Time-scaled phylogeny of the N gene of HOV-4. Coloured branches denote the geographic locations. Diamonds at the tips denote sequences generated in this study, while open circles represent sequences obtained from NCBI GenBank. Amino acid substitutions were mapped at major nodes. The geographic locations of newly generated sequences and clade numbering are indicated by bars.

**Appendix Figure S22.**
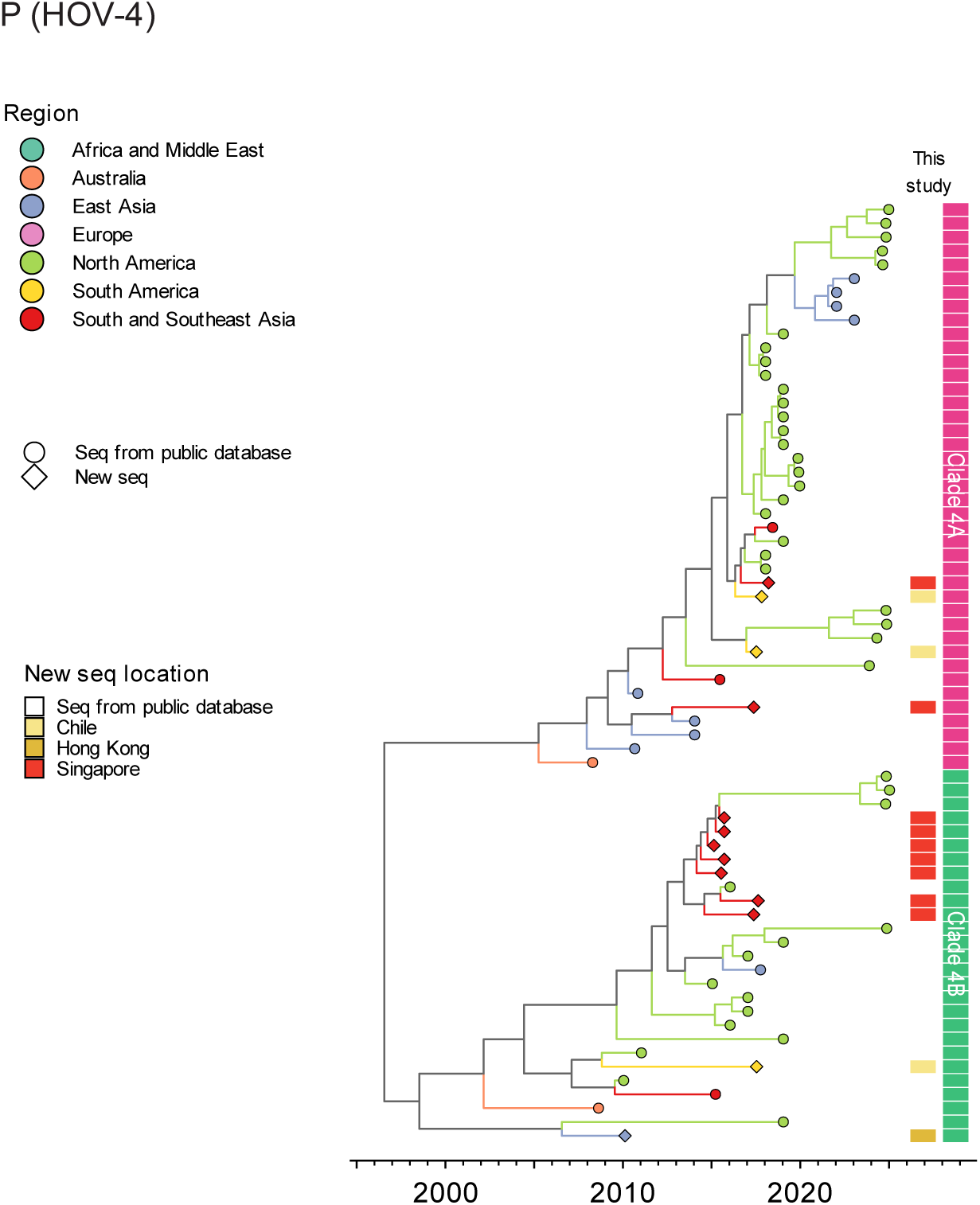
Time-scaled phylogeny of the P gene of HOV-4. Coloured branches denote the geographic locations. Diamonds at the tips denote sequences generated in this study, while open circles represent sequences obtained from NCBI GenBank. Amino acid substitutions were mapped at major nodes. The geographic locations of newly generated sequences and clade numbering are indicated by bars.

**Appendix Figure S23.**
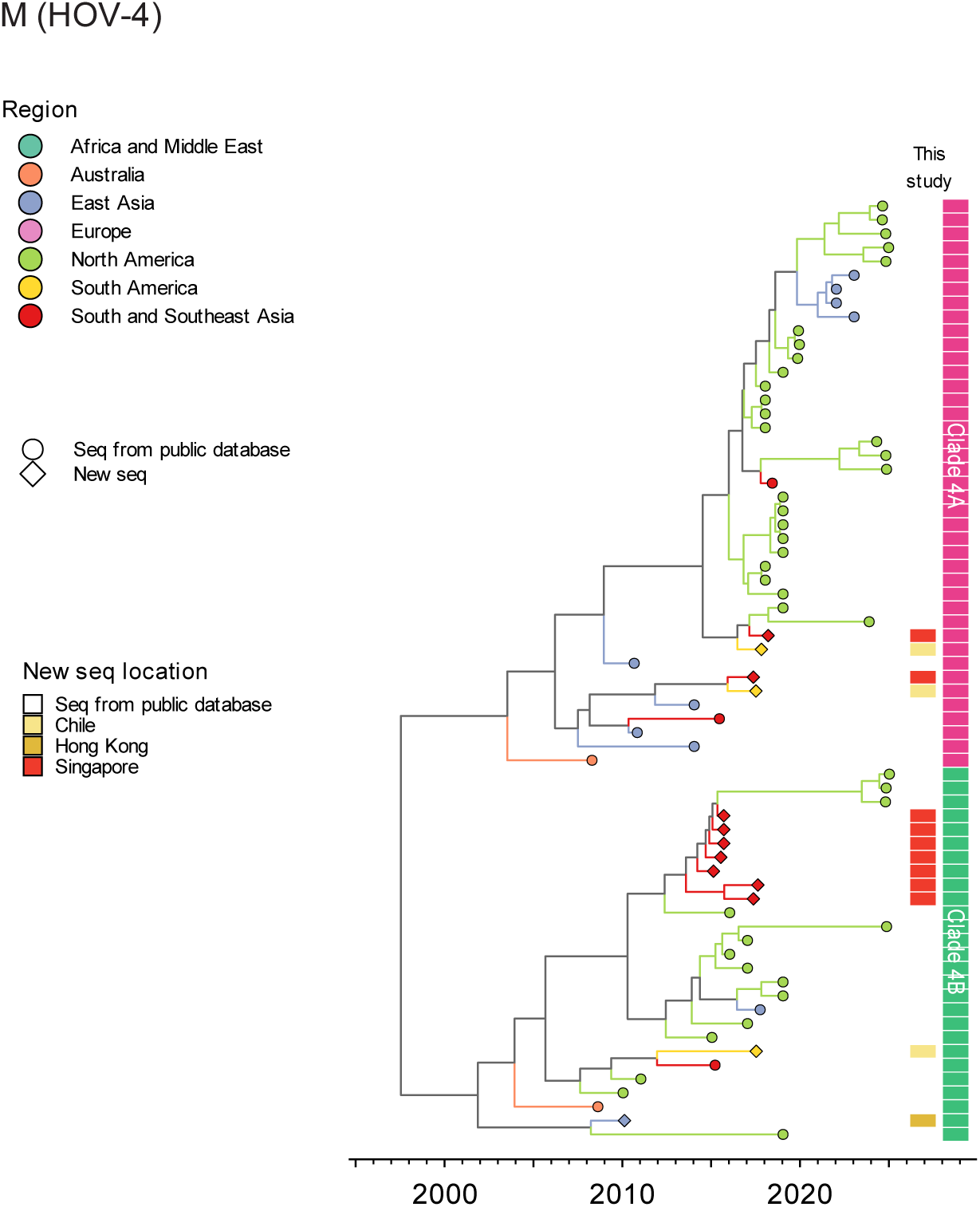
Time-scaled phylogeny of the M gene of HOV-4. Coloured branches denote the geographic locations. Diamonds at the tips denote sequences generated in this study, while open circles represent sequences obtained from NCBI GenBank. Amino acid substitutions were mapped at major nodes. The geographic locations of newly generated sequences and clade numbering are indicated by bars.

**Appendix Figure S24.**
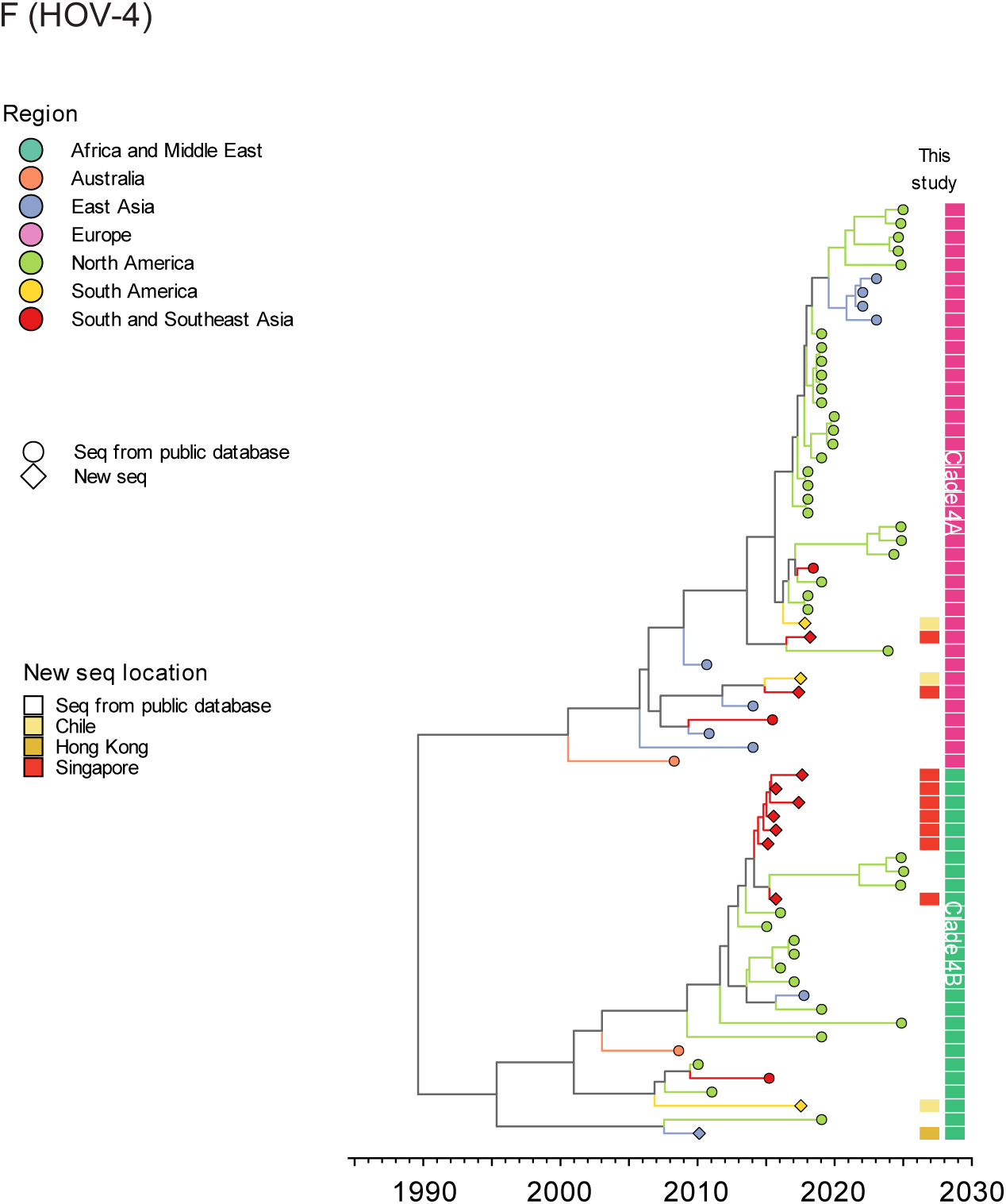
Time-scaled phylogeny of the F gene of HOV-4. Coloured branches denote the geographic locations. Diamonds at the tips denote sequences generated in this study, while open circles represent sequences obtained from NCBI GenBank. Amino acid substitutions were mapped at major nodes. The geographic locations of newly generated sequences and clade numbering are indicated by bars.

**Appendix Figure S25.**
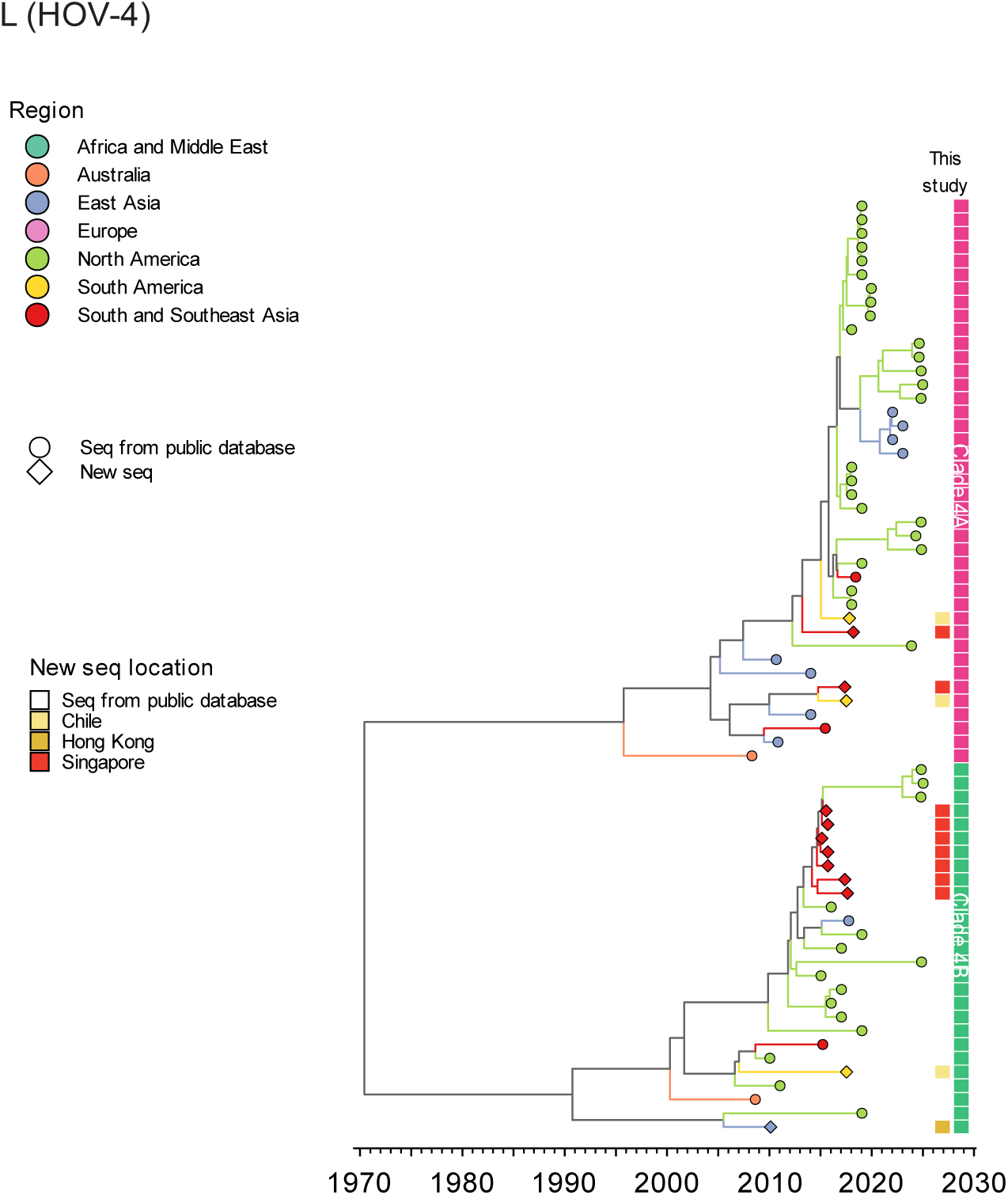
Time-scaled phylogeny of the L gene of HOV-4. Coloured branches denote the geographic locations. Diamonds at the tips denote sequences generated in this study, while open circles represent sequences obtained from NCBI GenBank. Amino acid substitutions were mapped at major nodes. The geographic locations of newly generated sequences and clade numbering are indicated by bars.

**Appendix Figure S26.**
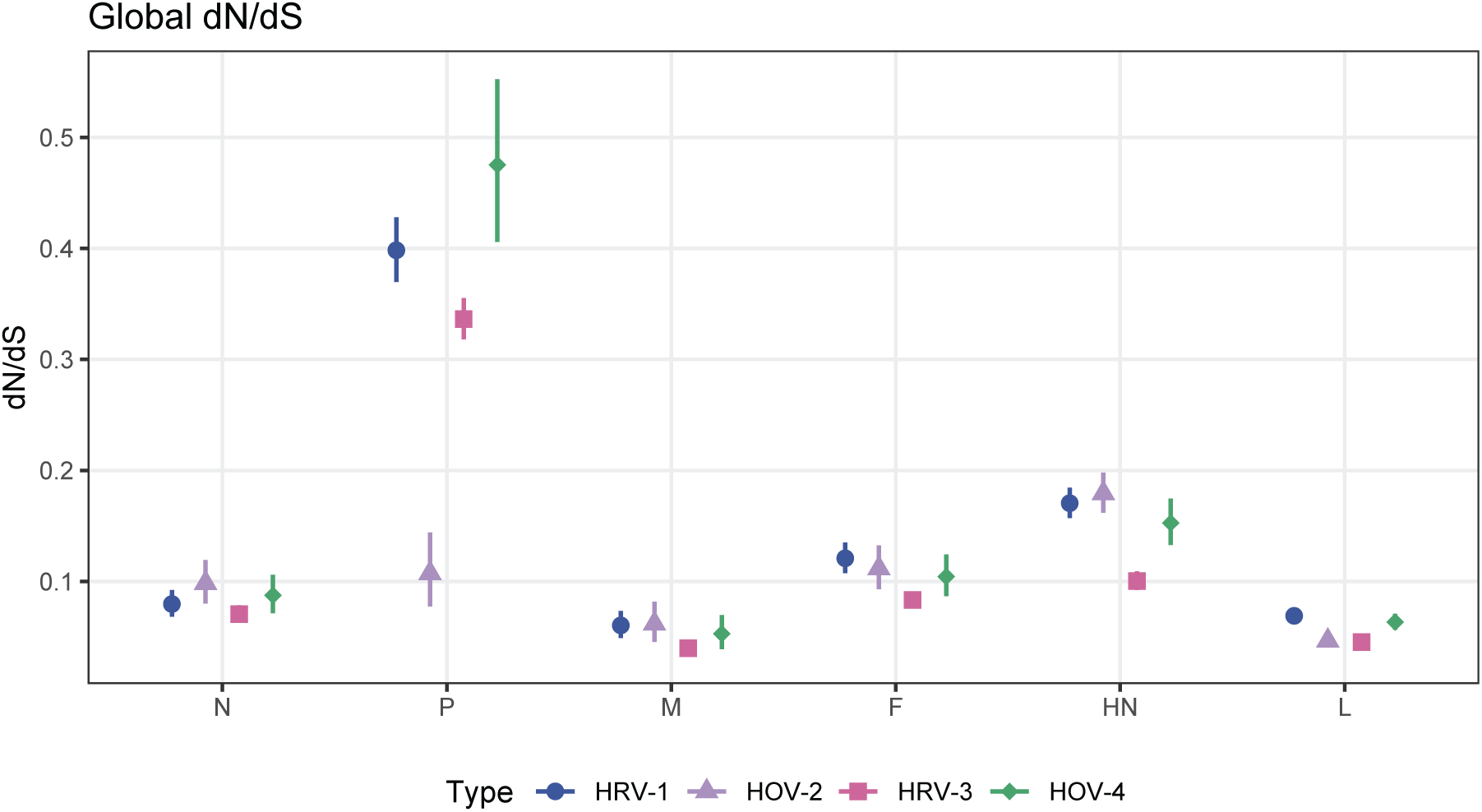
Comparative selection pressures across HPIV genes based on combined datasets across years. Global dN/dS estimates for six viral proteins, including nucleocapsid protein (N), phosphoprotein (P), matrix protein (M), fusion protein (F), hemagglutinin-neuraminidase glycoprotein (HN), and large polymerase protein (L). Global dN/dS ratios were estimated using the single-likelihood ancestor counting (SLAC) method. Means of dN/dS ratios are indicated by distinct symbols, with vertical bars representing 95% confidence intervals.

**Appendix Figure S27.**
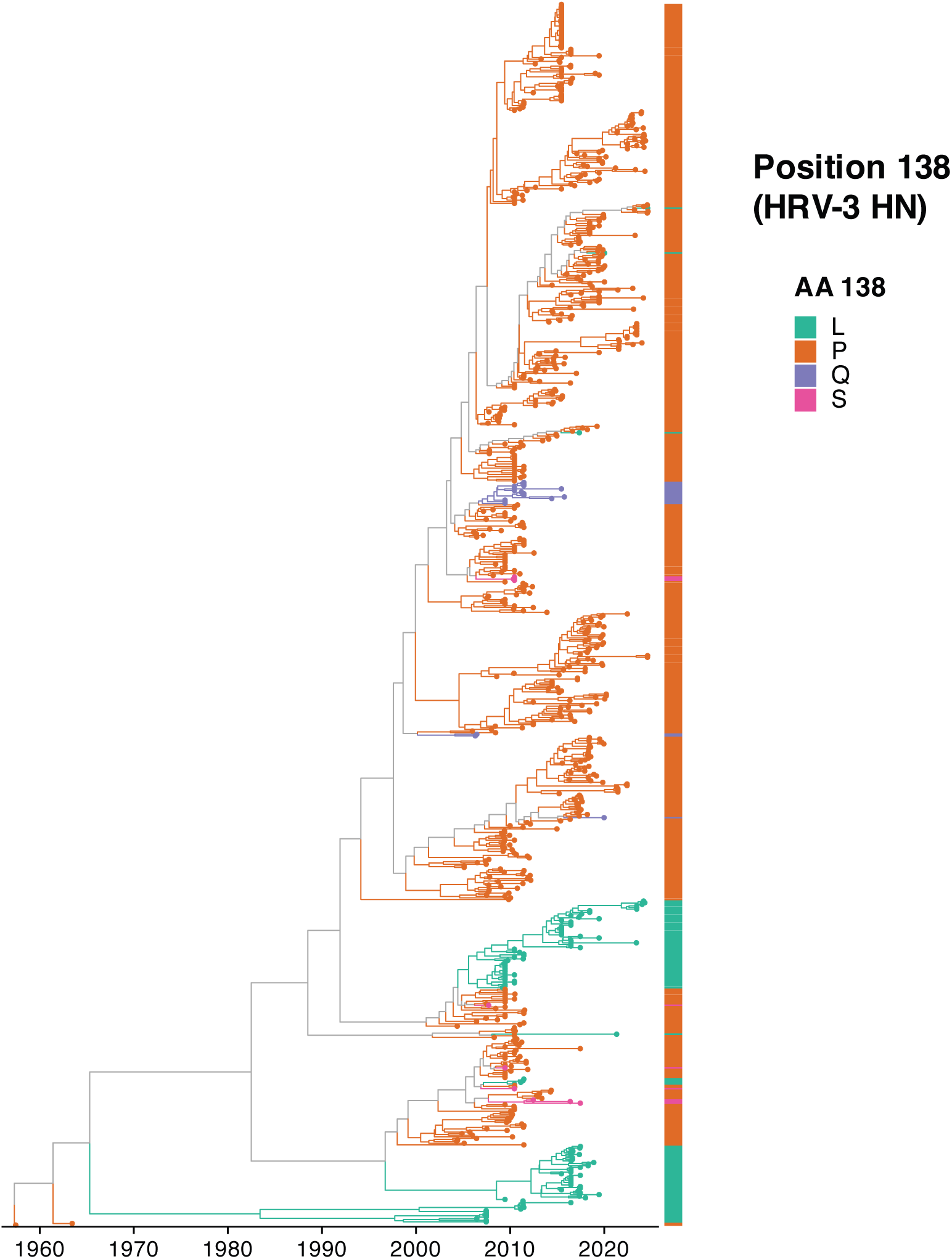
Mapping of positively selected amino acid residue 138 of the HN protein onto the HRV-3 HN phylogeny. Coloured branches indicate different amino acid residues present at position 138, highlighting the distribution of different amino acid substitutions across the phylogeny.

**Appendix Figure S28.**
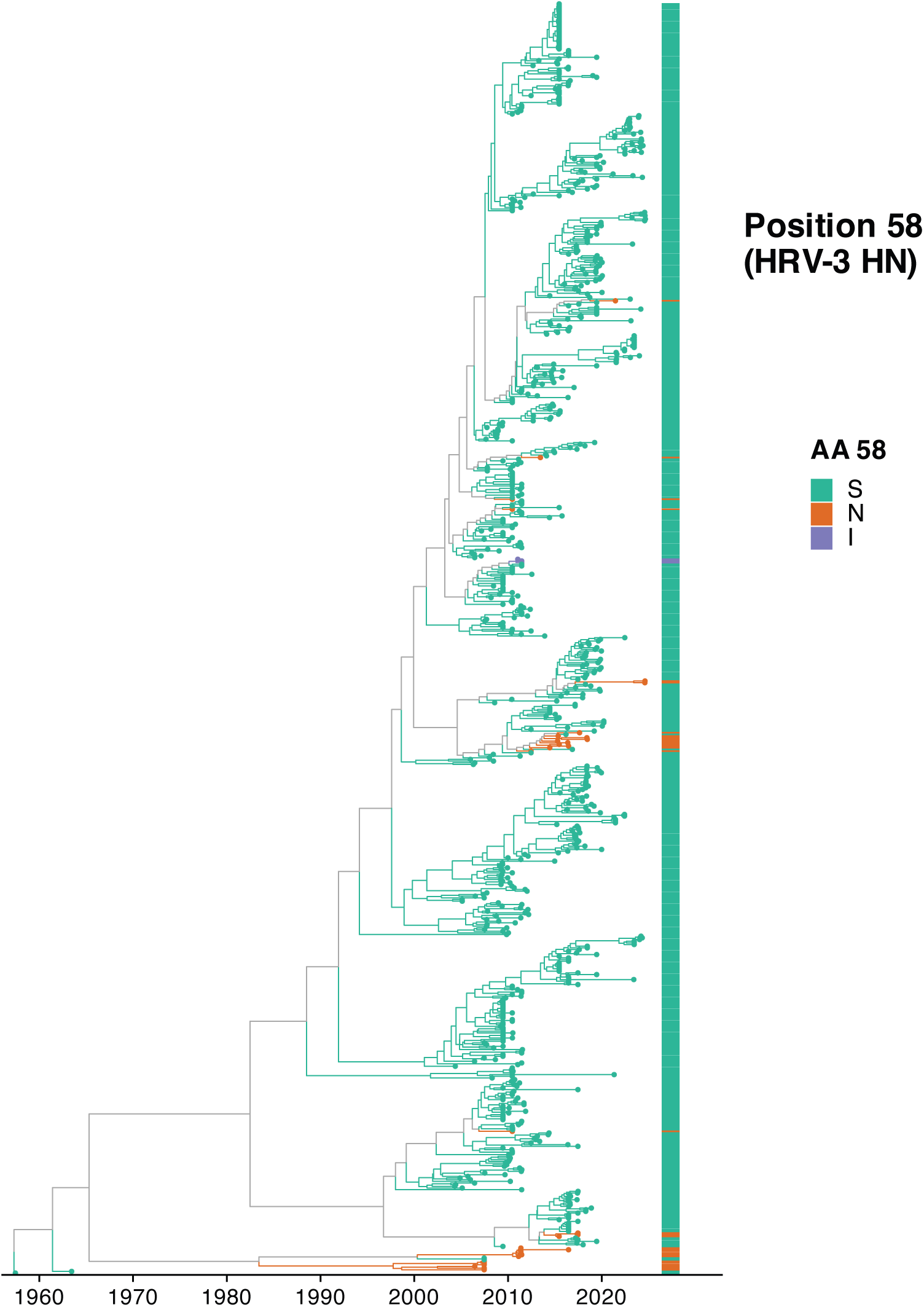
Mapping of positively selected amino acid residue 58 of the HN protein onto the HRV-3 HN phylogeny. Coloured branches indicate different amino acid residues present at position 58, highlighting the distribution of different amino acid substitutions across the phylogeny.

**Appendix Figure S29.**
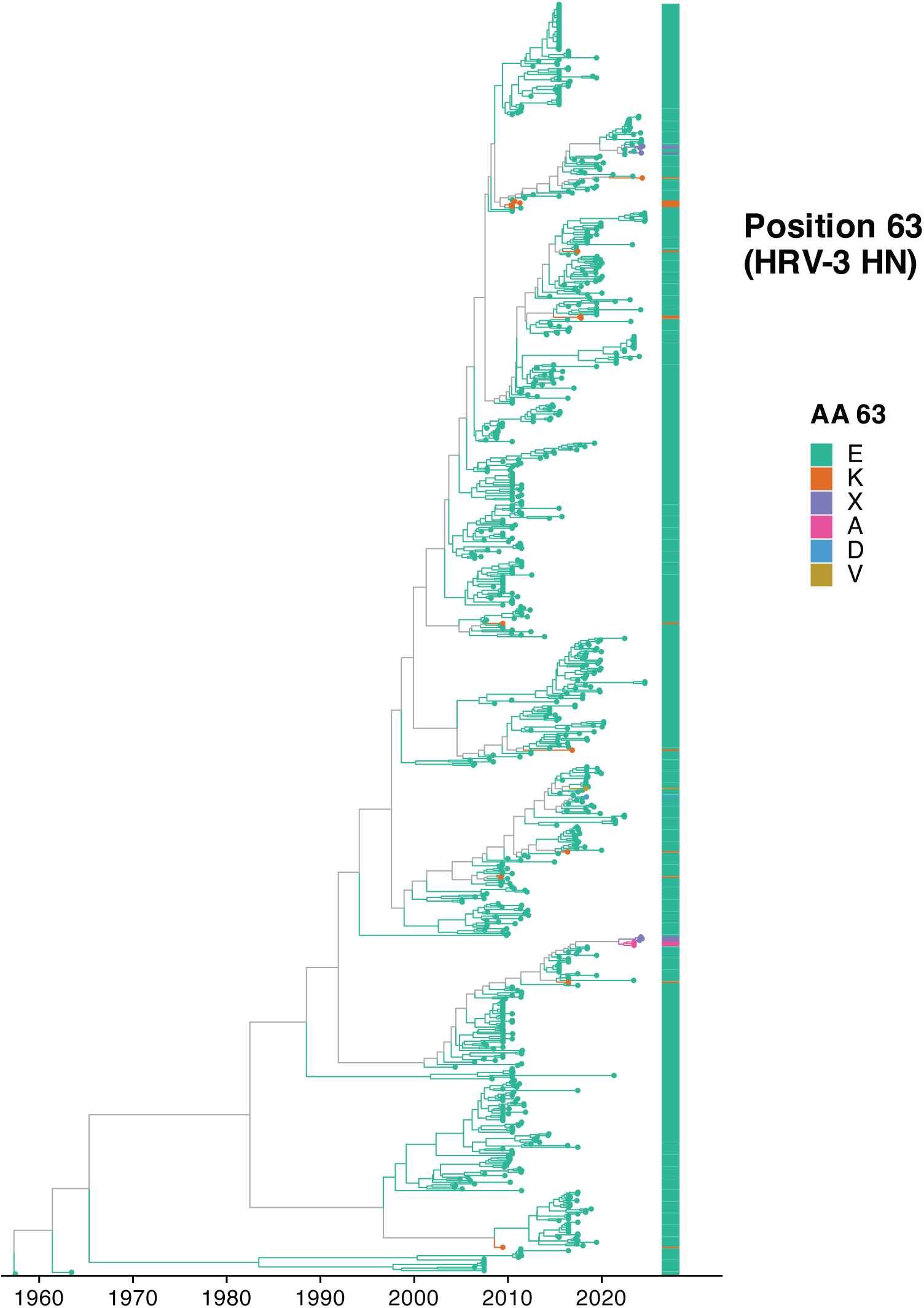
Mapping of positively selected amino acid residue 63 of the HN protein onto the HRV-3 HN phylogeny. Coloured branches indicate different amino acid residues present at position 63, highlighting the distribution of different amino acid substitutions across the phylogeny. X denotes missing amino acid residue at this position.

**Appendix Figure S30.**
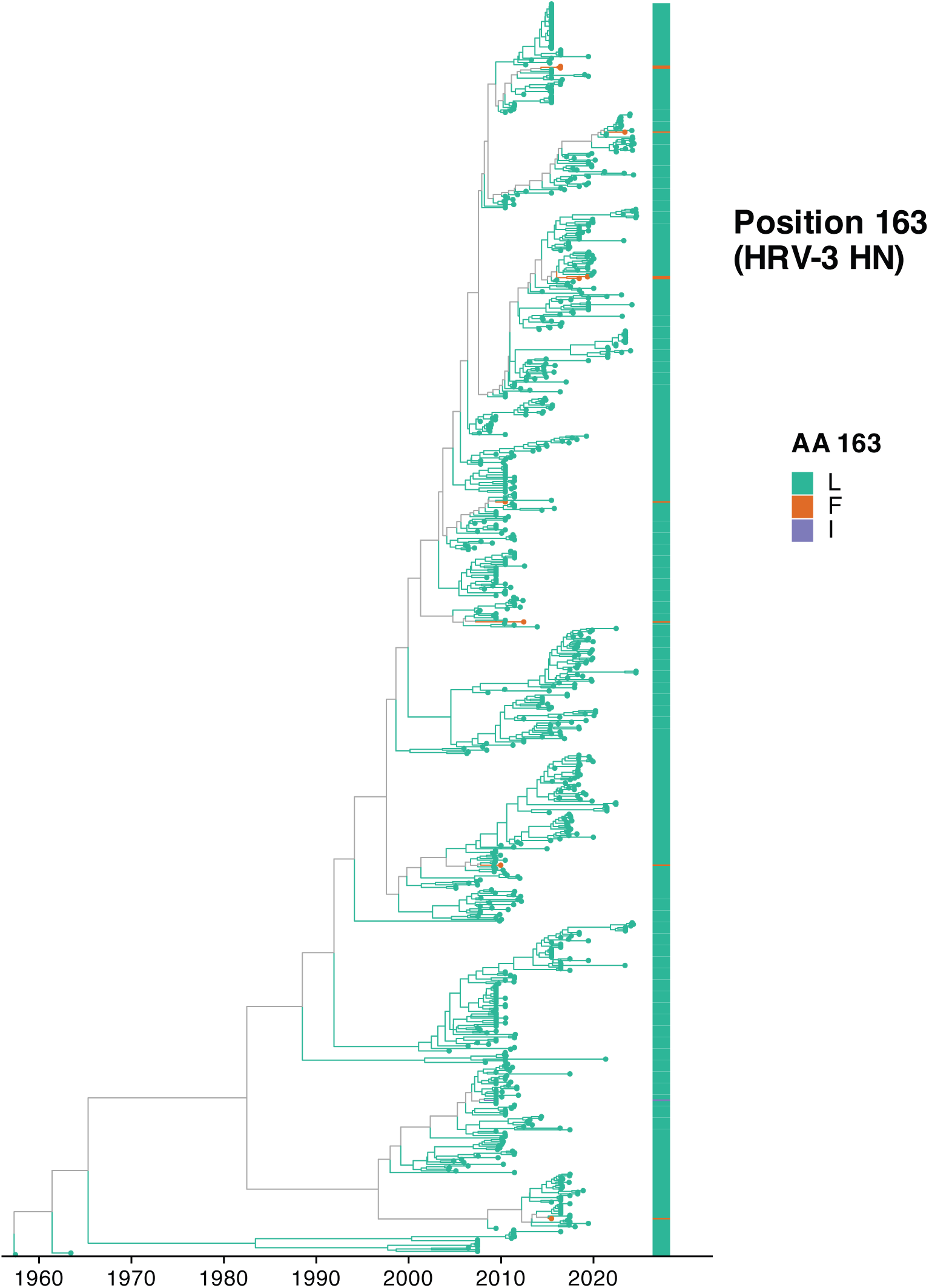
Mapping of positively selected amino acid residue 163 of the HN protein onto the HRV-3 HN phylogeny. Coloured branches indicate different amino acid residues present at position 163, highlighting the distribution of different amino acid substitutions across the phylogeny.

**Appendix Figure S31.**
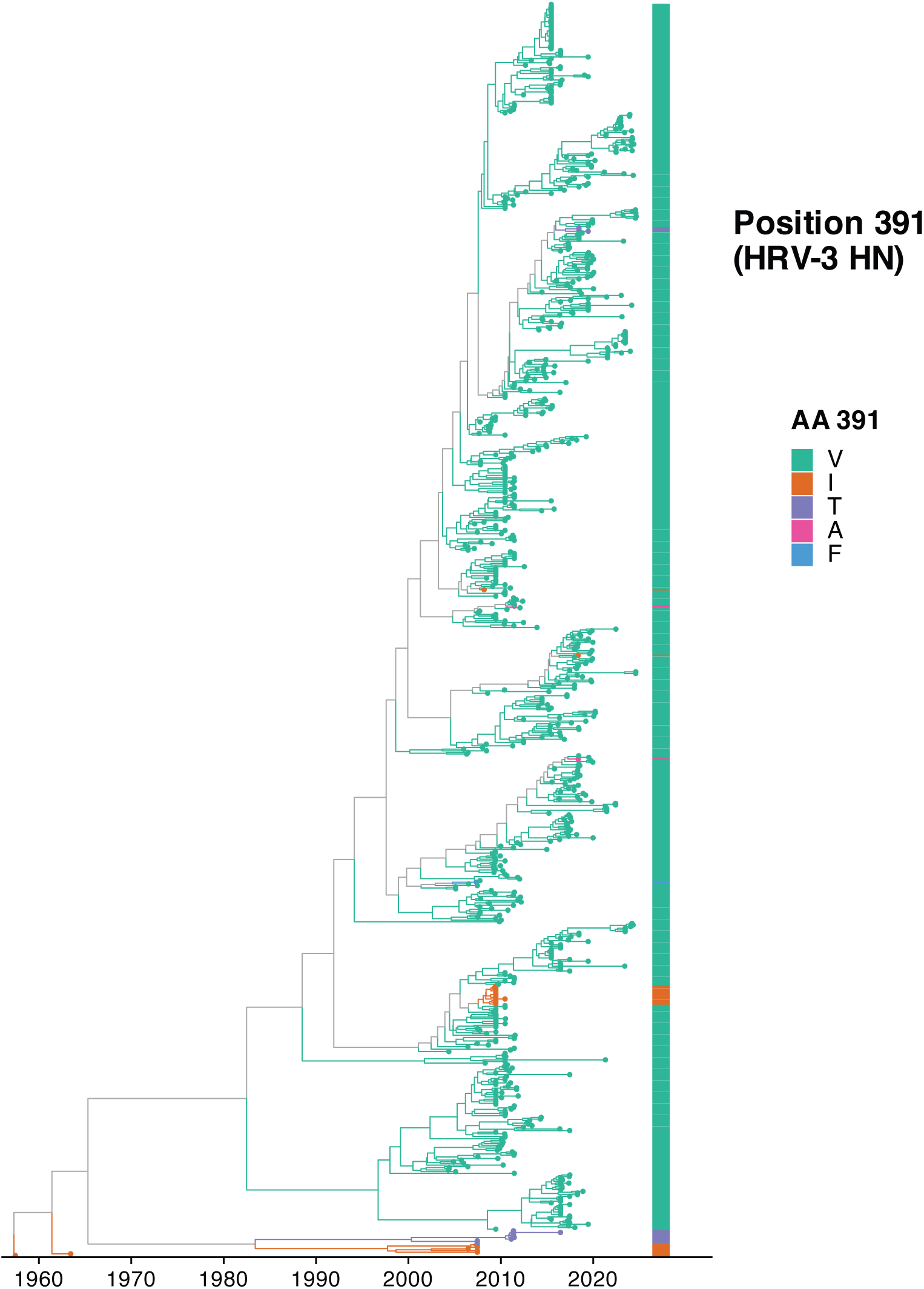
Mapping of positively selected amino acid residue 391 of the HN protein onto the HRV-3 HN phylogeny. Coloured branches indicate different amino acid residues present at position 391, highlighting the distribution of different amino acid substitutions across the phylogeny.

**Appendix Figure S32.**
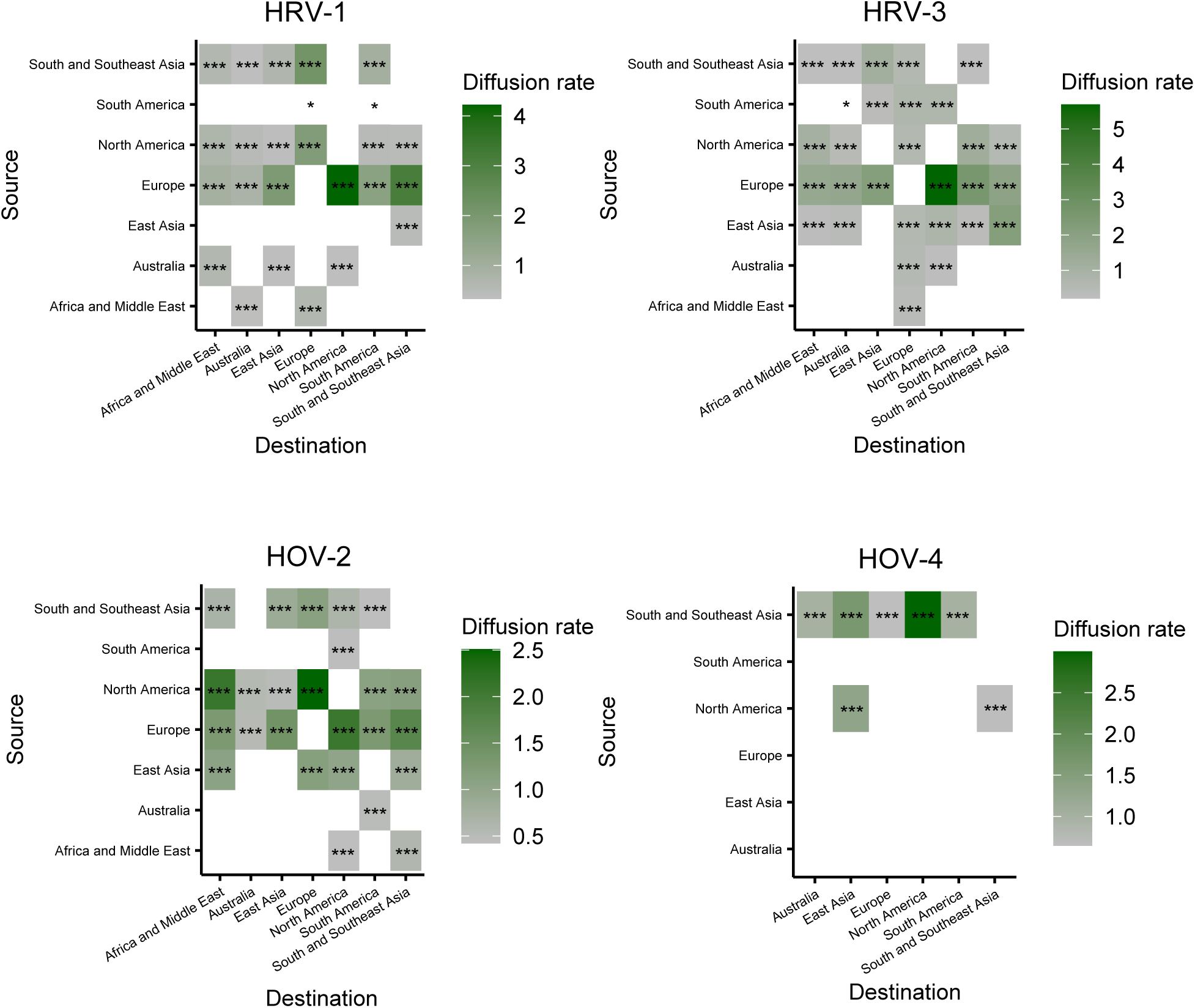
Comparative migration patterns among the four HPIV types. Heatmaps showing the mean migration rates between pairwise geographic locations. Only migration routes supported by a Bayes factor greater than 3 are shown. The degree of rate support is as follows: BF ≥ 100 indicates decisive (***), 10 ≤ BF < 100 indicates strong support (**) and 3 ≤ BF < 10 indicates supported (*).

**Appendix Figure S33.**
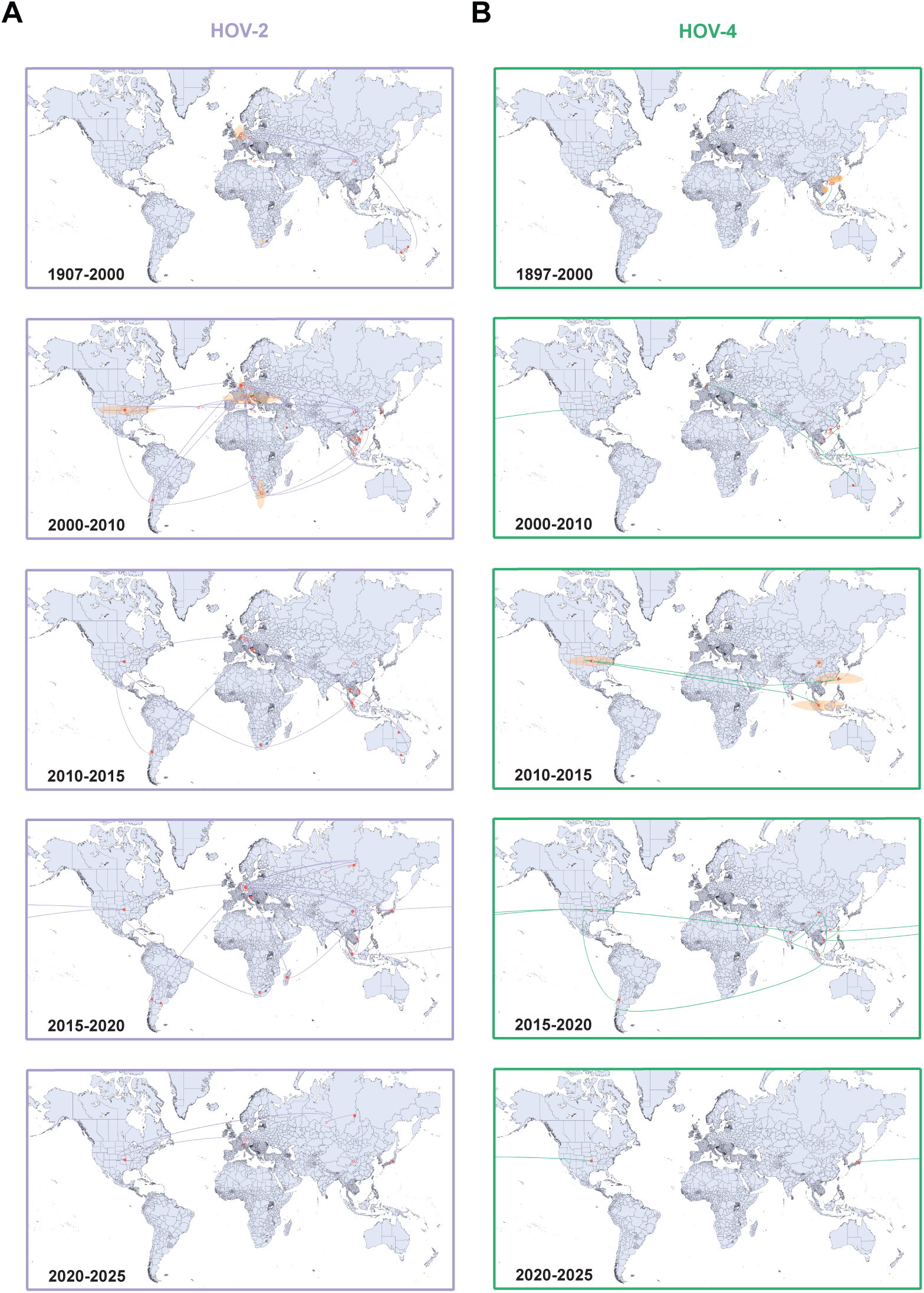
Continuous phylogeography of human orthorubulavirus. Spatiotemporal diffusion of (*A*) HOV-2 and (*B*) HOV-4 across five time periods. Spatial diffusion was modelled in continuous geographic space using latitude and longitude as bivariate traits under a Cauchy relaxed random walk (RRW) model.

